# Multi-omic analysis of a hyper-diverse plant metabolic pathway reveals evolutionary routes to biological innovation

**DOI:** 10.1101/136937

**Authors:** Gaurav D. Moghe, Bryan J. Leong, Steven Hurney, A. Daniel Jones, Robert L. Last

## Abstract

The diversity of life on Earth is a result of continual innovations in molecular networks influencing morphology and physiology. Plant specialized metabolism produces hundreds of thousands of compounds, offering striking examples of these innovations. To understand how this novelty is generated, we investigated the evolution of the Solanaceae family-specific, trichome-localized acylsugar biosynthetic pathway using a combination of mass spectrometry, RNA-seq, enzyme assays, RNAi and phylogenetics in non-model species. Our results reveal that hundreds of acylsugars are produced across the Solanaceae family and even within a single plant, revealing this phenotype to be hyper-diverse. The relatively short biosynthetic pathway experienced repeated cycles of innovation over the last 100 million years that include gene duplication and divergence, gene loss, evolution of substrate preference and promiscuity. This study provides mechanistic insights into the emergence of plant chemical novelty, and offers a template for investigating the ∼300,000 non-model plant species that remain underexplored.

## Impact statement

Integrative analyses of a specialized metabolic pathway across multiple non-model species revealed mechanisms of emergence of chemical novelty in plant metabolism.

## Introduction

Since the first proto-life forms arose on our planet some four billion years ago, the forces of mutation, selection and drift have generated a world of rich biological complexity. This complexity, evident at all levels of biological organization, has intrigued humans for millennia (Tipton, 2008; Mayr, 1985). Plant metabolism, estimated to produce hundreds of thousands of products with diverse structures across the plant kingdom (Fiehn, 2002; Afendi et al., 2012), provides striking examples of this complexity. Plant metabolism is traditionally divided into primary and secondary/specialized, the former referring to production of compounds essential for plant development and the latter encompassing metabolites documented as important for plant survival in nature and metabolites of as yet unknown functional significance (Pichersky and Lewinsohn, 2011; Moghe and Last, 2015). While primary metabolism generally consists of highly conserved pathways and enzymes, specialized metabolic pathways are in a state of continuous innovation (Milo and Last, 2012). This dynamism produced numerous lineage-specific metabolite classes such as steroidal glycoalkaloids in Solanaceae (Wink, 2003), benzoxazinoid alkaloids in Poaceae (Dutartre et al., 2012), betalains in Caryophyllales (Brockington et al., 2015) and glucosinolates in Brassicales (Halkier and Gershenzon, 2006). The structural diversity produced by lineage-specific pathways makes them exemplary systems for understanding the evolution of novelty in the living world.

Previous studies investigating the emergence of lineage-specific metabolite classes uncovered the central role of gene duplication and diversification in this process: e.g. duplications of isopropylmalate synthase (glucosinolates and acylsugars), cycloartenol synthase (avenacin), tryptophan synthase (benzoxazinoids) and anthranilate synthase (acridone alkaloids) (Moghe and Last, 2015). Duplications of members of enzyme families (e.g. cytochromes P450, glycosyltransferases, methyltransferases, BAHD acyltransferases) also play major roles in generating chemical novelty, with biosynthesis of >40,000 structurally diverse terpenoids — produced partly due to genomic clustering of terpene synthases and cytochrome P450 enzymes (Boutanaev et al., 2015) — as an extreme example. These duplicate genes, if retained, may experience sub- or neo-functionalization via transcriptional divergence and evolution of protein-protein interactions as well as via changes in substrate preference, reaction mechanism and allosteric regulation to produce chemical novelty (Moghe and Last, 2015). In this study, we sought to understand the molecular processes by which a novel class of metabolites — acylsugars — emerged in the Solanaceae family.

Acylsugars are lineage-specific plant specialized metabolites detected in multiple genera of the Solanaceae family including *Solanum* (King et al., 1990; Schilmiller et al., 2010; Ghosh et al., 2014), *Petunia* (Kroumova and Wagner, 2003), *Datura* (Forkner and Hare, 2000) and *Nicotiana* (Kroumova and Wagner, 2003; Kroumova et al., 2016). These compounds, produced in the tip cell of trichomes on leaf and stem surfaces (Schilmiller et al., 2012, 2015; Ning et al., 2015; Fan et al., 2016a), typically consist of a sucrose or glucose core esterified to groups derived from fatty acid or branched chain amino acid metabolism (Figure 1A). Multiple studies demonstrated that acylsugars mediate plant-insect and plant-fungus interactions (Puterka et al., 2003; Simmons et al., 2004; Weinhold and Baldwin, 2011; Leckie et al., 2016; Luu et al., 2017), with their structural diversity documented as functionally important in some cases (Puterka et al., 2003; Leckie et al., 2016).

**Figure 1:**
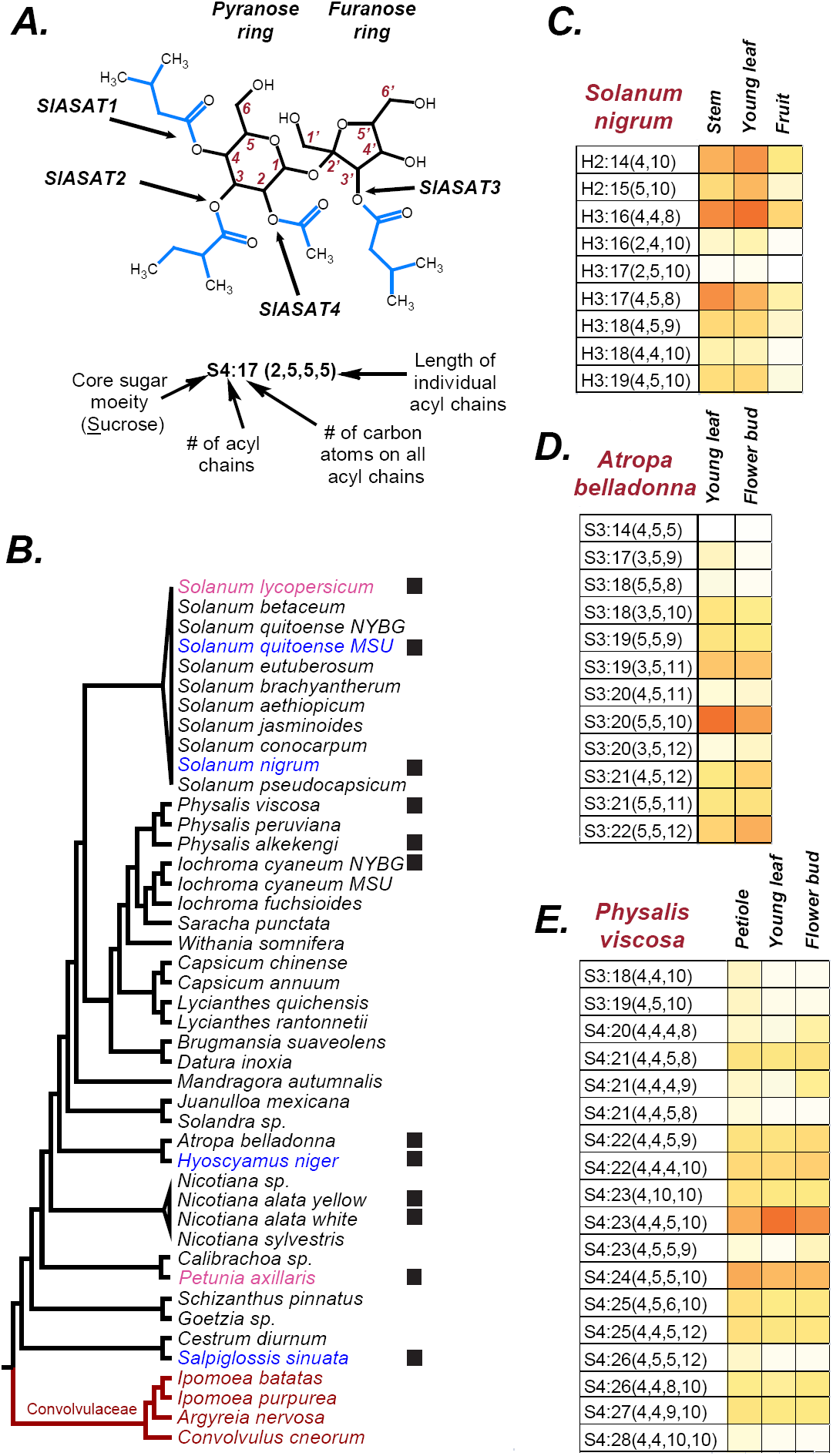
Acylsugars in Solanaceae. (A) An example acylsugar from tomato. The nomenclature of acylsugars and the ASAT enzymes responsible for acylation of specific positions are described. Carbon numbering is shown in red. Sl refers to *S. lycopersicum*. The phylogenetic position of the ASAT enzymes is shown in Figure Supplement 1. (B) Species sampled for acylsugar extractions. Phylogeny is based on the maximum likelihood tree of 1075 species (Särkinen et al, 2013). Species with black squares show presence of acylsugars in mass spectrometry. Species highlighted in blue were cultivated for RNA-seq. Species in red are Convolvulaceae species. Species in pink were not sampled in this study but have been extensively studied in the context of acylsugar biosynthesis (see main text). More information about these species sampled at the NYBG is provided in File Supplement 1 and Source Data 1. (C,D and E): Individual acylsugars from three representative species. Color scale ranges from no acylsugar (white) to maximum relative intensity in that species (orange). Peak areas of isomeric acylsugars were combined. *nigrum* produced acylsugars consistent with a hexose (H) core. Acylsugars identified from other species are described in Figure Supplement 2 and 3, and the raw peak intensity values obtained from different species are provided in Source Data 2. Figure Supplement 4 shows fragmentation patterns of select acylsugars under positive ionization mode.

Acylsugars, compared to other specialized metabolic classes such as alkaloids, phenylpropanoids and glucosinolates, are biosynthetically rather simple. These compounds are produced by a set of enzymes called acylsugar acyltransferases (ASATs), which catalyze sequential addition of acyl chains to the sucrose molecule (Figure 1A) (Schilmiller et al., 2012, 2015; Fan et al., 2016a). ASATs are members of Clade III of the large and functionally diverse BAHD enzyme family ( St Pierre and De Luca, 2000; D’Auria, 2006) (Figure 1-Figure Supplement 1). Despite their evolutionary relatedness, *S. lycopersicum* ASATs (SlASATs) are only ∼40% identical to each other at the protein level. ASATs exhibit different activities across wild tomatoes due to ortholog divergence, gene duplication and neo-functionalization, leading to divergence in acceptor and donor substrate repertoire of ASATs between wild tomato species (Schilmiller et al., 2015; Ning et al., 2015; Fan et al., 2016a). In addition, loss of an ASAT enzyme also contributed to diversification of acylsugar phenotypes between individuals of the same species (Kim et al., 2012). This diversity, primarily studied in closely related wild tomato species and underexplored in the broader Solanaceae family, creates opportunities for further exploration of the mechanisms generating biological complexity and variation.

Previous studies of acylsugar biosynthesis primarily focused on closely related *Solanum* species. In this study, we sought to understand the timeline for emergence of the ASAT activities and explore the evolution of the acylsugar biosynthetic pathway since the origin of the Solanaceae. Typically, significant hits obtained using BLAST searches are analyzed in a phylogenetic context to understand enzyme origins (Frey et al., 1997; Ober and Hartmann, 2000; Qi et al., 2004; Benderoth et al., 2006). However, ASAT orthologs can experience functional diversification due to single amino acid changes and/or duplication (Schilmiller et al., 2015; Fan et al., 2016a), precluding functional assignment based on sequence similarity. Thus, we inferred the origins and evolution of the pathway with a bottom-up approach; starting by assessing the diversity of acylsugar phenotypes across the family using mass spectrometry. Our findings illustrate the varied mechanisms by which this specialized metabolic pathway evolved and have implications for the study of chemical novelty in the plant kingdom.

## Results and Discussion

### Diversity of acylsugar profiles across the Solanaceae

While the Solanaceae family comprises 98 genera and >2700 species (Olmstead and Bohs, 2007), there are extensive descriptions of acylsugar diversity reported for only a handful of species (Severson et al., 1985; King et al., 1990; Shinozaki et al., 1991; Ghosh et al., 2014). In this study, we sampled vegetative tissue surface metabolites from single plants of 35 Solanaceae and four Convolvulaceae species. These species were sampled at the New York Botanical Gardens and Michigan State University (Figure 1B; Figure 1-File Supplement 1A), and acylsugar profiles were obtained using liquid chromatography-mass spectrometry (LC/MS) with collision-induced dissociation (CID; see Methods) (Schilmiller et al., 2010; Ghosh et al., 2014; Fan et al., 2016b). Molecular and substructure (fragment) masses obtained by LC/MS-CID were used to annotate acylsugars in *Solanum nigrum*, *Solanum quitoense*, *Physalis alkekengi*, *Physalis viscosa, Iochroma cyaneum*, *Atropa belladonna*, *Nicotiana alata*, *Hyoscyamus niger* and *Salpiglossis sinuata* (Salpiglossis) (Figure 1B,C; Figure 1-Figure Supplement 2; **Figure 1-Source Data 1**). Plant extracts without detectable acylsugars generally lacked glandular trichomes (Fisher Exact Test p=2.3e-6) (Figure 1-File Supplement 1B). In addition, the acylsugar phenotype is quite dynamic and can be affected by factors such as developmental stage, environmental conditions and the specific accession sampled (Kim et al., 2012; Ning et al., 2015; Schilmiller et al., 2015). These factors may also influence the detection of acylsugars in some species.

The suite of detected acylsugars exhibited substantial diversity, both in molecular and fragment ion masses. Most species accumulated acylsugars with mass spectra consistent with disaccharide cores — most likely sucrose — esterified with short- to medium-chain aliphatic acyl groups, similar to previously characterized acylsugars in cultivated and wild tomatoes (Schilmiller et al., 2010; Ghosh et al., 2014). However, *S. nigrum* acylsugar data suggested exclusive accumulation of acylhexoses (Figure 1C), with fragmentation patterns similar to previously analyzed *S. pennellii* acylglucoses (Schilmiller et al., 2012). Mass spectra of *S. quitoense* acylsugars also revealed acylsugars with features distinct from any known acylsugars (Figure 1-Figure Supplement 2), and structures of these will be described in detail in a separate report. In total, more than 100 acylsucroses and at least 20 acylsugars of other forms were annotated with number and lengths of acyl groups based on pseudomolecular and fragment ion masses (Figure 1-Figure Supplement 2). Several hundred additional low-abundance isomers and novel acylsugars were also detected (Figure 1-Figure Supplement 3). For example, in Salpiglossis alone, >400 chromatographic peaks had *m/z* ratios and mass defects consistent with acylsugars **(For review only - Additional File 1)**. This diversity is notable when compared to ∼33 detected acylsucroses in cultivated tomato (Ghosh et al., 2014).

Mass spectra also revealed substantial diversity in the number and lengths of acyl chains (Figure 1C,D,E, Figure 1-Figure Supplement 2). Based on negative ion mode data, the number of acyl chains on the sugar cores ranged from two to six (Figure 1-Figure Supplement 2), with chain lengths from 2-12 carbons. Across the Solanaceae, we found species that incorporate at least one common aliphatic acyl chain in all their major acylsugars: e.g. chains of length C5 in *A. belladonna*, C4 in *P. viscosa*, and C8 in *P. alkekengi* (Figure 1D,E; Figure 1-Figure Supplement 2). Longer C10 and C12 chain-containing acylsugars were found in multiple species (*P. viscosa*, *A. belladonna*, *H. niger*, *S. nigrum* and *S. quitoense*) (Figure 1C-E, Figure 1-Figure Supplement 2). Mass spectra consistent with acylsugars containing novel acyl chains were also detected. While we could not differentiate between acyl chain isomers (e.g. *iso*-C5 [iC5] vs. *anteiso*-C5 [aiC5]) based on CID fragmentation patterns, our data reveal large acyl chain diversity in acylsugars across the Solanaceae. Such diversity can result from differences in the intracellular concentrations of acyl CoA pools or divergent substrate specificities of individual ASATs. A previous study from our lab identified allelic variation in the enzyme isopropylmalatesynthase3, which contributes to differences in abundances of iC5 or iC4 chains in acylsugars of *S. lycopersicum* and some *S. pennellii* accessions (Ning et al., 2015). *ASAT* gene duplication (Schilmiller et al., 2015), gene loss (Kim et al., 2012) and single residue changes (Fan et al., 2016a) also influence chain diversity in acylsugars, illustrating the various ways by which acylsugar phenotypes may be generated in the Solanaceae.

Published data from *S. pennellii* and *S. habrochaites* revealed differences in furanose ring acylation on acylsucroses (Schilmiller et al., 2015). Ring-specific acylation patterns can be evaluated using positive mode mass spectrometry and CID, which generates fragment ion masses from cleavage of the glycosidic linkage. We found furanose ring acylation in almost all tested species in Solanaceae (Figure 1-Figure Supplement 4A-E); however, the lengths of acyl chains on the ring varied. All tri-acylsucroses analyzed using positive mode CID data bore all acyl chains on one ring, likely the pyranose ring — as evidenced by neutral loss of 197 Da (hexose plus NH_3_) from the [M+NH_4_]^+^ ion — unlike *S. lycopersicum,* which bears one acyl chain on the furanose ring. However, the substituents varied among tetra- and penta-acylsugars of different species, with some (e.g.: *N. alata*), showing up to four chains on the same ring (Figure 1-Figure Supplement 4A), and others (e.g.: *H. niger*, Salpiglossis) revealing spectra consistent with acylation on both pyranose and furanose rings (Figure 1-Figure Supplement 4B,C).

These results demonstrate the substantial acylsugar diversity across the Solanaceae family. To identify the enzymes that contribute to this diversity, we performed RNA-seq in four phylogenetically-spaced species with interesting acylsugar profiles, namely *S. nigrum*, *S. quitoense*, *H. niger* and Salpiglossis.

### Transcriptomic profiling of trichomes from multiple Solanaceae species

Our previous studies in cultivated tomato (Schilmiller et al., 2012, 2015; Ning et al., 2015; Fan et al., 2016a) demonstrated that identifying genes with expression enriched in stem/petiole trichomes compared to shaved stem/petiole without trichomes is a productive way to find acylsugar biosynthetic enzymes. We sampled polyA RNA from these tissues from four species and performed *de novo* read assembly (Table 1). These assemblies were used to find transcripts preferentially expressed in the trichomes (referred to as “trichome-high transcripts”) and to develop hypotheses regarding their functions based on homology.

**Table 1:** RNA-seq data statistics.

| **Item** | ***S. nigrum*** | ***S. quitoense*** | ***H. niger*** | ***S. sinuata*** |
| --- | --- | --- | --- | --- |
| **Original read pairs** | 81,314,841 | 85,374,110 | 86,161,659 | 80,302,734 |
| **Filtered read pairs**<br>**(% original)** | 73346531<br>(90.2%) | 76734781<br>(89.9%) | 76819022<br>(89.2%) | 71129160<br>(88.6%) |
| **Normalized read**<br>**pairs**<br>**(% normalized)** | 17301238<br>(23.6%) | 15350023<br>(20.0%) | 20779972<br>(27.1%) | 17905057<br>(25.2%) |
| **Total transcript**<br>**isoforms** | 160,583 | 124,958 | 189,711 | 149,136 |
| **Longest isoforms**<br>**(% total)** | 78,020<br>(49%) | 72,426<br>(58%) | 96,379<br>(51%) | 77,970<br>(52.3%) |
| **With >10 reads** | 32,105 | 32,044 | 38,252 | 32,798 |
| **With predicted**<br>**peptide>50aa** | 23,224 | 22,289 | 26,262 | 23,570 |
| **Differentially**<br>**expressed<sup>1,2</sup>**<br>**(% >10 reads)** | 10,386 (32%) | 12,194 (38%) | 9007 (24%) | 7091 (22%) |
| **Trichome high<sup>2</sup>**<br>**(% differentially**<br>**expressed)** | 2292 (22.1%) | 3547 (29.1%) | 3321 (36.8%) | 1888 (26.6%) |

<sup>1</sup> Differentially expressed genes at $p < 0.05$ (corrected for multiple testing); <sup>2</sup> $p < 0.05$ , fold change $> 2$
<sup>2</sup> Some differentially expressed transcripts were confirmed by RT-PCR, as shown in Figure Supplement 1.

Overall, 1888-3547 trichome-high transcripts (22-37% of all differentially expressed transcripts) were identified across all four species (False Discovery Rate adjusted p<0.05, fold change≥2) (Table 1). These transcripts were subjected to a detailed analysis including coding sequence prediction, binning into 25,838 orthologous groups, assignment of putative functions based on tomato gene annotation and Gene Ontology enrichment analysis (see Methods, Table 1). Analysis of the enriched categories (Fisher exact test corrected p<0.05) revealed that only 20 of 70 well-supported categories (≥10 transcripts) were enriched in at least three species **(Table 1-File Supplement 1)**, suggesting existence of diverse transcriptional programs in the trichomes at the time of their sampling. Almost all enriched categories were related to metabolism, protein modification or transport, with metabolism-related categories being dominant **(Table 1-File Supplement 1)**. These results support the notion of trichomes as “chemical factories” (Schilmiller et al., 2008) and point to the metabolic diversity that might exist in trichomes across the Solanaceae.

A major goal of this study was to define the organization of the acylsugar biosynthetic pathway at the origin of the Solanaceae, prompting us to focus on Salpiglossis, whose phylogenetic position is of special interest in inferring the ancestral state of the biosynthetic pathway. We first validated the plant under study as Salpiglossis using a phylogeny based on *ndhF* and *trnLF* sequences (Figure 2-Figure Supplement 1A,B). A previously published maximum likelihood tree of 1075 Solanaceae species suggested *Salpiglossis* as an extant species of the earliest diverging lineage in Solanaceae (Särkinen et al., 2013). However, some tree reconstruction approaches show *Duckeodendron* and *Schwenckia* as emerging from the earliest diverging lineages, and Salpiglossis and Petunioideae closely related to each other (Olmstead et al., 2008; Särkinen et al., 2013). Thus, our further interpretations are restricted to the last common ancestor of Salpiglossis-Petunia-Tomato (hereafter referred to as the Last Common Ancestor [LCA]) that existed ∼22-28 mya. To infer the ancestral state of the acylsugar biosynthetic pathway in the LCA, we characterized the pathway in Salpiglossis using *in vitro* and *in planta* approaches.

### *In vitro* investigation of Salpiglossis acylsugar biosynthesis

The acylsugar structural diversity and phylogenetic position of Salpiglossis led us to characterize the biosynthetic pathway of this species. NMR analysis of Salpiglossis acylsugars revealed acylation at the R2, R3, R4 positions on the pyranose ring and R1′, R3′, R6′ positions on the furanose ring (Figure 2A; Figure 2A-Table Supplement 1) (For review only – Additional File 1). The acylation positions are reminiscent of *Petunia axillaris* (Pa) acylsucroses where PaASAT1, PaASAT2, PaASAT3 and PaASAT4 acylate with aliphatic precursors at R2, R4, R3 and R6 on the six-carbon pyranose ring, respectively **(For review only – Additional File 2)**. Thus, we tested the hypothesis that PaASAT1,2,3 orthologs in Salpiglossis function as SsASAT1,2,3 respectively.

**Figure 2:**
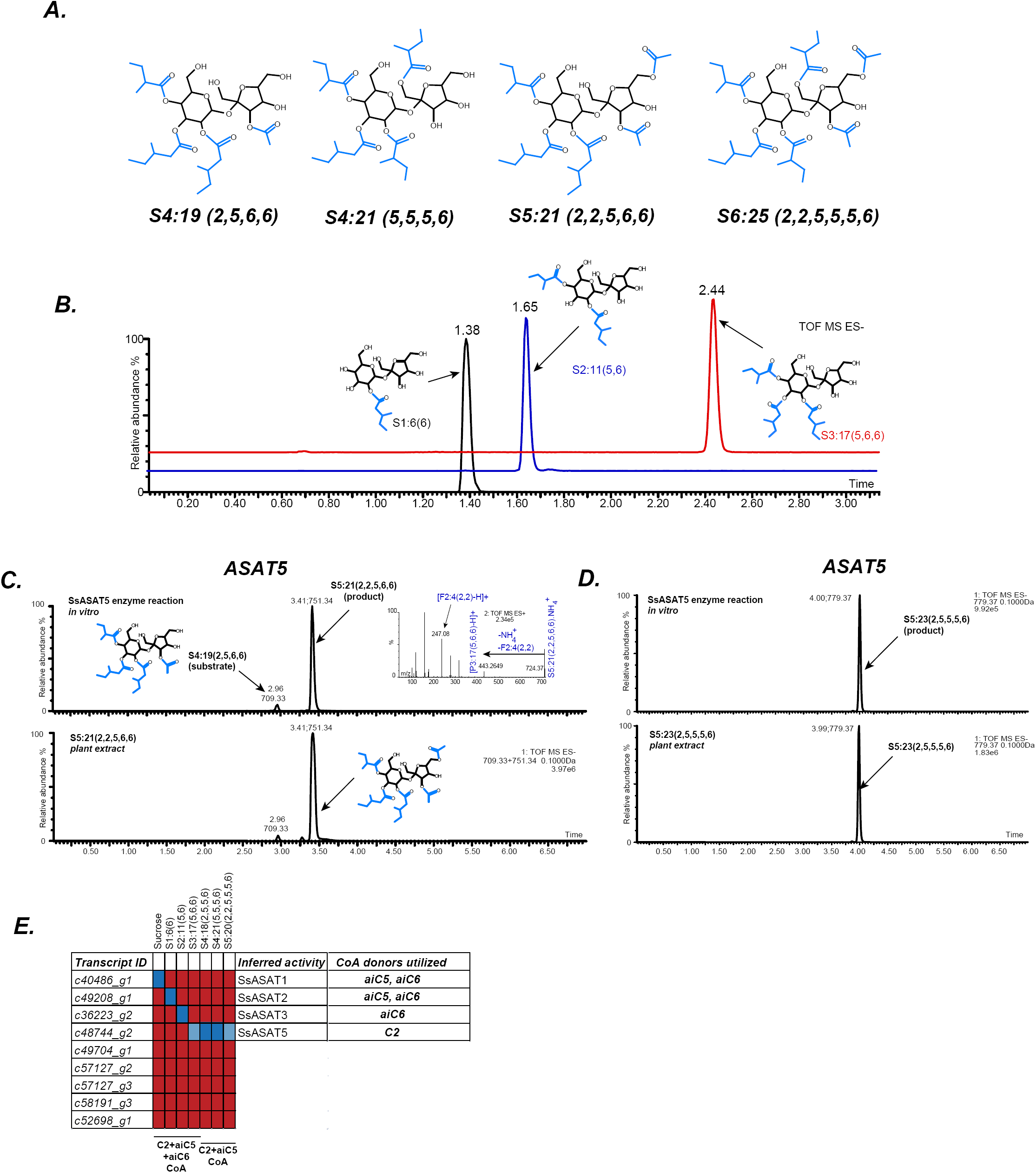
In vitro validation of Salpiglossis ASAT candidates. (A) NMR derived structures of three Salpiglossis acylsugars. NMR resonances used to interpret the first three structures are described in Table Supplement 1. We verified the plant under study as Salpiglossis using genetic markers. These results are shown in Figure Supplement 1A,B. (B) Results of enzyme assays for SsASAT1 (black), SsASAT2 (blue) and SsASAT3 (red). Numbers above the peaks represent the retention times of the individual compounds (see Methods), whose predicted structures are shown alongside. Additional validation of these *in vitro* results is described in Figure Supplements 2-5. (C,D) The SsASAT5 reactions, whose products have the same retention time as *in planta* compounds. Inset in panel C shows positive mode fragmentation and predicted acyl chains on pyranose [P] and furanose [F] rings. SsASAT5 also performs additional acylation activities as shown in Figure Supplement 6. (E) Testing various ASAT candidates with different acceptor (top) and donor (bottom) substates. Red indicates no activity seen by LC/MS, dark blue indicates a likely true activity, which results in a product usable by the next enzyme and/or a product that co-migrates with the most abundant expected compound. Light blue color indicates that the enzyme can acylate a given substrate, but the product cannot be used by the next enzyme or does not co-migrate with the most abundant expected compound. The relationships of the enzymes with each other are shown in Figure Supplement 1C.

Thirteen Salpiglossis trichome-high BAHD family members were found (Figure 2- Figure Supplement 1C), with nine expressed in *Escherichia coli* (Figure 2-Table Supplement 2; Figure 2-File Supplements 1,2). Activities of the purified enzymes were tested using sucrose or partially acylated sucroses (Fan et al., 2016a, 2016b) as acceptor substrates. Donor C2, aiC5 and aiC6 acyl CoA substrates were tested based on the common occurrence of these ester groups in a set of 16 Salpiglossis acylsucroses purified for NMR **(For review only – Additional File 1)**. Representative NMR structures that illustrate the SsASAT positional selectivity described in the results below are shown in Figure 2A. Four of the tested candidates catalyzed ASAT reactions (Figure 2B-E). In the following description, we name the enzymes based on their order of acylation in the Salpiglossis acylsugar biosynthetic pathway. A description is provided in **Figure 2-Figure Supplement 2** to assist in understanding the chromatograms.

*<u>S</u>alpiglossis <u>s</u>inuata* ASAT1 (SsASAT1) generated mono-acylsugars from sucrose using multiple acyl CoAs (Figure 2B, Figure 2-Figure Supplements 2,3), similar to the donor substrate diversity of SlASAT1 (Fan et al., 2016a) (Figure 2-Figure Supplement 2). We infer that SsASAT1 primarily acylates the R2 position on the pyranose ring. This is based on (a) the S1:6(6) negative mode CID fragmentation patterns (Figure 2-Figure Supplement 3A) and (b) comparisons of chromatographic migration of the mono-acylsucroses produced by SsASAT1, PaASAT1, which acylates sucrose at R2 and matches the major SsASAT1 product, and SlASAT1, which acylates sucrose at R4 (Figure 2-Figure Supplement 3A,B) (**For review only - Additional File 2)**. This activity is similar to *P. axillaris* PaASAT1 and unlike *S. lycopersicum* SlASAT1.

SsASAT2 was identified by testing the ability of each of the other eight cloned enzymes to acylate the mono-acylated product of SsASAT1 (S1:5 or S1:6) as acyl acceptor and using aiC5 CoA as acyl donor. SsASAT2 catalyzed the formation of di-acylated sugars, which co-eluted with the PaASAT2 product but not with the SlASAT2 product (Figure 2B; Figure 2-Figure Supplement 3C,D). Positive mode fragmentation suggested that the acylation occurs on the same ring as SsASAT1 acylation (Figure 2-Figure Supplement 3C). This enzyme failed to acylate sucrose (Figure 2-Figure Supplement 4), supporting its assignment as SsASAT2.

SsASAT3 (Figure 2B, red chromatogram) added aiC6 to the pyranose ring of the di-acylated sugar acceptor. This aiC6 acylation reaction is in concordance with the observed *in planta* acylation pattern at the R3 position — aiC6 is present at this position in the majority of *S. sinuata* acylsugars **(For review only – Additional File 1)**. Surprisingly, the enzyme did not use aiC5 CoA, despite the identification of S3:15(5,5,5) and likely its acylated derivates [S4:20(5,5,5,5), S4:17(2,5,5,5), S5:22(2,5,5,5,5) and S5:19(2,2,5,5,5)] from *S. sinuata* extracts **(For review only – Additional File 1)**. SsASAT3-dependent R3 position acylation was further confirmed by testing the tri-acylated product with PaASAT4, which acylates at the R6 position of the pyranose ring (Figure 2-Figure Supplement 5) (For review only – Additional File 2). The successful R6 acylation by PaASAT4 is consistent with the hypothesis that SsASAT3 acylates the R3 position. Taken together, these results suggest that the first three enzymes generate acylsugars with aiC5/aiC6 at the R2 position (SsASAT1), aiC6 at the R3 position (SsASAT3) and aiC5/aiC6 at the R4 position (SsASAT2).

We could not identify the SsASAT4 enzyme(s) that performs aiC5 and C2 acylations on the R1′ and R3′ positions of tri-acylsucroses, respectively. However, we identified another enzyme, which we designate SsASAT5, that showed three activities acetylating tri-, tetra- and penta-acylsucroses (see Methods). SsASAT5 can perform furanose ring acetylation on tri- (Figure 2-Figure Supplement 6A,B) and tetra-acylsucroses (Figure 2C,D), and both furanose as well as pyranose ring acetylation on penta-acylsucroses (Figure 2-Figure Supplement 6C-F). All of the products produced by SsASAT5 *in vitro* co-migrate with acylsugars found in plant extracts, suggesting the SsASAT5 acceptor promiscuity also occurs *in planta*. Our observation that SsASAT5 can perform pyranose ring acetylation — albeit weakly **(Figure 2-Figure Supplements 6D,F)** — is at odds with NMR-characterized structures of a set of 16 purified Salpiglossis acylsugars, which show all acetyl groups on the furanose ring. However, a previous study described one pyranose R6-acylated penta-acyl sugar S5:22(2,2,6,6,6) in Salpiglossis (Castillo et al., 1989), suggesting the presence of accession-specific variation in enzyme function. Despite showing SsASAT4-, SsASAT5- and SsASAT6-like activities, we designate this enzyme SsASAT5 because its products have both acylation patterns and co-migration characteristics consistent with the most abundant penta-acylsugars from the plant (Figure 2C,D).

Overall, *in vitro* analysis revealed four enzymes that could catalyze ASAT reactions and produce compounds also detected in plant extracts (Figure 2E). We further verified that these enzymes are involved in acylsugar biosynthesis by testing the effects of perturbing their transcript levels using Virus Induced Gene Silencing (VIGS).

### *In planta* validation of acylsugar biosynthetic enzymes

To test the role of the *in vitro* identified ASATs *in planta*, we adapted a previously described tobacco rattle virus-based VIGS procedure (Dong et al., 2007; Velásquez et al., 2009) for Salpiglossis. We designed ∼300 bp long gene-specific silencing constructs for transient silencing of SsASAT1, SsASAT2, SsASAT3 and SsASAT5 **(Figure 2-File Supplements 1,2**), choosing regions predicted to have a low chance of reducing expression of non-target genes (see Methods). The Salpiglossis ortholog of the tomato phytoene desaturase (PDS) carotenoid biosynthetic enzyme was used as positive control (Figure 3A), with transcript level decreases confirmed for each candidate using qRT-PCR in one of the VIGS replicates (Figure 3B). As no standard growth or VIGS protocol was available for Salpiglossis, we tested a variety of conditions for agroinfiltration and plant growth, and generated at least two biological replicate experiments for each construct, run at different times **(Figure 3-Table Supplement 1)**. ASAT knockdown phenotypes were consistent, regardless of variation in environmental conditions.

**Figure 3:**
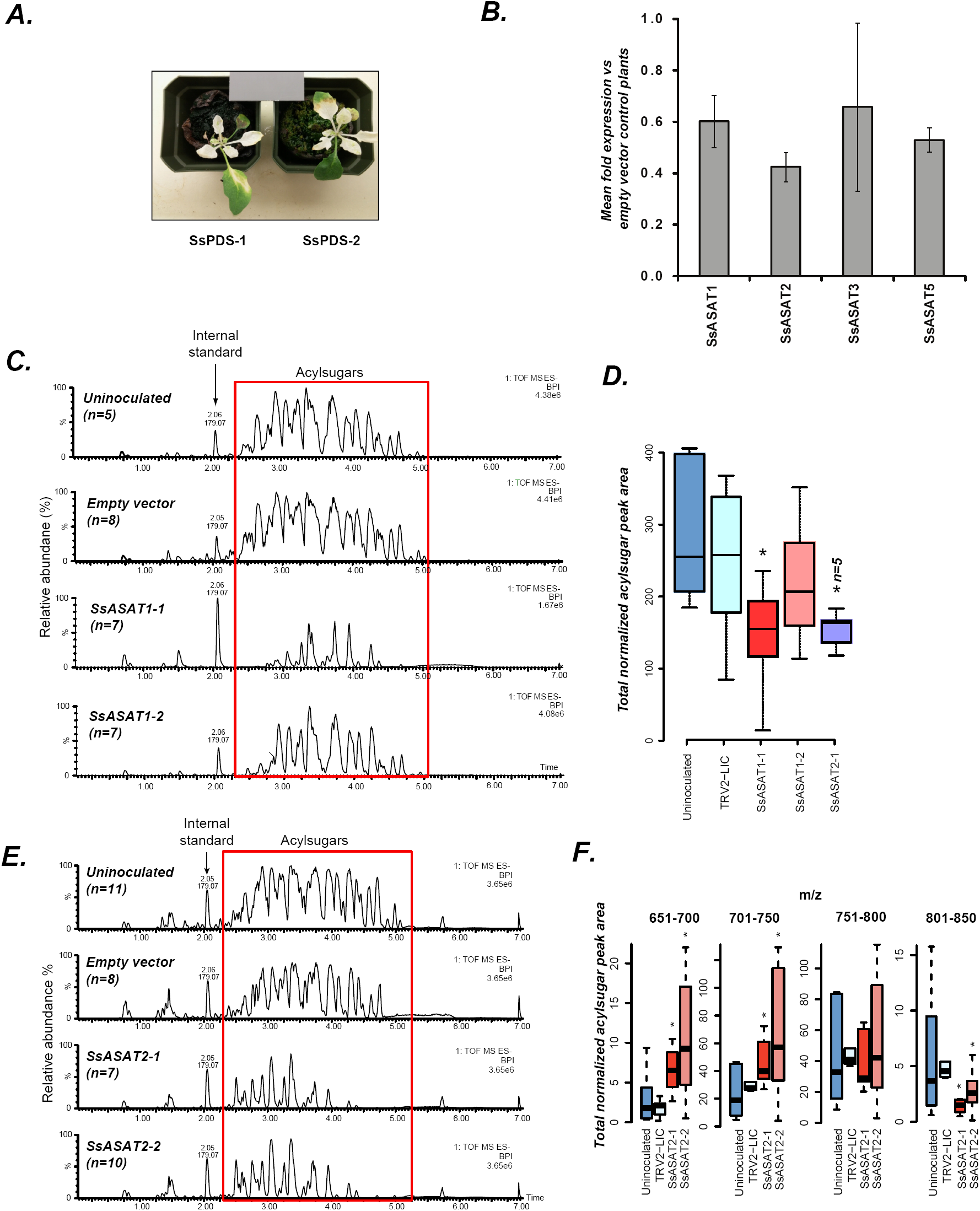
In vivo validation of SsASAT1 and SsASAT2 candidates. (A) Two representative plants with the phytoene desaturase gene silenced using VIGS shown with 18% reflectance gray card. SsPDS-1 and SsPDS-2 have two different regions of the SsPDS transcript targeted for silencing. (B) qPCR results of SsASAT VIGS lines. Relative fold change in ASAT transcript abundance in VIGS knockdown plants compared to empty vector plants. Error bars indicate standard error obtained using three technical replicates. Expression level of the phytoene desaturase (PDS) gene was used as the reference control. File Supplement 1 is the Python script used for picking the most optimal fragments and Source Data 1 includes values obtained from qPCR analysis. (C,D) SsASAT1 knockdown using two different constructs (SsASAT1-1, SsASAT1-2) shows reduction in acylsugar levels. The SsASAT1-1 phenotype is more prominent than the ASAT1-2 phenotype, being significantly lower (p=0.05; KS test). One construct for SsASAT2 (SsASAT2-1) also showed significant decrease in acylsugar levels (p=0.03; KS test). Note that the Y-axis total ion intensity in (C) is different for each chromatogram. (E,F) SsASAT2 knockdown leads to drops in levels of higher molecular weight acylsugars. In (C-F), number of plants used for statistical analysis is noted. Source Data 2 describes normalized peak areas from VIGS plants used for making these inferences. Results of the second set of biological replicate experiments performed at a different time under a different set of conditions -- as described in Table Supplement 1 -- is shown in Figure Supplement 1. Figure Supplement 2 is a more detailed analysis of the SsASAT2 knock-down phenotype showing the individual acylsugar levels under the experimental conditions.

SsASAT1 VIGS revealed statistically significant reductions in acylsugar levels in at least one construct across two experimental replicates (Kolmogorov-Smirnov [KS] test, p-value <0.05) (Figure 3C,D; Figure 3-Figure Supplement 1A), consistent with its predicted role in catalyzing the first step in acylsugar biosynthesis. *SsASAT2* expression reduction also produced plants with an overall decrease in acylsugars (Figure 3E,F; Figure 3-Figure Supplement 1B). However, these plants also showed some additional unexpected acylsugar phenotypes, namely increases in lower molecular weight acylsugars (*m/z* ratio: 651-700, 701-750; KS test p<0.05), decreases in levels of higher molecular weight products (*m/z* ratio: 751-800, 801-850) (Figure 3E,F) as well as changes in levels of some individual acylsugars as described in **Figure 3-Figure Supplement 2**. These results provide *in planta* support for the involvement of SsASAT2 in acylsugar biosynthesis, and suggest the possibility of discovering additional complexity in the future.

Silencing the *SsASAT3* transcript also led to an unexpected result - significantly higher accumulation of the normally very low abundance tri-acylsugars [S3:13(2,5,6); S3:14(2,6,6); S3:15(5,5,5); S3:16(5,5,6); S3:18(6,6,6)] and their acetylated tetra-acylsugar derivatives (KS test p<0.05), compared to empty vector control infiltrated plants (Figure 4A,B; Figure 4-Figure Supplement 1). The tetra-acylsugars contained C2 or C5 acylation on the furanose ring (Figure 4-Figure Supplement 2). These observations are consistent with the hypothesis that di-acylated sugars accumulate upon SsASAT3 knockdown and then serve as substrate for one or more other enzyme (Figure 4-Figure Supplement 3). Our working hypothesis is that this inferred activity is the as-yet-unidentified SsASAT4 activity; this is based on comparisons of *in vitro* enzyme assay products and *in vivo* purified acylsugars from Salpiglossis plants (Figure 2A) (For review only – Additional File 1). We propose that the hypothesized SsASAT4 may promiscuously acylate di-acylated sugars in addition to performing C2/C5 additions on tri-acylsugars.

**Figure 4:**
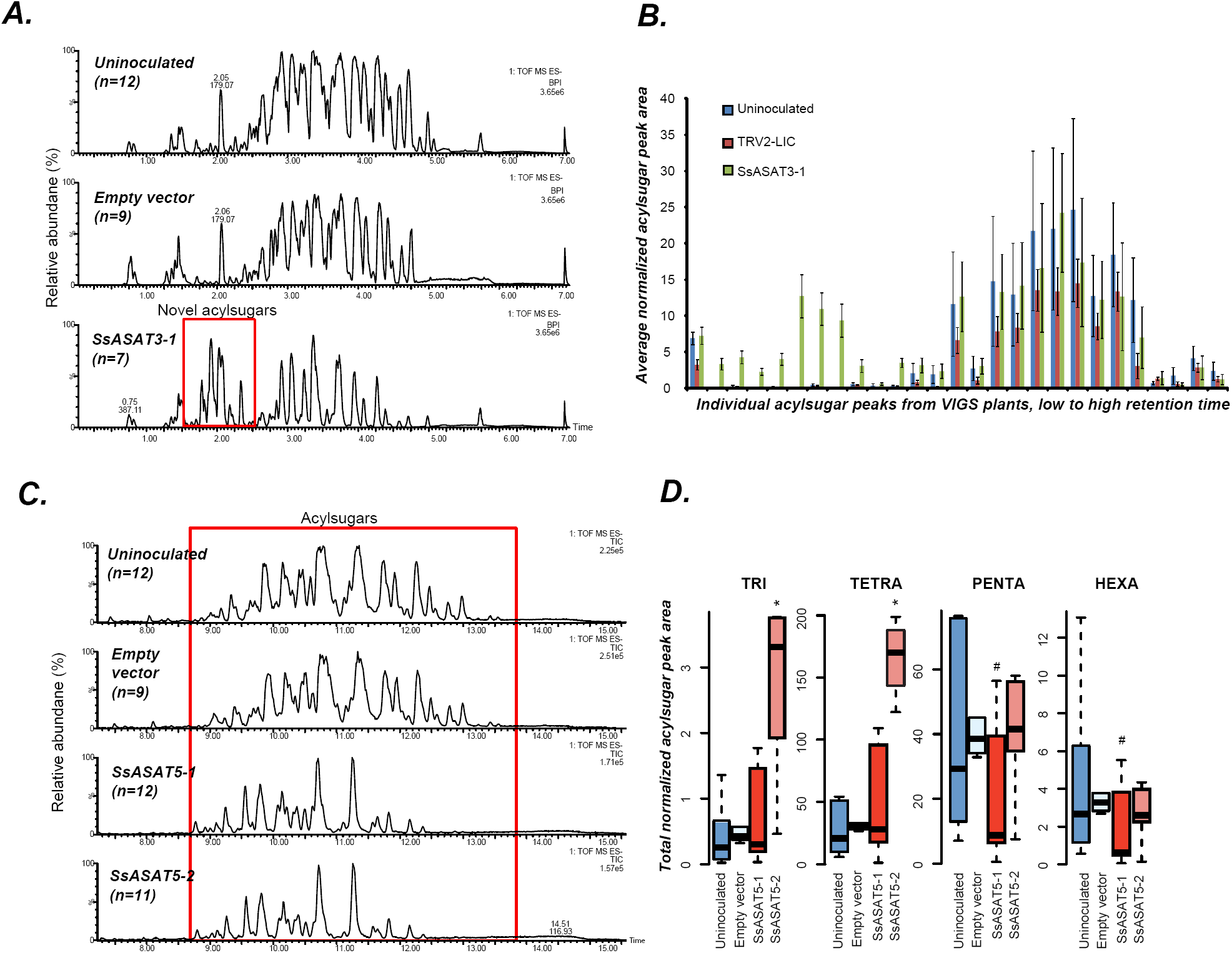
VIGS phenotypes of SsASAT3 and SsASAT5 knockdown plants. In each sub-figure, the left hand panel shows a representative chromatographic phenotype while the right hand panel shows distributions of the aggregated peak areas of all plants of the tested genotypes. (A,B) SsASAT3 VIGS knockdown experiment using a single targeting fragment SsASAT3-1 resulted in appearance of novel acylsugar peaks whose levels are significantly higher (p<0.05, KS test) vs. control. Error bars indicate standard error. Individual acylsugar peak areas are shown in Figure Supplement 1, while positive mode fragmentation patterns of the novel acylsugars are shown in Figure Supplement 2. (C) SsASAT5-1 and SsASAT5-2 chromatograms are from individual plants with two different regions of the SsASAT5 transcripts targeted for silencing. (D) Distributions of the aggregated peak areas of all plants of the tested genotypes. The boxplots show that SsASAT5 knockdown leads to a significant (*: KS test p<0.05; #: KS test 0.05<p<0.1) accumulation of tri-and tetra-acylsugars, and the effect is prominent in the SsASAT5-2 construct. A graphical explanation of the SsASAT3 and SsASAT5 knockdown results is presented in Figure Supplement 3.

SsASAT5, based on *in vitro* analysis, was proposed to catalyze acetylation of tetra- to penta-acylsugars. As expected, its knockdown led to accumulation of tri- and tetra-acylsugars (Figure 4C,D; Figure 4-Figure Supplement 3B). The accumulating tetra-acylsugars were C2/C5 furanose ring-acylated derivatives of the tri-acylsugars, suggesting presence of a functional SsASAT4 enzyme and further validating the annotation of the knocked down enzyme as SsASAT5 (Figure 4-Figure Supplement 3B). Thus, in summary, we identified four ASAT enzymes and validated their impact on acylsugar biosynthesis in Salpiglossis trichomes using VIGS.

Taken together, the Salpiglossis metabolites produced *in vivo*, combined with *in vitro* and RNAi results, lead to the model of the Salpiglossis acylsugar biosynthetic network shown in Figure 5. SsASAT1 – the first enzyme in the network – adds aiC5 or aiC6 to the sucrose R2 position. SsASAT2 then converts this mono-acylated sucrose to a di-acylated product, via addition of aiC5 or aiC6 at the R4 position. Four possible products are thus generated by the first two enzymes alone. Next, the SsASAT3 activity adds aiC6 at the R3 position, followed by one or more uncharacterized enzyme(s) that adds either C2 or aiC5 at the furanose ring R1′ or R3′ positions, respectively. SsASAT5 next performs acetylation at the R6′ position to produce penta-acylsugars, which can then be further converted by an uncharacterized SsASAT6 to hexa-acylsugars via C2 addition at the R3′ position.

**Figure 5:**
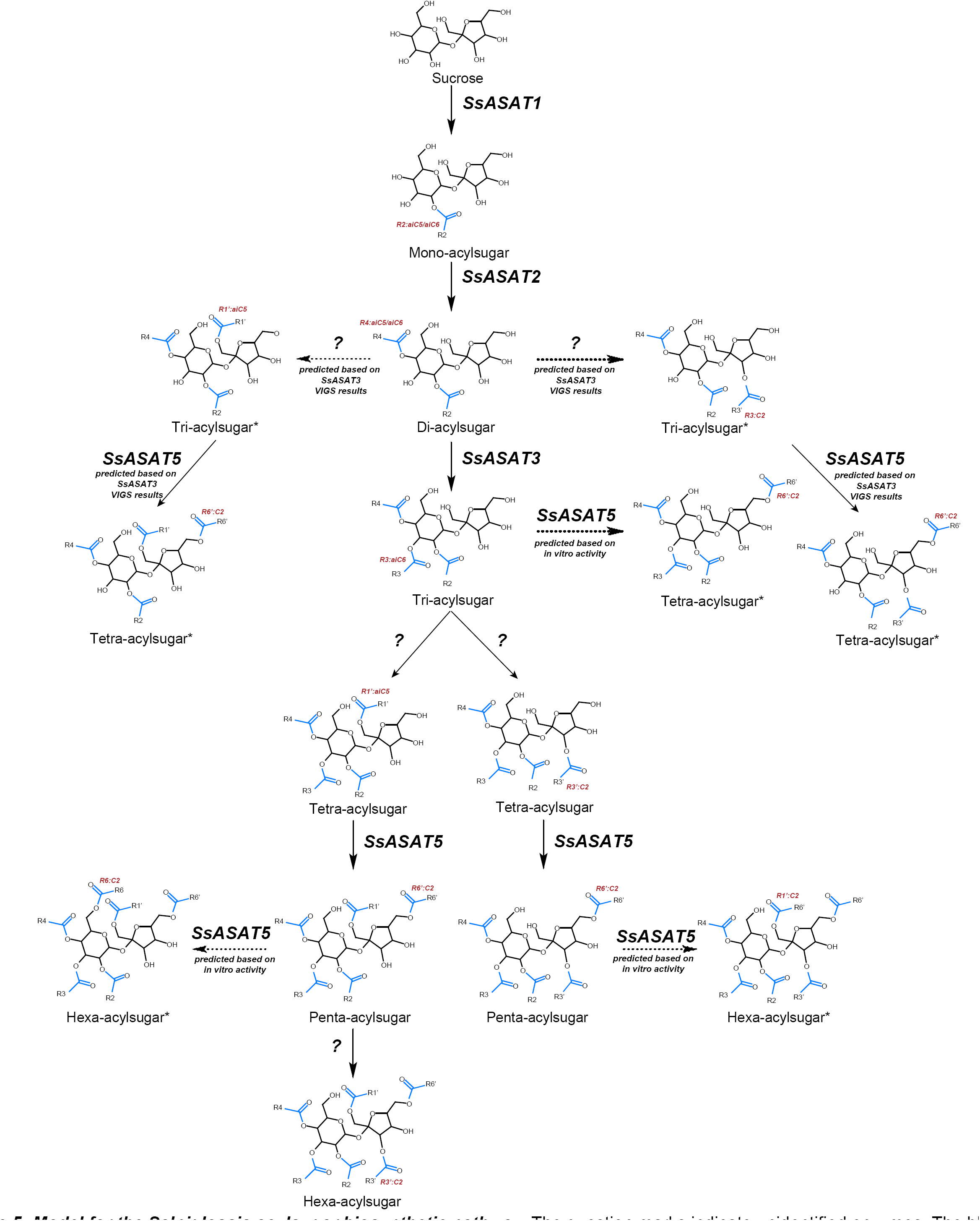
Model for the Salpiglossis acylsugar biosynthetic pathway. The question marks indicate unidentified enzymes. The blue colored acyl chains are positioned on the sucrose molecule based on results of positive mode fragmentation characteristics, co-elution assays, and comparisons with purified acylsugars. Main activities are shown in solid arrows and potential alternate activities - where acylation positions and enzymatic activities are hypothesized based on *in vitro* and *in vivo* findings - are shown in dashed arrows. An asterisk (*) next to the acylsugar names indicates no NMR structure is available for the acylsugar in Salpiglossis or Petunia, and the acyl chain positions are postulated based on their fragmentation patterns in positive and/or negative mode, and on hypothesized enzyme activities as described in the main text.

Our results are consistent with the existence of at least two additional activities – SsASAT4 and SsASAT6. These enzymes may be included in the five BAHD family candidates highly expressed in both the trichome and stem (average number of reads > 500), and thus not selected for our study because they did not meet the differential expression criterion. Also, the fact that there are >400 detectable acylsugar-like peaks in the Salpiglossis trichome extracts suggests the existence of additional ASAT activities, either promiscuous activities of characterized ASATs or of other uncharacterized enzymes. Nonetheless, identification of the four primary ASAT activities can help us to investigate the origins and evolution of the acylsugar biosynthetic pathway over time.

### The evolutionary origins of acylsugar biosynthesis

We used our analysis of SsASAT1, SsASAT2, SsASAT3 and SsASAT5 activities with information about ASATs in Petunia and tomato species (Schilmiller et al., 2012, 2015; Fan et al., 2016a) **(For review only – Additional File 2)** to infer the origins of the acylsugar biosynthetic pathway. Based on BLAST searches across multiple plant genomes, ASAT-like sequences are very narrowly distributed in the plant phylogeny (Figure 6-Figure Supplement 1). This led us to restrict our BLAST searches, which used SlASATs and SsASATs as query sequences, to species in the orders Solanales, Lamiales, Boraginales and Gentianales, all in the Lamiidae clade (Refulio-Rodriguez and Olmstead, 2014). Phylogenetic reconstruction was performed with the protein sequences of the most informative hits obtained in these searches to obtain a “gene tree”. Reconciliation of this gene tree with the phylogenetic relationships between the sampled species (Figure 6B) allowed inference of the acylsugar biosynthetic pathway before the emergence of the Solanaceae (Figure 6C; Figure 6-Figure Supplements 2A-C, 3A-C).

**Figure 6:**
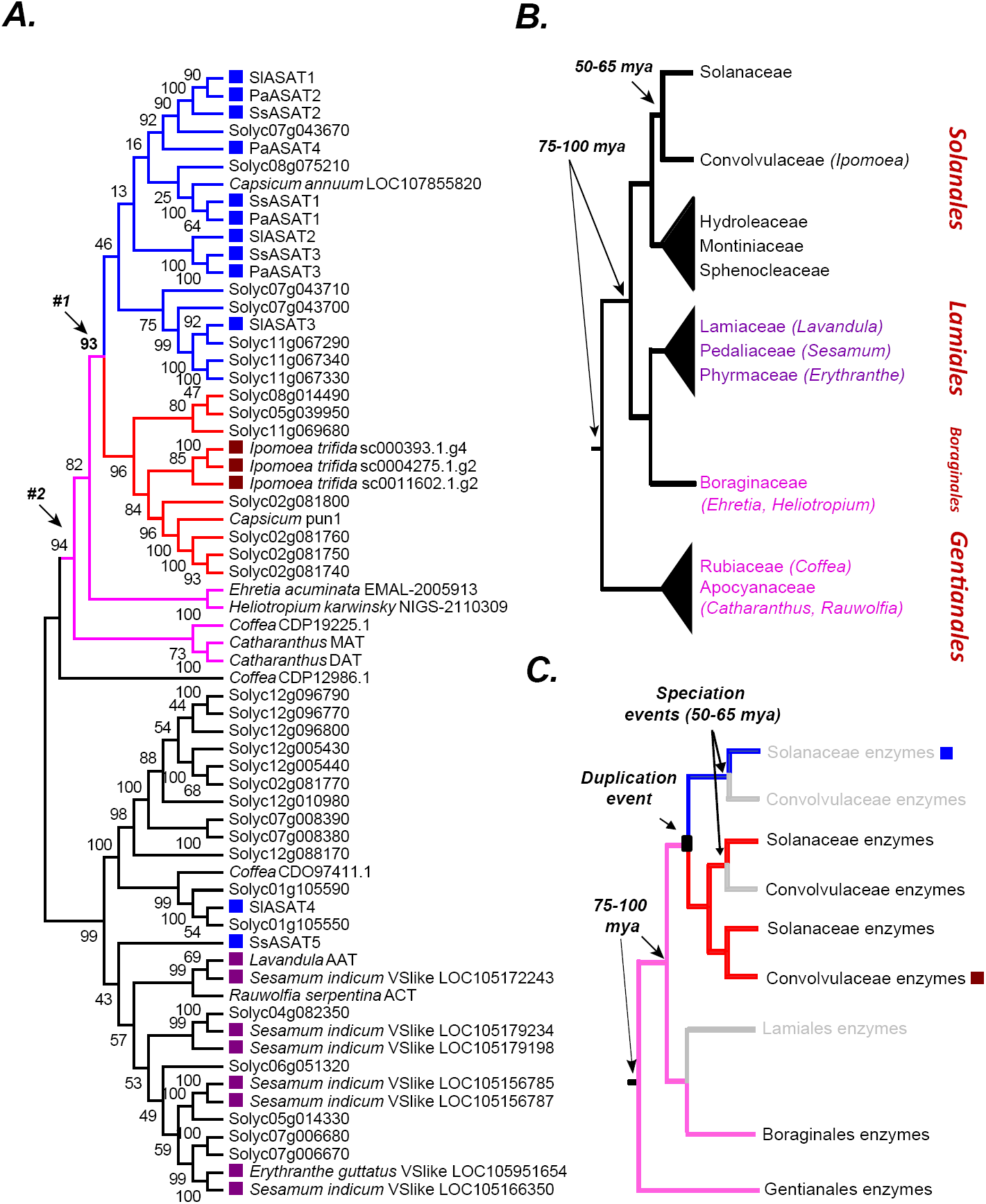
The origins of acylsugar biosynthesis. (A) Gene tree showing the characterized ASATs (blue squares) and other BAHDs in Clade III of the BAHD family of enzymes ( D’Auria, 2006). Only Lamiid species are included in this tree, given significantly similar ASAT sequences were not detected elsewhere in the plant kingdom, as shown in Figure Supplement 1. The blue sub-clade is where most ASAT activities lie, while the red monophyletic sub-clade with high bootstrap support does not contain ASAT activities. Pink sub-clade includes enzymes present in outgroup species beyond Solanales. The purple squares highlight the closest homologs in the Lamiales order. The robustness of this topology was also tested using additional tree reconstruction approaches showed in Figure Supplements 2 and 3. Source Data 1 is the alignment file in the MEGA mas format used for making this tree. (B) Known relationships between different families and orders included in the study, based on Refulio-Rodriguez and Olmstead, 2014. The times indicated are based on a range of studies as described in the main text. (C) Reconciled evolutionary history of the blue ASAT sub-clade based on (A) and (B). Grey color indicates loss or lack of BLAST hits in the analyzed sequence datasets from that lineage.

Three major subclades in the gene tree – highlighted in blue, red and pink – are relevant to understanding the origins of the ASATs (Figure 6A). A majority of characterized ASATs (**blue squares in the blue subclade**, Figure 6A) are clustered with *Capsicum* PUN1 — an enzyme involved in biosynthesis of the alkaloid capsaicin — in a monophyletic group with high bootstrap support (Group #1, red and blue subclades Figure 6A). Two of the most closely related non-Solanales enzymes in the tree — *Catharanthus roseus* minovincinine-19-O-acetyltransferase (MAT) and deacetylvindoline-4-O-acetyltransferase (DAT) — are also involved in alkaloid biosynthesis (Magnotta et al., 2007). This suggests that the blue ASAT subclade emerged from an alkaloid biosynthetic enzyme ancestor.

A second insight from the gene tree involves Salpiglossis SsASAT5 and tomato SlASAT4, which reside outside of the blue subclade. Both enzymes catalyze C2 addition on acylated sugar substrates in downstream reactions of their respective networks. Multiple enzymes in this region of the phylogenetic tree (Figure 2-Figure Supplement 1C; light blue clade) are involved in *O*-acetylation of diverse substrates e.g. indole alkaloid 16-epivellosimine (Bayer et al., 2004), the phenylpropanoid benzyl alcohol (D’Auria et al., 2002) and the terpene geraniol (Shalit et al., 2003). This observation is consistent with the hypothesis that *O*-acetylation activity was present in ancestral enzymes within this region of the phylogenetic tree.

Gene tree reconciliation with known relationships between plant families and orders (Figure 6B) was used to infer acylsugar pathway evolution in the context of plant evolution. We used historical dates as described by Särkinen and co-workers (Särkinen et al., 2013) in our interpretations, as opposed to a recent study that described a much earlier origination time for the Solanaceae (Wilf et al., 2017). Based on known relationships, Convolvulaceae is the closest sister family to Solanaceae; however, we found no putative ASAT orthologs in any searched Convolvulaceae species. The closest Convolvulaceae homologs were found in the *Ipomoea trifida* genome (Hirakawa et al., 2015) in the red subclade. This suggests that the blue and red subclades arose via a duplication event before the Solanaceae-Convolvulaceae split, estimated to be ∼50-65 mya (Särkinen et al., 2013). The lack of ASAT orthologs in Convolvulaceae could indicate that Convolvulaceae species with an orthologous acylsugar biosynthetic pathway were not sampled in our analysis. Alternatively, the absence of orthologs of the blue subclade enzymes in the Convolvulaceae, coupled with the lack of acylsugars in the sampled Convolvulaceae species (Figure 1B), leads us to hypothesize that ASAT orthologs were likely lost early in Convolvulaceae evolution.

The known phylogenetic relationships between species can help assign a maximum age for the duplication event leading to the blue and red subclades. Database searches identified homologs from species in the Boraginales (*Ehretia*, *Heliotropium*) and Gentianales (*Coffea, Catharanthus*) orders that clustered outside Group #1, in a monophyletic group with high bootstrap support (Group #2, pink branches, Figure 6A). Boraginales is sister to the Lamiales order (Figure 6B), and although Boraginales putative orthologs were identified through BLAST searches, no orthologs from Lamiales species were detected. The closest Lamiales (*Sesamum, Lavandula, Mimulus*) homologs were more closely related to SsASAT5 than to the ASATs in the blue subclade (**purple squares**, Figure 6A). The most parsimonious explanation for these observations is that the duplication event that gave rise to the blue and red subclades occurred before the Solanaceae-Convolvulaceae split 50-65 mya but after the Solanales-Boraginales/Lamiales orders diverged 75-100 mya (Hedges et al., 2006) (Figure 6B,C).

Overall, these observations support a view of the origins of the ASAT1,2,3 blue subclade from an alkaloid biosynthetic ancestor via a single duplication event >50mya, prior to the establishment of the Solanaceae. This duplicate would have undergone further rounds of duplication to generate the multiple ASATs found in the blue subclade. Thus, a logical next question is ‘what was the structure of the acylsugar biosynthetic network early in the Solanaceae family evolution?’ To address this, we focused on ASAT evolution across the Solanaceae family.

### Evolution of acylsugar biosynthesis in the Solanaceae

ASAT enzymes in the blue subclade represent the first three steps in the acylsugar biosynthetic pathway. The likely status of these three steps in the Salpiglossis-Petunia-Tomato last common ancestor (LCA) was investigated using the gene tree displayed in Figure 7A. Mapping the pathway enzymes on the gene tree suggests that the LCA likely had at least three enzymatic activities, which we refer to as ancestral ASAT1 (aASAT1), ASAT2 (aASAT2) and ASAT3 (aASAT3); these are shown in Figure 7A as red, dark blue and yellow squares, respectively. We further traced the evolution of the aASAT1,2,3 orthologs in the Solanaceae using existing functional data and BLAST-based searches. These results reveal that the aASAT1 ortholog (red squares, Figure 7A) was present until the *Capsicum*-*Solanum* split ∼17 mya (Särkinen et al., 2013) and was lost in the lineage leading to *Solanum*. This loss is evident both in similarity searches and in comparisons of syntenic regions between genomes of Petunia, *Capsicum* and tomato (Figure 7B). On the other hand, the aASAT2 (dark blue squares, Figure 7A) and aASAT3 orthologs (yellow squares, Figure 7A) have been present in the Solanaceae species genomes at least since the last common ancestor of Salpiglossis-Petunia-Tomato ∼25 mya, and perhaps even in the last common ancestor of the Solanaceae family.

**Figure 7:**
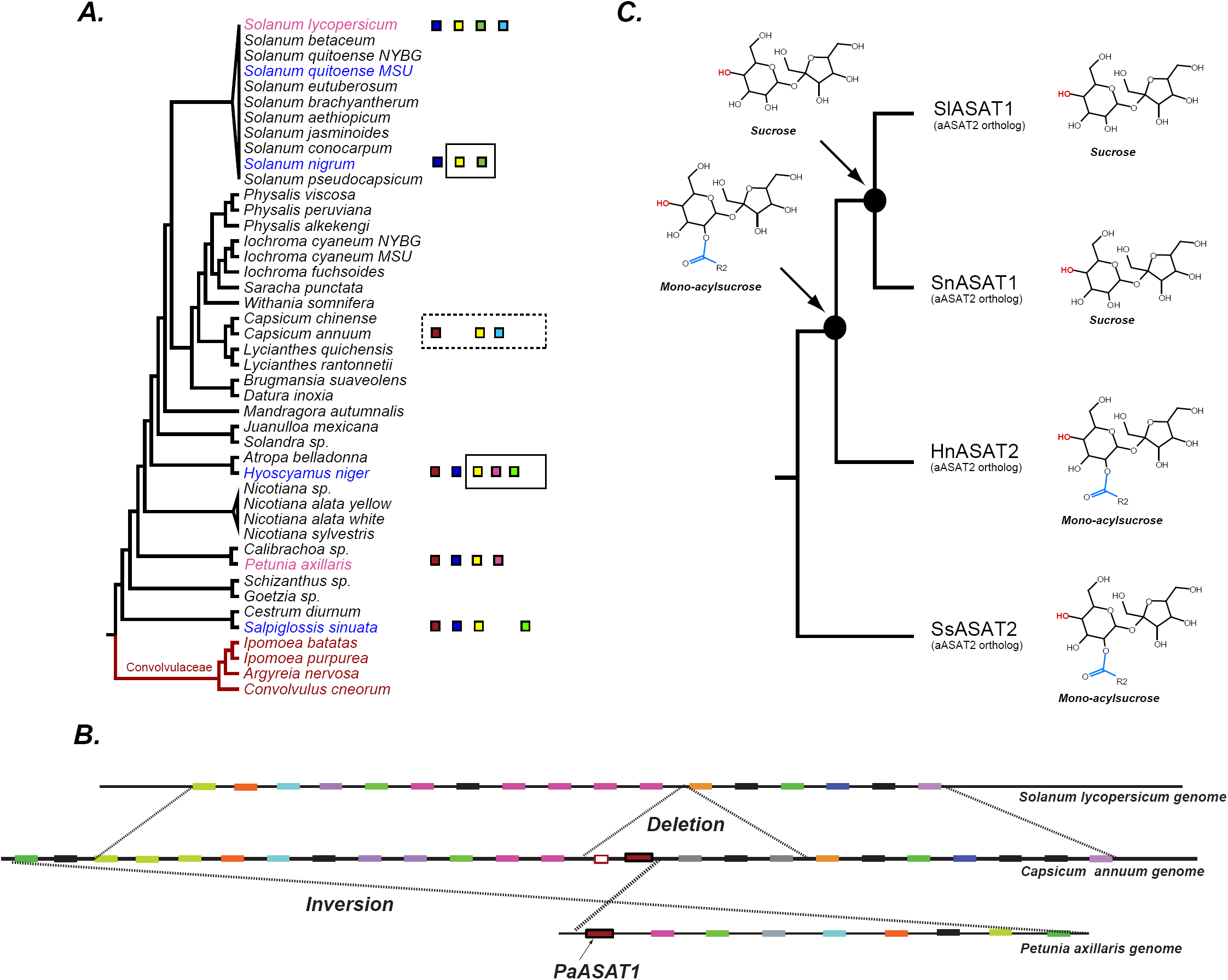
The evolution of acylsugar biosynthesis in Solanaceae. (A) ASAT activities overlaid on the Solanaceae phylo-genetic relationships. Each colored square represents a single ASAT, starting from ASAT1 and moving sequentially down the pathway to ASAT5 (left to right, sequentially). Homologs are represented by the same color. Squares not contained within a box are experimentally validated activities. A solid black box indicates that a highly identical transcript exists in the RNA-seq dataset and is trichome-high. A dashed box indicates that, based on a BLAST search, the sequence exists in the genome for the contained enzymes. *In vitro* validated *S. nigrum* and *H. niger* ASAT activities, including their phylogenetic positions, are shown in Figure Supplement 1. Species selected for RNA-seq in this study are colored blue, previously studied species are highlighted in pink and Convolvulaceae species are in red. See Source Data 1 for results of the BLAST analysis. (B) Ortholo-gous genomic regions between three species harboring aASAT1 orthologs. Each gene in the region is shown by a colored block. Orthologous genes are represented by the same color. The PaASAT1 gene (red box) has two homologous sequences in the Capsicum syntenic region, but one of them is truncated. aASAT1 ortholog is not seen in the syntenic region in tomato. Genes used to make this figure are described in Source Data 2. (C) Substrate utilization of aASAT2 orthologs from multiple species is described based on the activities presented in Figure Supplement 1. *Hyoscyamus niger* (Hn), *Salpiglossis sinuata* (Ss), *Solanum nigrum* (Sn) and *Solanum lycopersicum* (Sl). The hydroxyl group highlighted in red shows the predicted position of acylation by the respective aASAT2 ortholog.

One inference from this analysis is that aASAT2 orthologs switched their activity from ASAT2-like acylation of mono-acylsucroses in Salpiglossis/Petunia to ASAT1-like acylation of unsubstituted sucrose in cultivated tomato (Figure 7A). Interestingly, despite the switch, both aASAT2 and aASAT3 orthologs in tomato continue to acylate the same pyranose-ring R4 and R3 positions, respectively (Figure 7C; Figure 8). In addition, cultivated tomato has two ‘new’ enzymes — SlASAT3 and SlASAT4 — which were not described in Petunia or Salpiglossis acylsugar biosynthesis.

**Figure 8:**
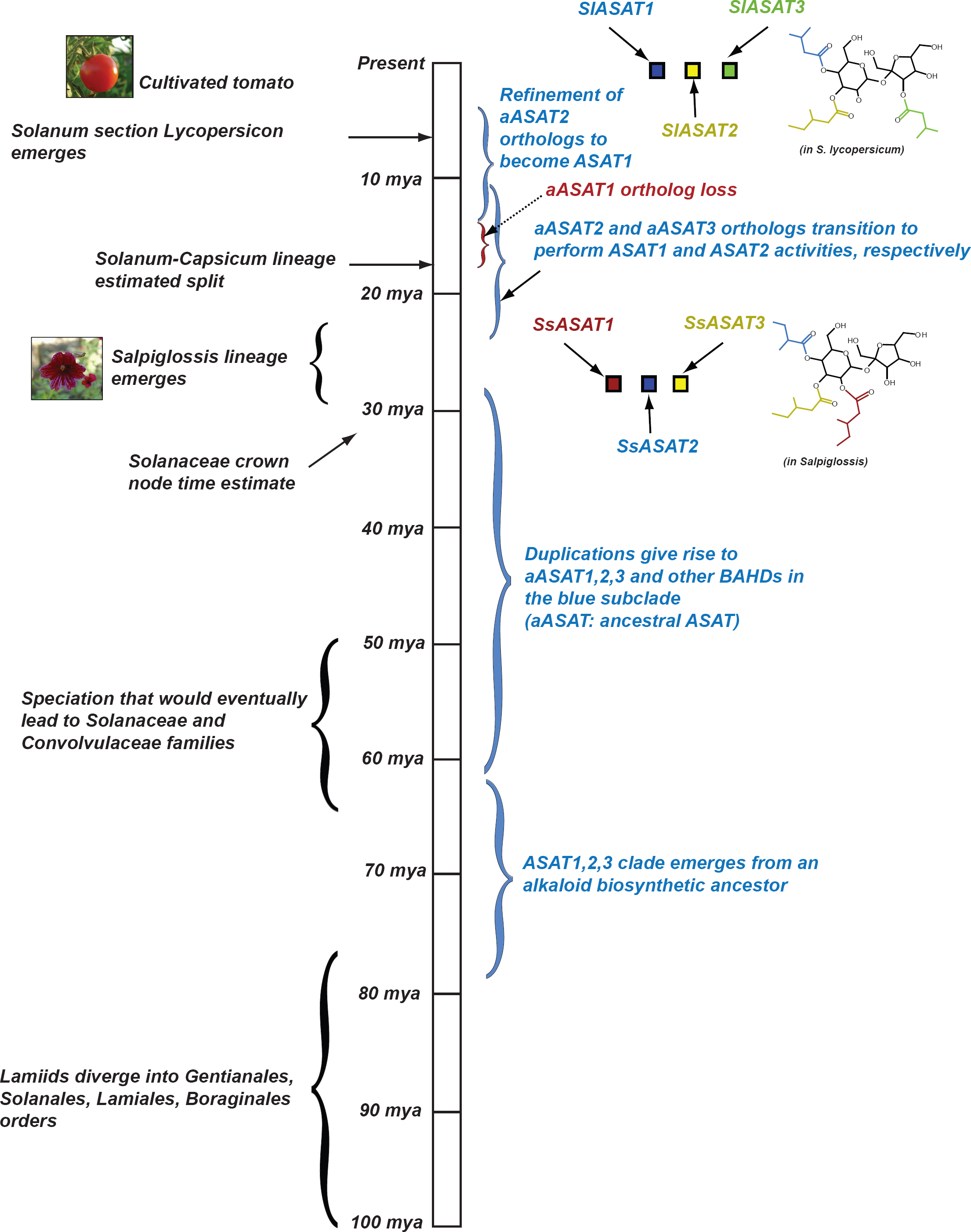
Proposed model for evolution of acylsugar biosynthesis over 100 million years. Events in plant evolution are noted on the left and the acylsugar biosynthetic pathway developments are noted on the right of the timeline. Tri-acylsugars produced by Salpiglossis and tomato ASAT1,2,3 enzymes are shown with the chain color corresponding to the color of the enzyme that adds the chain. Images obtained from Wikimedia Commons under Creative Commons CC BY-SA 3.0 license. We note that Salpiglossis and SsASAT activities should not be considered as “ancestral” and Tomato and SlASAT activities as “modern”, since all of them exist today. We use the word “lineages” (eg: Salpiglossis lineage) to refer to the branch leading up to the currently existing (extant) species.

The functional transitions of aASAT2 and aASAT3 could have occurred via (i) functional divergence between orthologs or (ii) duplication, neo-functionalization and loss of the ancestral enzyme. Counter to the second hypothesis, we found no evidence of aASAT2 ortholog duplication in the genomic datasets; however, we cannot exclude the possibility of recent polyploidy in extant species producing duplicate genes with divergent functions. We explored the more parsimonious hypothesis that functional divergence between orthologs led to the aASAT2 functional switch.

Functional divergence between aASAT2 orthologs may have occurred by one of two mechanisms. One, it is possible that aASAT2 orthologs had weak/strong sucrose acylation activity prior to aASAT1 loss. Alternatively, sucrose acylation activity arose completely anew after the loss of aASAT1. We sought evidence for acceptor substrate promiscuity in extant species by characterizing the activity of additional orthologous aASAT2 enzymes from *H. niger* and *S. nigrum* (Figure 7C) using aiC6 and nC12 as acyl CoA donors. HnASAT2 — like SsASAT2 — only performed the ASAT2 reaction (acylation of mono-acylsucrose), without any evidence for sucrose acylation under standard testing conditions (Figure 7C, Figure 7-Figure Supplement 1A-D). However, both SnASAT1 and SlASAT1 could produce S1:6(6) and S1:12(12) from sucrose.

Overall, these results suggest that up until Atropina-Solaneae common ancestor, the aASAT2 ortholog still primarily conducted the ASAT2 activity (Figure 7C). However, at some point, the aASAT2 ortholog moved in reaction space towards being ASAT1. This activity shift was likely complete by the time the *S.nigrum*-*S.lycopersicum* lineage diverged (Figure 7C), given that aASAT2 orthologs in both species utilize sucrose. Future detailed characterization of aASAT2 orthologs from phylogenetically intermediate, pre-and post-aASAT1 loss species can help pinpoint the route taken by the aASAT2 ortholog in its functional diversification as well as determine whether positive selection played a role.

## Conclusion

We performed experimental and computational analysis of BAHD enzymes from several lineages spanning ∼100 million years, and characterized the emergence and evolution of the acylsugar metabolic network. We identified four biosynthetic enzymes in *Salpiglossis sinuata*, the extant species of an early emerging Solanaceae lineage, characterized *in vitro* activities, validated *in planta* roles of these ASATs and studied their emergence over 100 million years of plant evolution. These results demonstrate the value of leveraging genomics, phylogenetics, analytical chemistry and enzymology with a mix of model and non-model organisms to understand the evolution of biological complexity.

Based on our findings, the acylsugar phenotype across the Solanaceae is “hyper-diverse”, meaning there is large diversity within and between closely related species. Previous studies indicate that all eight hydroxyls on the sucrose core are subject to acylation (Ghosh et al., 2014; Schilmiller et al., 2015; Fan et al., 2016a). A simple calculation (see Supplemental Methods) considering only 12 aliphatic CoA donors suggests >6000 theoretical possible structures for tetra-acylsugars alone. This estimate does not even include estimates of tri-, penta-, hexa-acylsugars nor does it consider non-aliphatic CoAs such as malonyl CoA, esters of other sugars such as glucose or positional isomers. Although the theoretical possibilities are restricted by availability of CoAs in trichome tip cells and existing ASAT activities, we still observe hundreds of acylsugars across the Solanaceae, with >400 detectable acylsugar-like chromatographic peaks in single plants of Salpiglossis and *N. alata*, including very low abundance hepta-acylsugars and acylsugars containing phenylacetyl and tigloyl chains **(For review only – Additional File 1)**. This diversity may be less than among some terpenoid classes e.g. >850 diterpenes have been described in different members of the *Euphorbia* genus alone (Vasas and Hohmann, 2014). However, the diversity is significantly greater than other specialized metabolite classes restricted to a single clade such as glucosinolates in Brassicales (∼150 across the order) (Olsen et al., 2016), betalains in Caryophyllales (∼75 non-glycosylated betalains across the order) (Khan and Giridhar, 2015) and acridone alkaloids in Rutaceae (∼100 in the family) (Roberts et al., 2010). Acylsugar diversity may be higher than, or at least comparable to, resin glycosides in Convolvulaceae (>250 structures across the family) (Pereda-Miranda et al., 2010) and pyrrolizidine alkaloids in Boraginaceae (>225 structures across the family) (El-Shazly and Wink, 2014). Much of the observed acylsugar phenotypic complexity arises due to the diversity in acyl chains — their types, lengths, isomeric forms and positions on the sucrose molecule.

Does such immense diversity within the same metabolite class serve a purpose? Glucosinolate diversity was shown to influence fitness in the wild (Prasad et al., 2012) but pyrrolizidine alkaloid diversity may not be so relevant (Macel et al., 2002). While acylsugar diversity is ecologically relevant to some extent (Weinhold and Baldwin, 2011; Leckie et al., 2016; Luu et al., 2017), it is unlikely that each novel acylsugar has a specific function. The diversity — perhaps occurring as an unintended consequence of ASAT promiscuity — may, in some cases, also provide benefit in the form of physical defense (e.g. stickiness).

We studied the question of how acylsugar biosynthesis originated. The results provide evidence for repeated cycles of innovation involving potentiation, actualization and refinement that contributed to the origin and diversification of the acylsugar biosynthetic pathway (Blount et al., 2012) (Figure 8). We propose that the ASAT enzymes catalyzing sucrose acylation first emerged (or actualized) between 30-80 mya. Events prior to the emergence of this activity — including existence of alkaloid biosynthetic BAHDs and the duplication event 60-80 mya that gave rise to the blue and the red subclades — potentiated the emergence of the ASAT activities. Presumably, acylsugars provided a fitness advantage to the ancestral plants, leading to the refinement of ASAT activities over the next few million years. For acylsugar production to emerge, these steps in enzyme evolution would have been accompanied by parallel innovations in ASAT transcriptional regulation leading to gland cell expression as well as production of precursors (i.e., acyl CoA donors and sucrose) in Type I/IV trichomes.

Since the origin of the Solanaceae, this pathway experienced another round of major innovation (Figure 8) leading to the aASAT2 functional switch. We propose that this cycle of innovation was made possible by the potentiation provided by the similarity in activities of aASAT1,2 and 3, conducive to the emergence of promiscuity. Thus, the sucrose acylation activity may have already emerged in aASAT2 before the aASAT1 ortholog loss. It is unclear whether the loss of aASAT1 was a result of it being rendered redundant by the emergence of the aASAT2 promiscuous activity or an unrelated loss event. Regardless, this loss event was associated with the reorganization of acylsugar biosynthesis, and perhaps was followed by a refinement in the actualized aASAT2 sucrose acylation activity. Additional sampling of intermediate species can help test this model and determine the dynamics that played out between aASAT1,2,3 orthologs between 20-15 my. We cannot rule out the possibility of aASAT2 switching to ASAT1 after the aASAT1 ortholog loss. However, this model is less parsimonious, because it needs the emergence of the ASAT1 activity to occur “just in time” before the entire downstream pathway decayed or diversified through drift. In addition to this second cycle of innovation, previous results from closely related *Solanum* species also revealed more recent cycles involving gene duplication, regulatory divergence of ASATs (Schilmiller et al., 2015) and neo-functionalization of duplicates (Ning et al., 2015) in influencing the phenotypes produced by this pathway. These findings highlight the striking plasticity of the acylsugar biosynthetic network and illustrate the many ways by which chemical novelty emerges in specialized metabolism.

In this study, we used an integrative strategy combining high-throughput technologies and traditional biochemistry. *S. lycopersicum* is the flagship species of the Solanaceae family, and we were able to apply the insights derived with this crop as a foundation to discover novel enzymes in related species. Our investigations in Salpiglossis were also aided by the availability of ASATs and NMR structures of their products in Petunia, which is more closely related to Salpiglossis than tomato. “Anchor species” such as Petunia, which are phylogenetically distant from flagship/model species, can enable study of a different region of the phylogenetic tree of the clade of interest. Development of limited genomic and functional resources in such anchor species, coupled with integrative, comparative approaches can offer more efficient routes for the exploration of biochemical complexity in the ∼300,000 plant species estimated to exist on our planet (Mora et al., 2011).

## Methods

### Plant acylsugar extractions and mass spectrometric analyses

Acylsugar extractions were carried out from plants at the New York Botanical Gardens and from other sources (Figure 1-File Supplement 1). The plants sampled were at different developmental stages and were growing in different environments. The extractions were carried out using acetonitrile:isopropanol:water in a 3:3:2 proportion similar to previous descriptions (Schilmiller et al., 2010; Ghosh et al., 2014; Fan et al., 2016a) with the exception of gently shaking the tubes by hand for 1-2 minutes. All extracts were analyzed on LC/MS using 7-min, 22-min or 110-min LC gradients on Supelco Ascentis Express C18 (Sigma Aldrich, USA) or Waters BEH amide (Waters Corporation, USA) columns (Supplemental Table 1), as described previously (Schilmiller et al., 2010; Ghosh et al., 2014; Fan et al., 2016a). While the 110-min method was used to minimize chromatographic overlap in support of metabolite annotation in samples with diverse mixtures of acylsugars, targeted extracted ion chromatogram peak area quantification was performed using the 22-min method data. The QuanLynx function in MassLynx v4.1 (Waters Corporation, USA) was used to integrate extracted ion chromatograms for manually selected acylsugar and internal standard peaks. Variable retention time and chromatogram mass windows were used, depending on the experiment and profile complexity. Peak areas were normalized to the internal standard peak area and expressed as a proportion of mg of dry weight.

### RNA-seq data analysis

RNA was extracted and sequenced as described in the **Supplemental Methods**. The mRNA-seq reads were adapter-clipped and trimmed using Trimmomatic v0.32 using the parameters (*LEADING:20 TRAILING:20 SLIDINGWINDOW:4:20 MINLEN:50*). The quality-trimmed reads from all datasets of a species were assembled *de novo* into transcripts using Trinity v.20140413p1 after read normalization (max_cov=50,KMER_SIZE=25,max-pct-stdev=200,SS_lib_type=RF) (Grabherr et al., 2011). We tested three different kmer values (k=25, 27, 31) and selected the best kmer value for each species based on contig N50 values, BLASTX hits to the *S. lycopersicum* annotated protein sequences and CEGMA results (Parra et al., 2007). A minimum kmer coverage of 2 was used to reduce the probability of erroneous or low abundance kmers being assembled into transcripts. After selecting the best assembly for each species, we obtained a list of transcripts differentially expressed between trichomes and stem/petiole for each species using RSEM/EBSeq (Li and Dewey, 2011; Leng et al., 2013) at an FDR threshold of p<0.05. The differential expression of five transcripts in *S. quitoense* and four transcripts in Salpiglossis was confirmed using semi-quantitative RT-PCR, along with the PDS as negative control (Table 1-Figure Supplement 1).

### Purification and structure elucidation of acylsugars using NMR

For metabolite purification, aerial tissues of 28 Salpiglossis plants (aged 10 weeks) were extracted in 1.9 L of acetonitrile:isopropanol (AcN:IPA, v/v) for ∼10 mins, and ∼1 L of the extract was concentrated to dryness on a rotary evaporator and redissolved in 5 mL of AcN:IPA. Repeated injections from this extract were made onto a Thermo Scientific Acclaim 120 C18 HPLC column (4.6 x 150 mm, 5 μm particle size) with automated fraction collection. HPLC fractions were concentrated to dryness under vacuum centrifugation, reconstituted in AcN:IPA and combined according to metabolite purity as assessed by LC/MS. Samples were dried under N_2_ gas and reconstituted in 250 or 300 μL of deuterated NMR solvent CDCl_3_ (99.8 atom % D) and transferred to solvent-matched Shigemi tubes for analysis. ^1^H, ^13^C, *J*-resolved ^1^H, gCOSY, gHSQC, gHMBC and ROESY NMR experiments were performed at the Max T. Rogers NMR Facility at Michigan State University using a Bruker Avance 900 spectrometer equipped with a TCI triple resonance probe. All spectra were referenced to non-deuterated CDCl_3_ solvent signals (δ_H_ = 7.26 and δ_C_ = 77.20 ppm).

### Candidate gene amplification and enzyme assays

Enzyme assays were performed as previously described (Fan et al., 2016b) with the following modifications: Enzyme assays were performed in 30 μL reactions (3μL enzyme + 3μL 1mM acyl CoA + 1μL 10mM (acylated) sucrose + 23μL 50mM NH_4_Ac buffer pH 6.0) or 20μL reactions with proportionately scaled components. Reactions that used an NMR-characterized substrate were performed with 0.2μL substrate and 23.8μL buffer. Reaction products were characterized using Waters Xevo G2-XS QToF LC/MS (Waters Corporation, USA) using previously described protocols (Fan et al., 2016b).

### Characterizing the SsASAT5 activity

Identification of SsASAT5 activity required a significant amount of the starting substrate, however, standard sequential reactions used to isolate SsASAT1,2,3 activities do not produce enough starting substrate for SsASAT5. Hence, we used S3:15(5,5,5) purified from a back-crossed inbred line (BIL6180, (Ofner et al., 2016)) whose NMR resolved structure shows R2, R3, R4 positions acylated by C5 chains (unpublished data) — same as Salpiglossis tri-acylsugars. SsASAT5 could use this substrate to acylate the furanose ring (Figure 2-Figure Supplement 6A,B), however, the tetra-acylsugar thus produced is not further acetylated to penta-acylsugar, suggesting the C2 acylation by SsASAT5 is not the same as the C2 acylation on tetra-acylsugars *in vivo*. SsASAT5 also acetylated the acceptors S4:21(5,5,5,6) and S4:19(2,5,6,6) purified from Salpiglossis trichome extracts on the furanose ring to penta-acylsugars that co-migrated with the most abundant penta-acylsugars purified from the plant (Figure 2C,D). Finally, S5:23(2,5,5,5,6) and S5:21(2,2,5,6,6) purified from plant extracts were enzymatically acetylated to hexa-acylsugars, which co-migrated with two low abundance hexa-acylsugars from the plant (Figure 2-Figure Supplement 6C,D). Positive mode fragmentation patterns suggested that SsASAT5 possessed the capacity to acetylate both pyranose (weak) and furanose rings of the penta-acylsugars (Figure 2-Figure Supplement 6E,F).

### Virus induced gene silencing

Primers to amplify fragments of transcripts for VIGS were chosen to minimize probability of cross silencing other Salpiglossis transcripts due to sequence similarity or due to homopolymeric regions (Figure 2-File Supplement 1,2). This was achieved using a custom Python script (Figure 3-File Supplement 1), which divided the entire transcript of interest *in silico* into multiple overlapping 20nt fragments, performed a BLAST against the Salpiglossis transcriptome and tomato genome and flagged fragments with >95% identity and/or >50% homopolymeric stretches. Contiguous transcript regions with >12 unflagged, high-quality fragments were manually inspected and considered for VIGS. 1-2 300bp regions were cloned into the pTRV2-LIC vector (Dong et al., 2007) and transformed into *Agrobacterium tumefaciens* GV3101. Agroinfiltration of Salpiglossis plants was performed as described previously (Velásquez et al., 2009) using the prick inoculation method. Salpiglossis phytoene desaturase was used as the positive control for silencing. Empty pTRV2-LIC or pTRV2 vectors were used as negative controls. Each VIGS trial was done slightly differently while modifying the growth, transformation and maintenance conditions of this non-model species. The experimental details, including the optimal growth and VIGS conditions that give the fastest results, are noted in **Figure 3-Table Supplement 1**.

### Phylogenetic inference

All steps in the phylogenetic reconstruction were carried out using MEGA6 (Tamura et al., 2013). Amino acid sequences were aligned using Muscle with default parameters. Maximum likelihood and/or neighbor joining (NJ) were used to generate phylogenetic trees. For maximum likelihood, the best evolutionary model (JTT[Jones-Taylor-Thornton]+G+I with 5 rate categories) was selected based on the Akaike Information Criterion after screening several models available in the MEGA6 software, while for NJ, the default JTT model was used. Support values were obtained using 1000 bootstrap replicates, however trees obtained using 100 bootstrap replicates also showed similar overall topologies. Trees were generated either using “complete deletion” or “partial deletion with maximum 30% gaps/missing data” options for tree reconstruction. Sequences <350aa (eg: Solyc10g079570) were excluded from this analysis.

### Data release

All RNA-seq datasets are deposited in the NCBI Short Read Archive under the BioProject PRJNA263038. Coding sequences of ASATs are provided in Figure 2-File Supplement 1 and have been deposited in GenBank (KY978746-KY978750). RNA-seq transcripts and orthologous group membership information has been uploaded to Dryad (provisional DOI: 10.5061/dryad.t7r64).

## Acknowledgements

We thank Dr. Rachel Meyers, Dr. Amy Litt and the New York Botanical Garden staff for permission and assistance in sample collection. We also thank the support staff at the Research Technology Support Facility at MSU for transcriptome sequencing and mass spectrometry assistance, and the Max T. Rogers NMR Facility for assistance with NMR data generation. We acknowledge Daniel Lybrand for cloning the *Solanum nigrum ASAT1* gene, and Dr. Anthony Schilmiller and Dr. Pengxiang Fan for assisting with sample collection at NYBG. We also acknowledge the valuable feedback received from members of the Last lab, Dr. Cornelius Barry and Dr. Eran Pichersky in the research design of this manuscript. This work was funded by National Science Foundation grants IOS–1025636 and IOS-PGRP-1546617 to R.L.L. and A.D.J. and National Institute of General Medical Sciences of the National Institutes of Health graduate training grant no. T32–GM110523 to B.L. A.D.J. acknowledges support from the USDA National Institute of Food and Agriculture, Hatch project MICL-02143.

---

**Figure 1- Figure Supplement 1:**
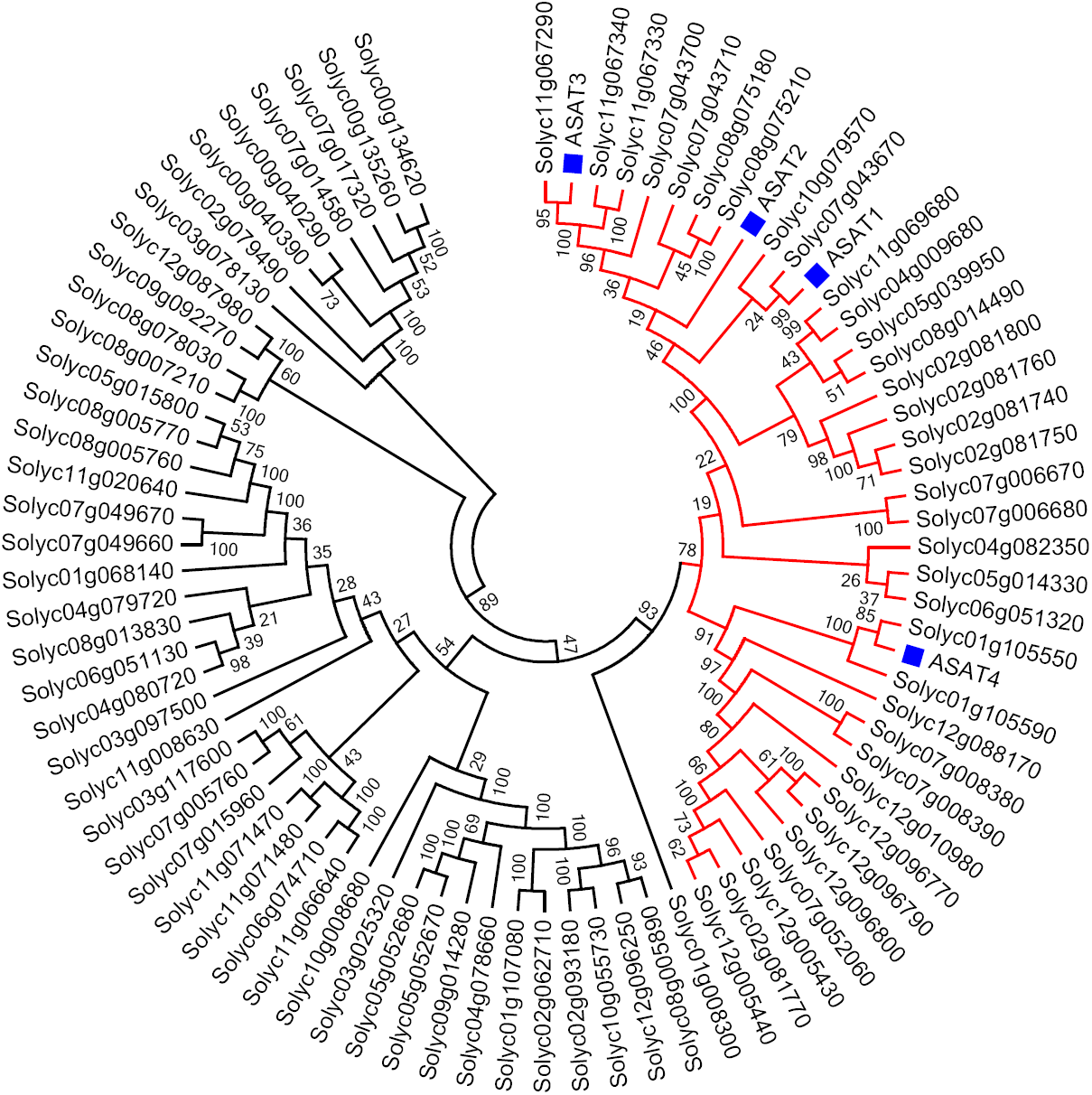
Tomato ASATs are members of the BAHD enzyme family. (A) SlASATs shown in a phylogenetic tree with all other BAHDs in cultivated tomato genome. BAHD is an acronym for four acyltransferase enzymes: benzyl alcohol O-acetyltransferase (BEAT), anthocyanin O-hydroxycinnamoyltransferase (AHCT), N-hydroxycinnamoyl/benzoyltransferase (HCBT), deacetylvindoline 4-O-acetyltransferase (DAT) - the first discovered members of this family. Please refer to Figure 1 for the biochemical activities of SlASATs. Only sequences >200aa in length were used in this analysis. Tree was constructed using Neighbor Joining with the JTT model and 100 bootstrap replicates.

**Figure 1- Figure Supplement 2:**
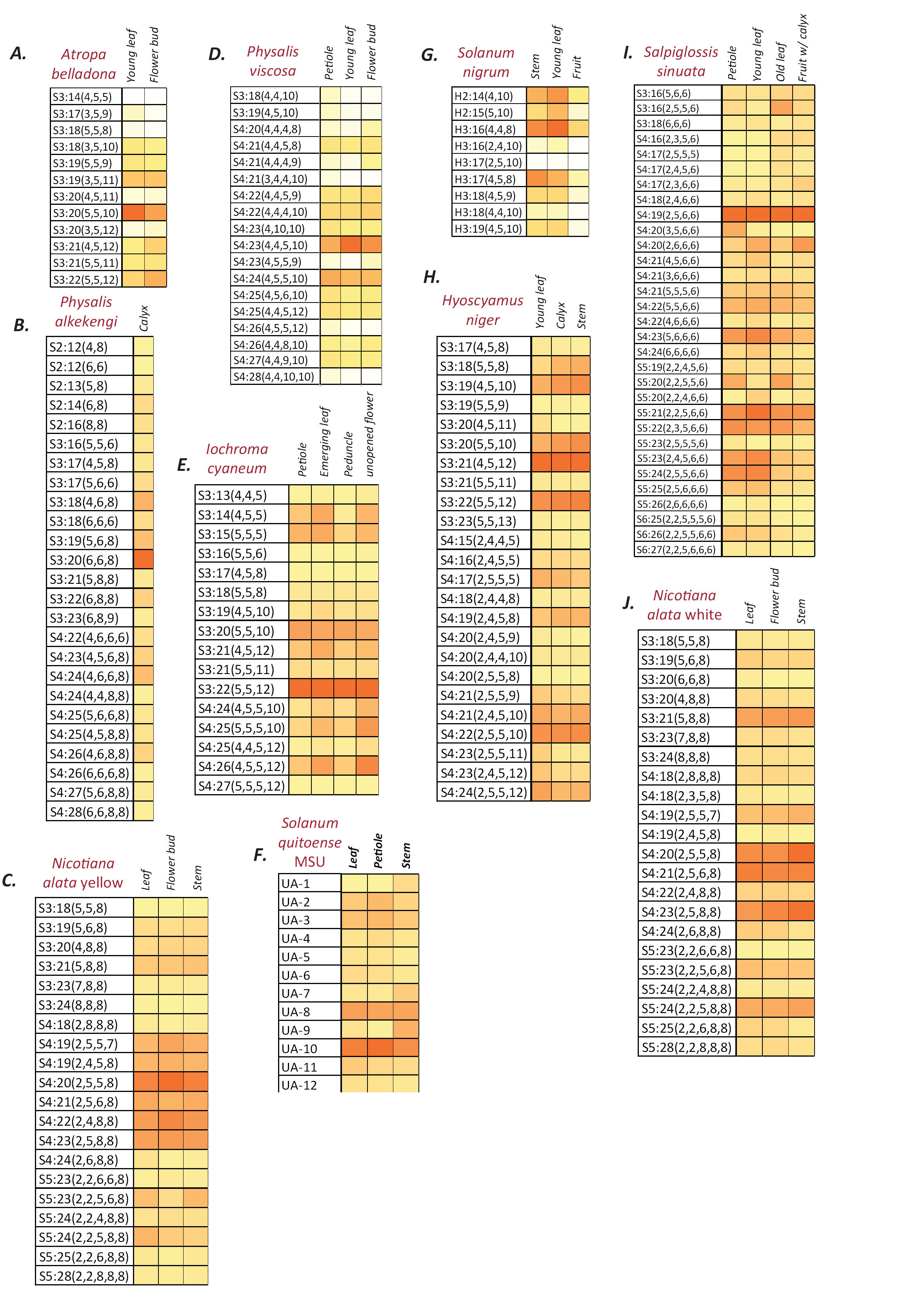
Normalized and integrated acylsugar peak areas in different species. (A-J): Peak area of each acylsugar was normalized using the internal standard peak area and dry weight, and were summed across different isomers of the same acylsugar. Colors represent % normalized peak area of a given acylsugar compared to the total normalized acylsugar peak area in the given tissue. The color scale ranges from no acylsugars (white square) to the species-specific maximum peak area (orange square). The described acylsugars are only a subset of all the detectable acylsugar-like peaks in the plant extract. UA: Unidentified acylsugar, H:Hexose

**Figure 1- Figure Supplement 3:**
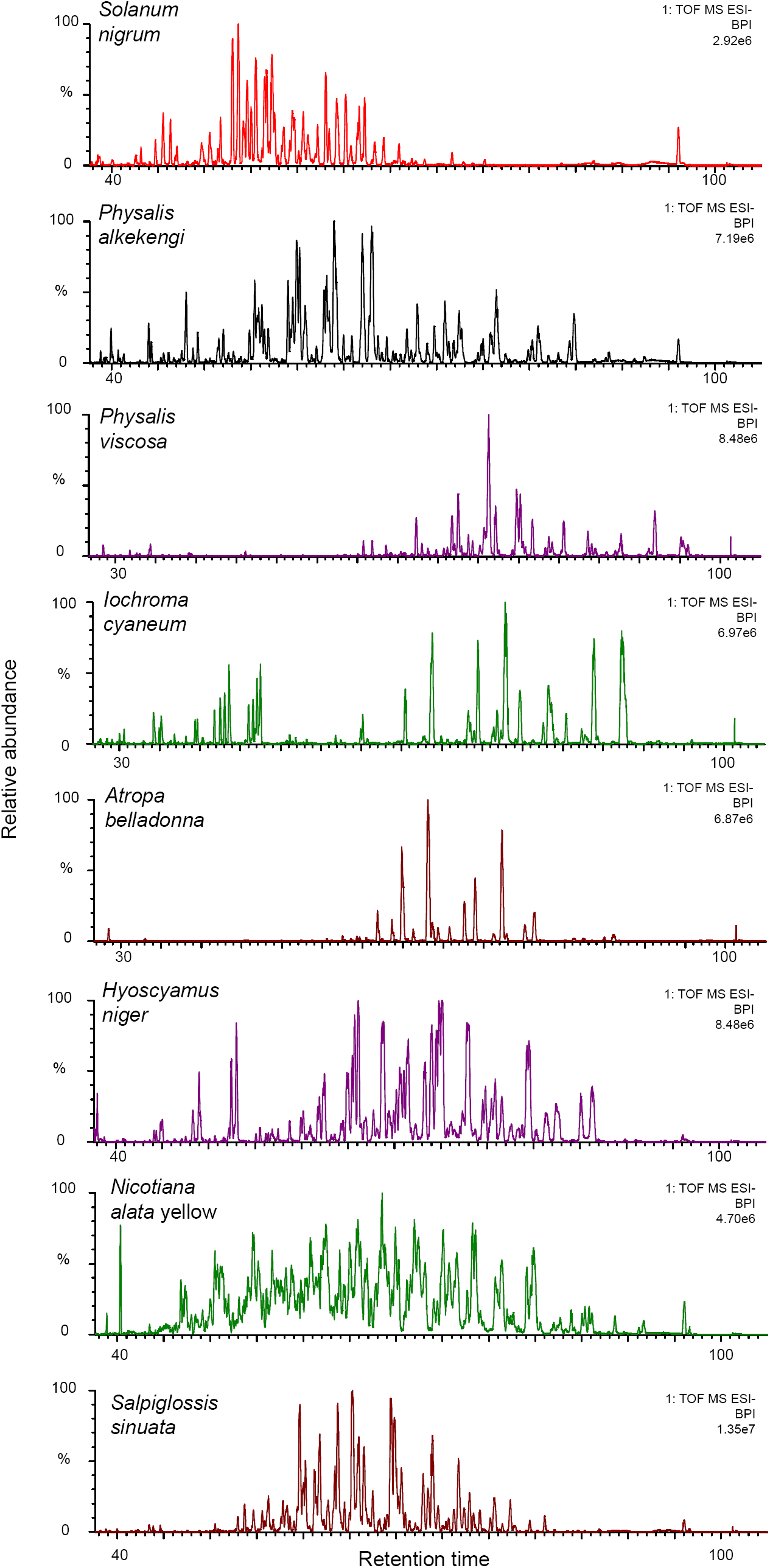
The complexity of acylsugar phenotype across multiple acylsugar producing species. Undiluted trichome extracts collected from plants at the NYBG were run on a 110-min gradient on a C18 column as described in the Methods (Supplemental Table 1). Peaks consistent with acylsugar masses and fragmentation patterns eluted between 30/40-min and 100-min for all species. Most of the peaks shown in chromatograms above have *m/z* and fragmentation patterns consistent with being acylsugars with aliphatic acyl chains. The most abundant peaks were identified, and have been described in Figure Supplement 2.

**Figure 1- Figure Supplement 4:**
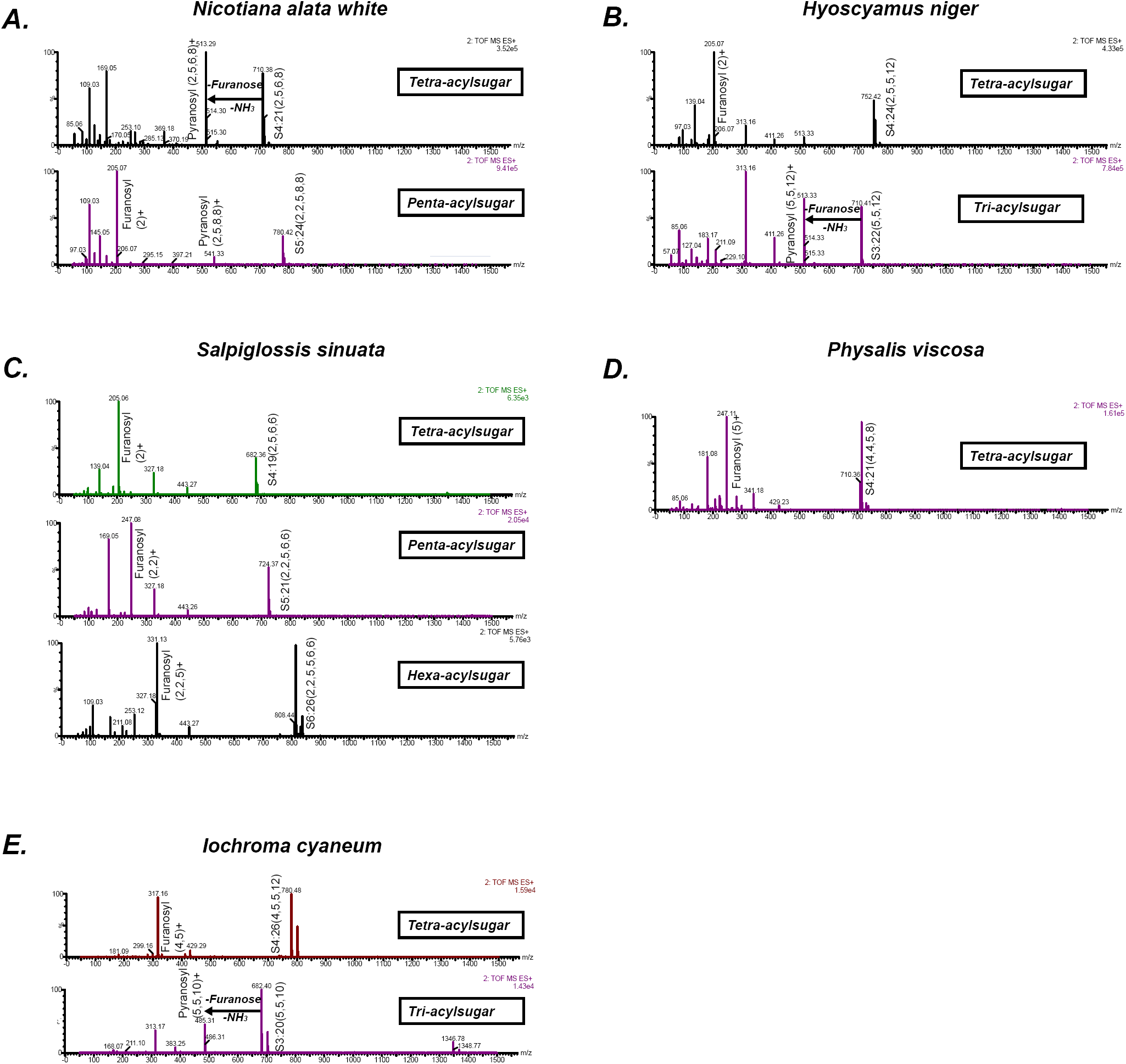
Positive mode CID mass spectra of select representative acylsugars from five species. (A-E) Each spectrum was generated at elevated collision energy to generate fragment ions. Selected fragment ions obtained at the same retention time as the assigned pseudomolecular ion of the acylsugar are annotated. The pseudomolecular ion masses are masses of the adducts of the depicted acylsugars ([M+NH4]+). The mass difference between the pseudomolecular ion (intact acylsugar) and the fragment ion provides information about the number of acyl carbon atoms on each of the pyranose and furanose rings. We interpreted the losses and the fragment ions as involving the pyranose vs. furanose ring based on knowledge about their fragmentation patterns derived from previous studies.

**Table 1-Figure Supplement 1:**
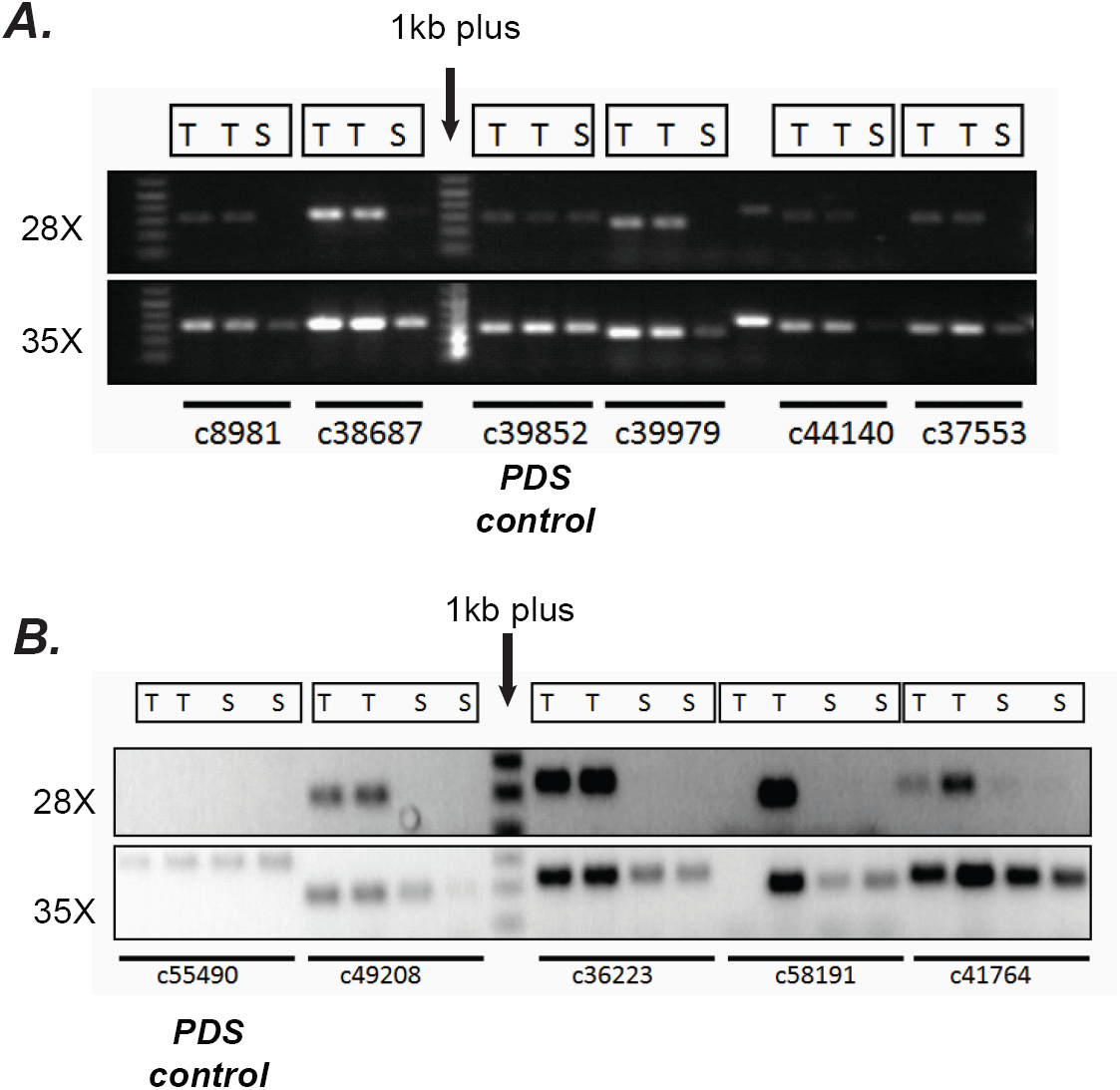
Confirmation of differential expression results from RNA-seq using semi-quantitative RT-PCR. (A, B) Verification of trichome high expression of certain transcripts using semi-quantitative RT-PCR in *S.quitoense* and Salpiglossis (B). The comparisons between Trichome (T) and Shaved stem (S) tissues involved using the same amount of starting cDNA, stopping the reactions after 28 or 35 cycles of PCR and running them on an agarose gel. Phytoene desaturase (PDS) which was not estimated to be differentially expressed based on RSEM/EBSeq analysis was used as control.

**Figure 2- Table Supplement 1:**
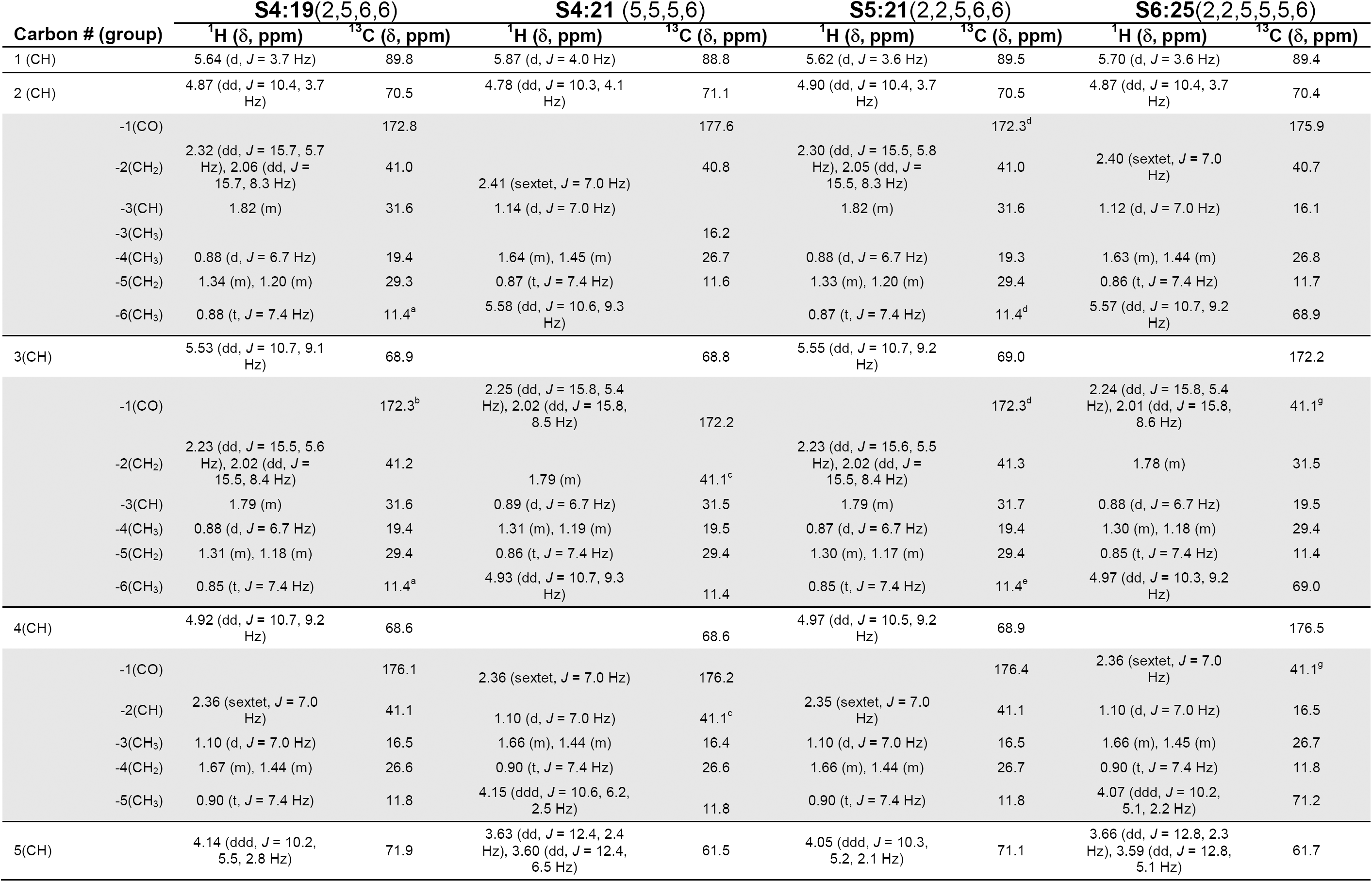
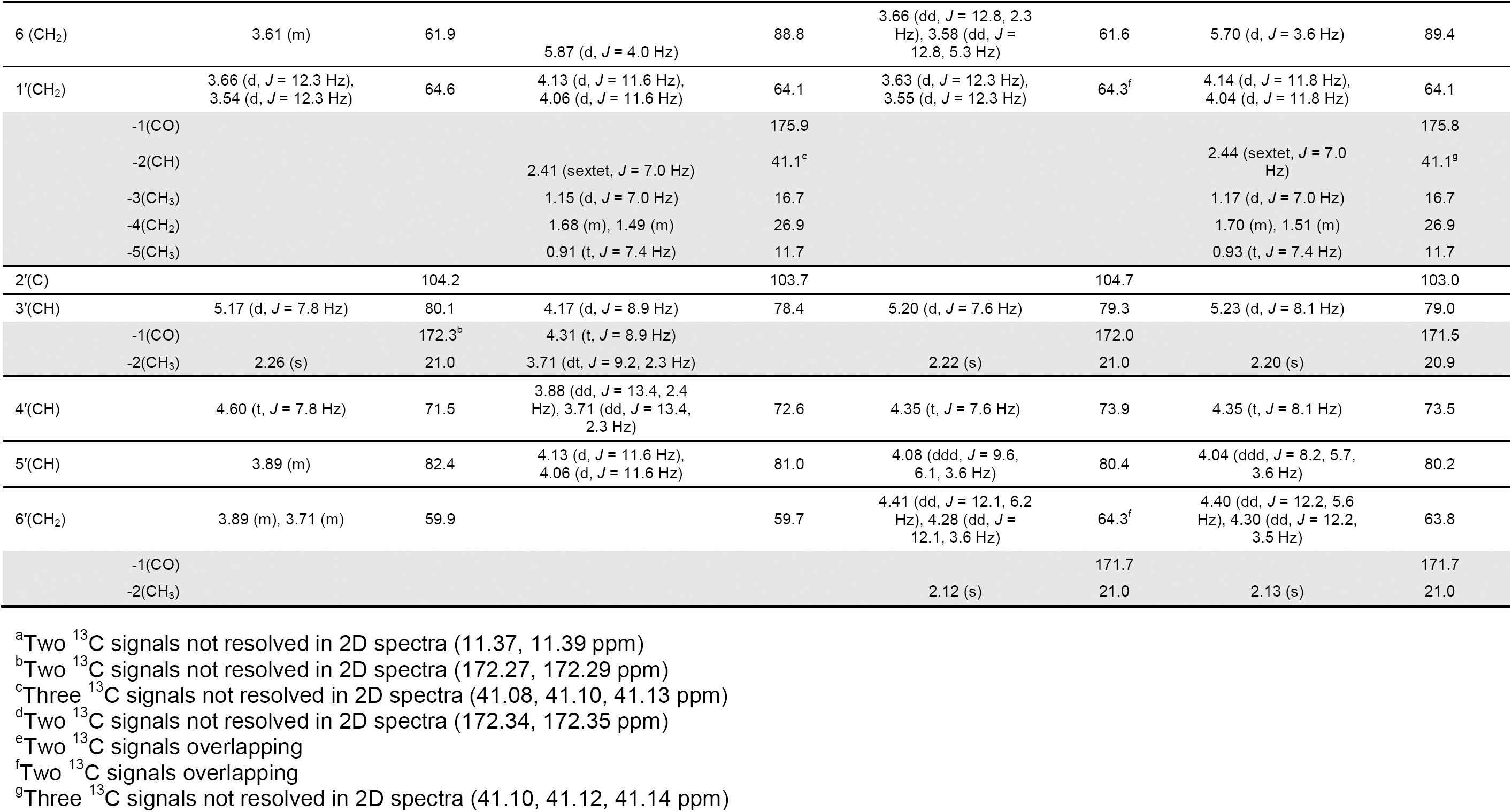
NMR chemical shifts for four acylsugars purified from Salpiglossis plants.

**Figure 2-Figure Supplement 1:**
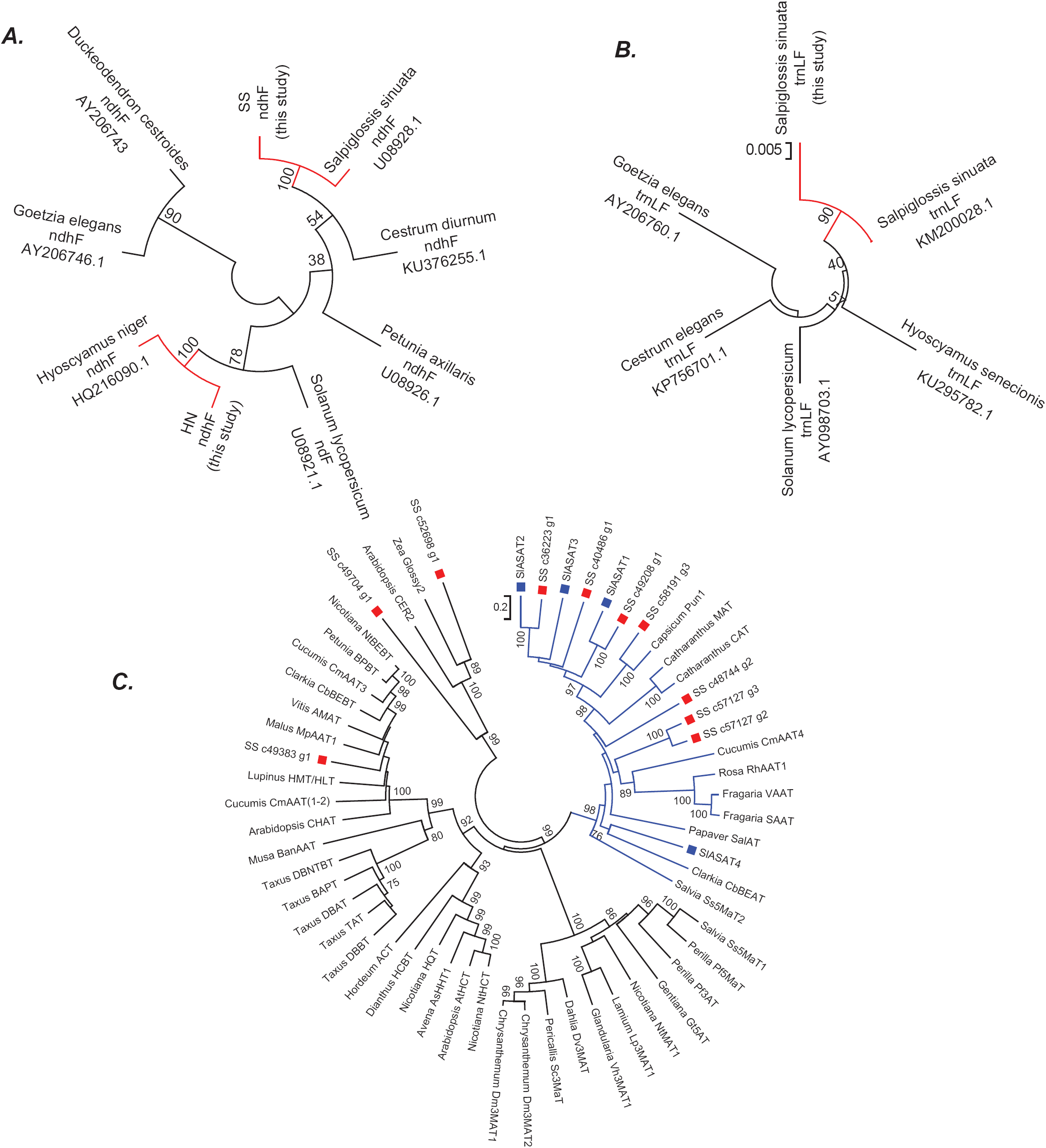
Phylogenetic positions of Salpiglossis, Hyoscyamus and Salpiglossis candidate enzymes. (A,B) Phylogeny based on the ndhF (A) and trnLF spacer (B) sequence amplified from Salpiglossis and Hyoscyamus DNA (this study) and other sequences downloaded from NCBI. Accession numbers of NCBI sequences are noted. Tree was constructed using NJ approach with 1000 bootstrap replicates. (C) Neighbor joining tree obtained using default parameters in MEGA6, with BAHD enzymes described in D’Auria (2006), SlASAT protein sequences (blue squares) and protein sequences of candidate Salpiglossis enzymes (red squares). 1000 bootstraps were performed, and all sites with <70% coverage were discarded. Only bootstrap values >70 are displayed in the figure for ease of representation. Nodes without any values have <70 bootstrap support. The blue clade represents Clade III.

**Figure 2-Table Supplement 2:**
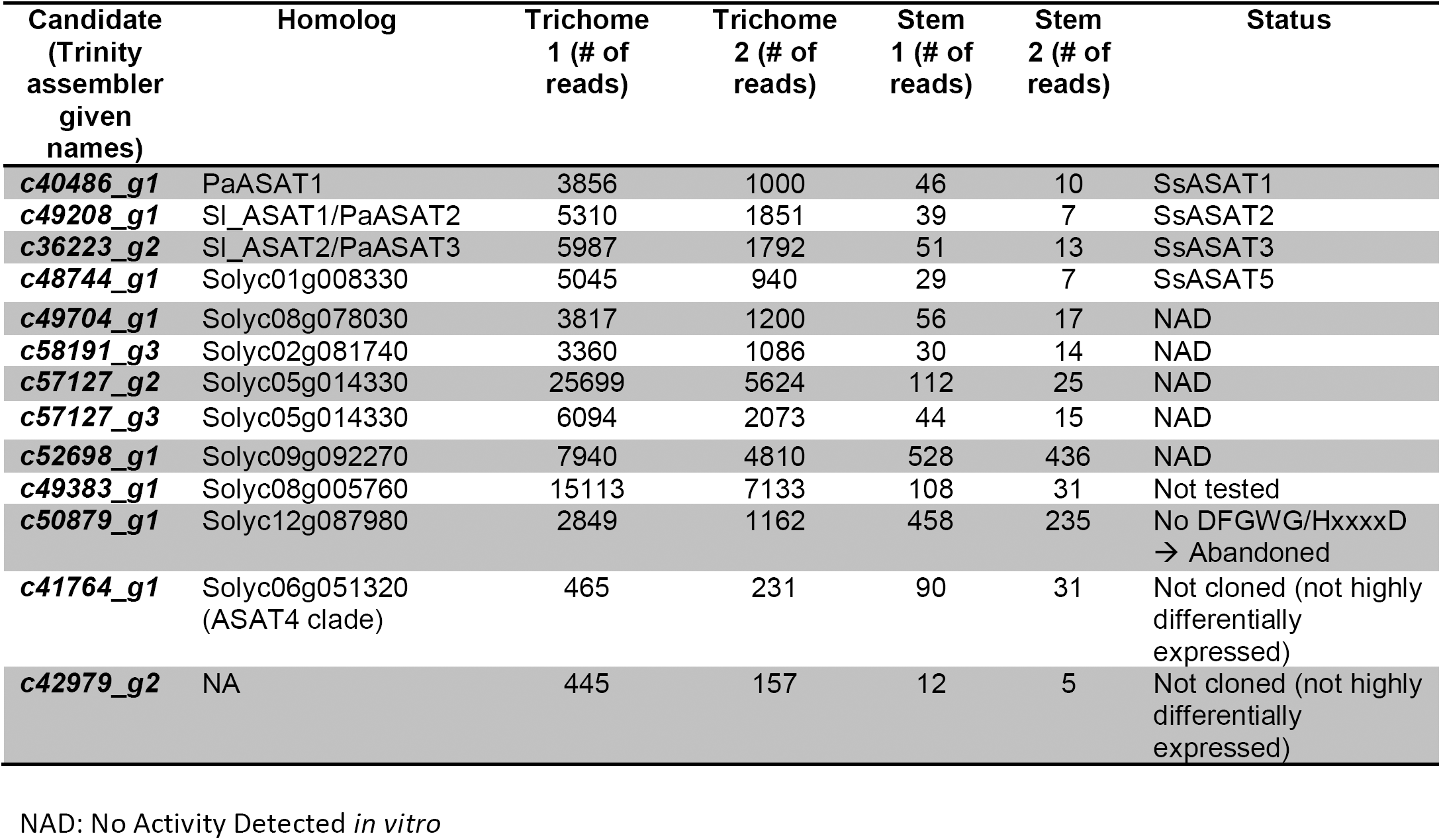
Trichome preferentially expressed BAHD enzymes.

**Figure 2-File Supplement 1.**
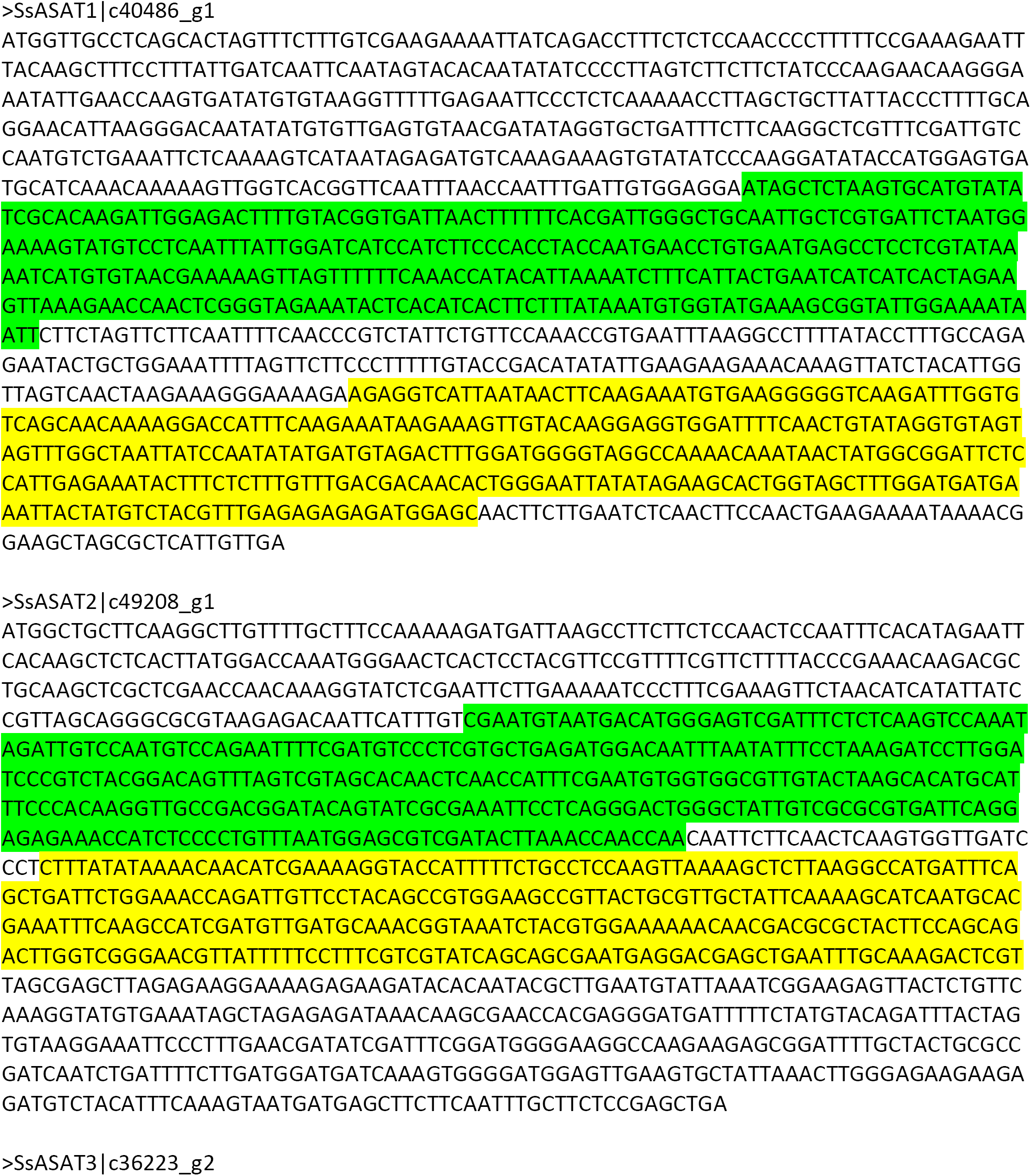
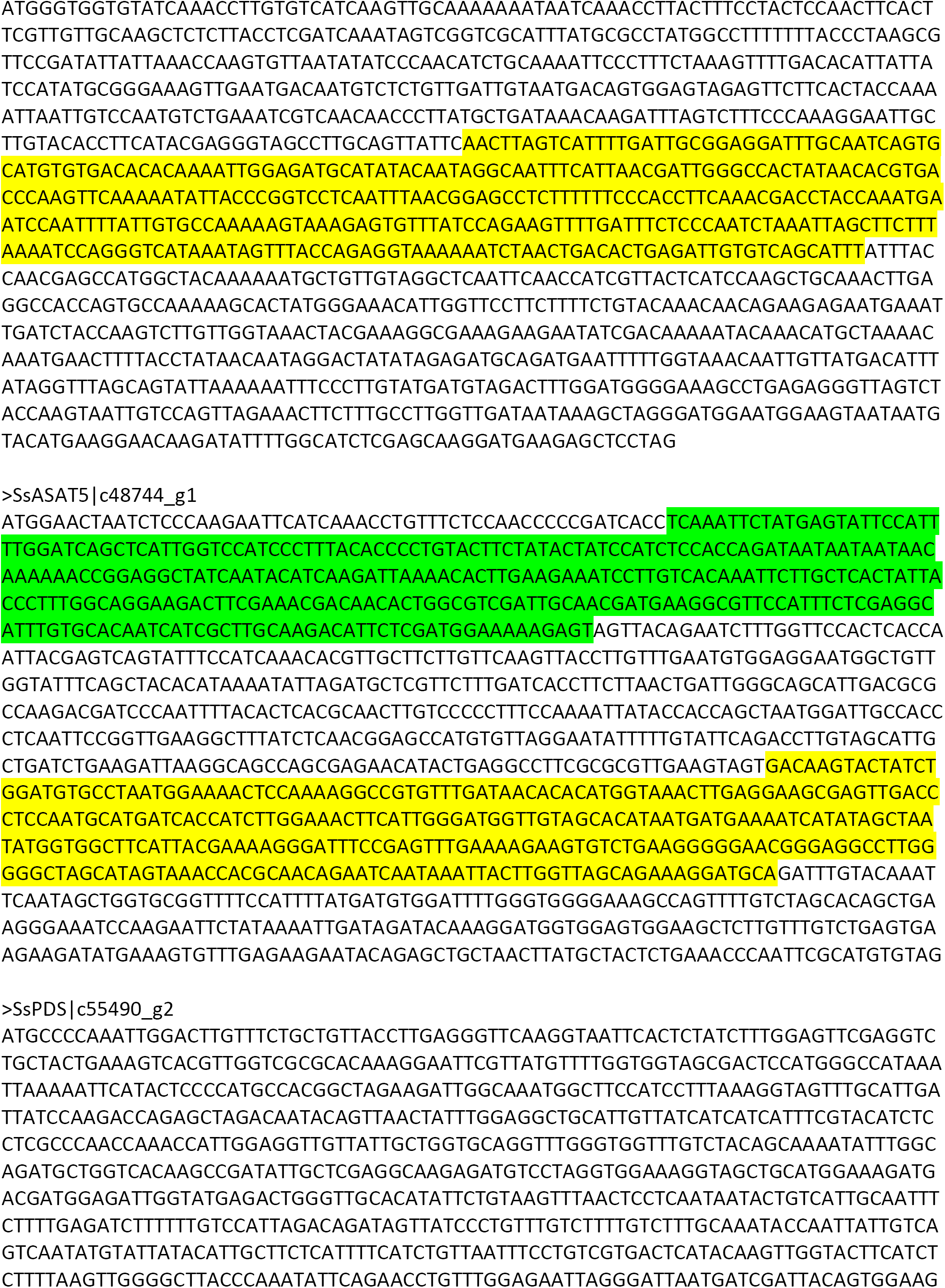
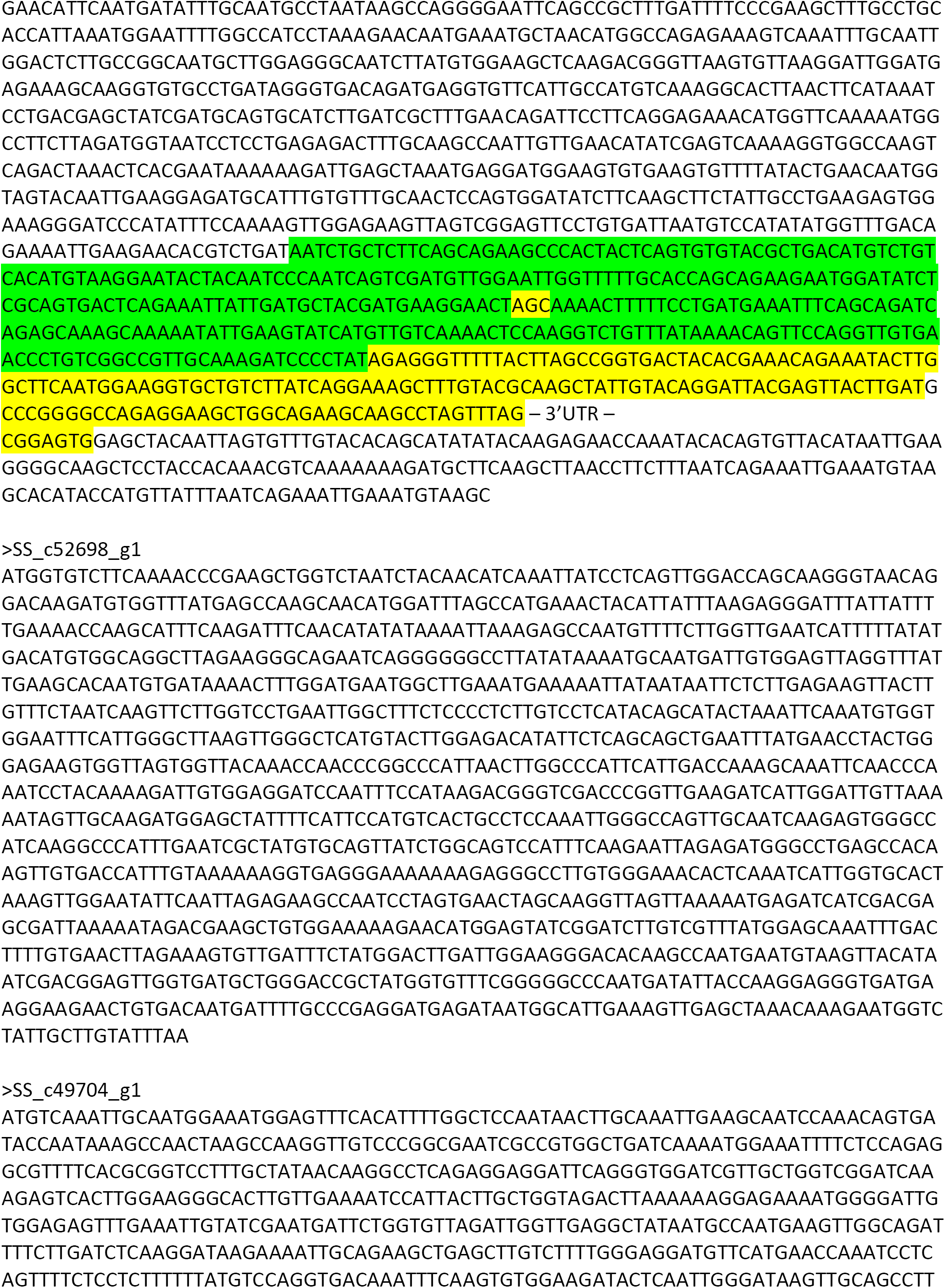
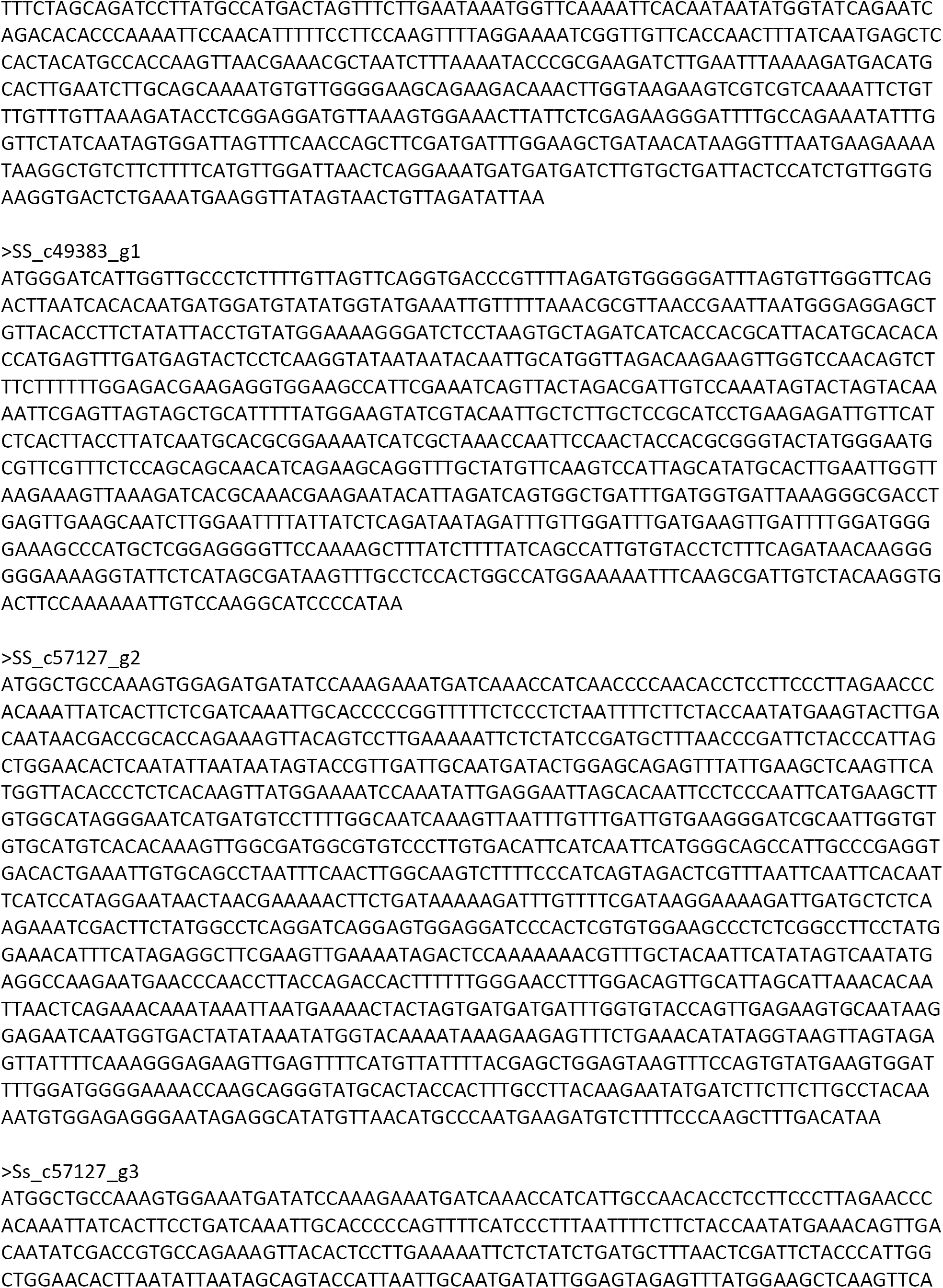
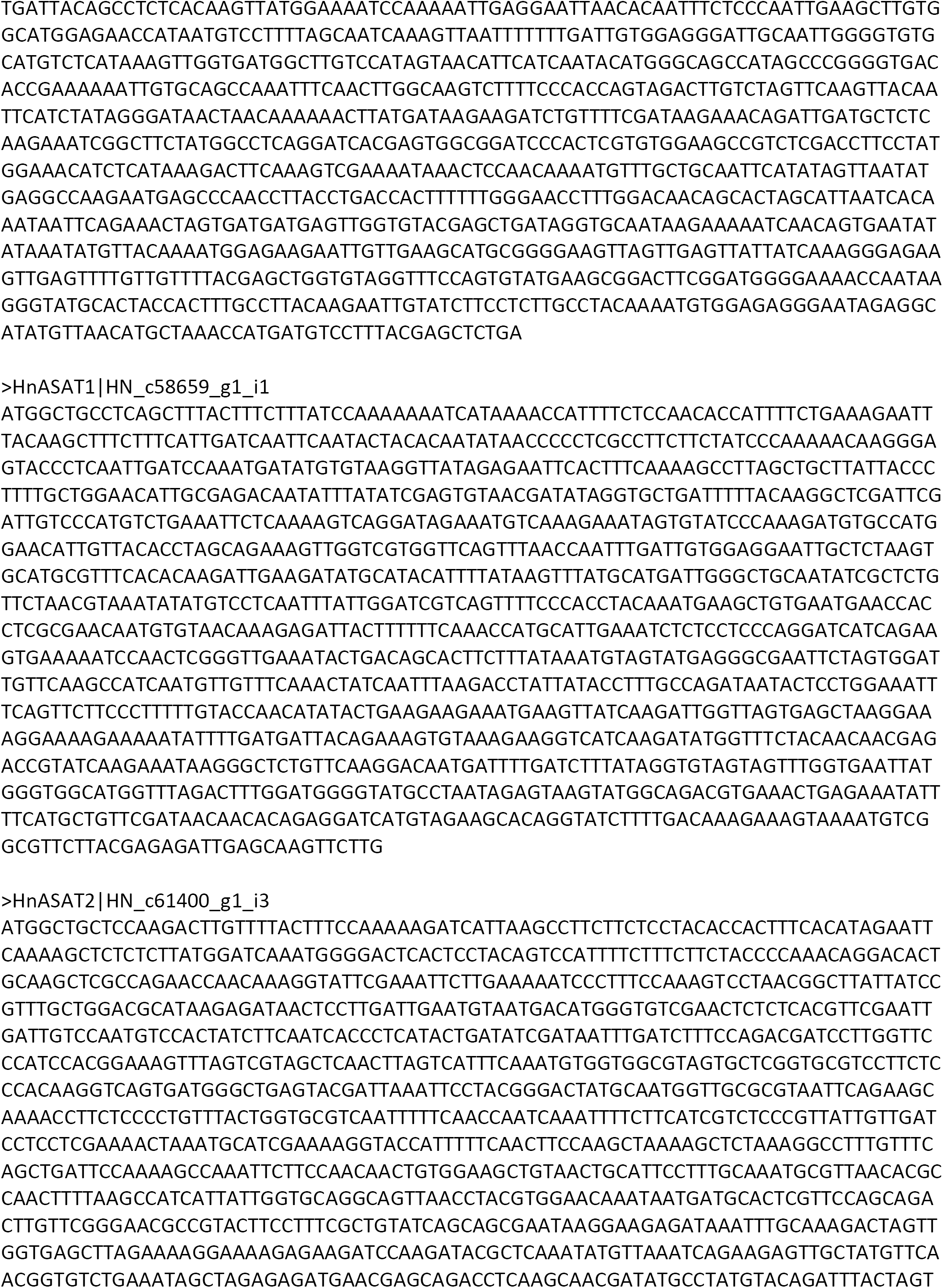
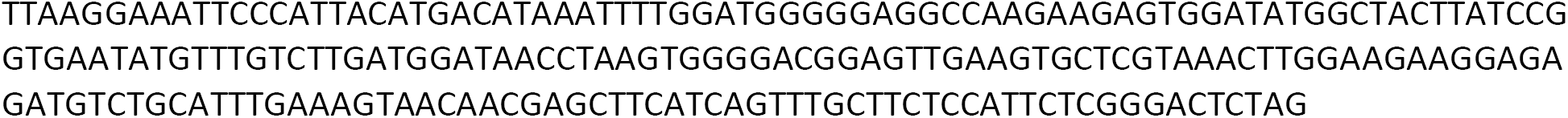
Sequences identified in this study. The transcript identifier as per the assembler Trinity nomenclature is noted in the part after the bar (|). Green and yellow highlighted sequences are the two regions targeted for VIGS in Salpiglossis.

**Figure 2-File Supplement 2:**
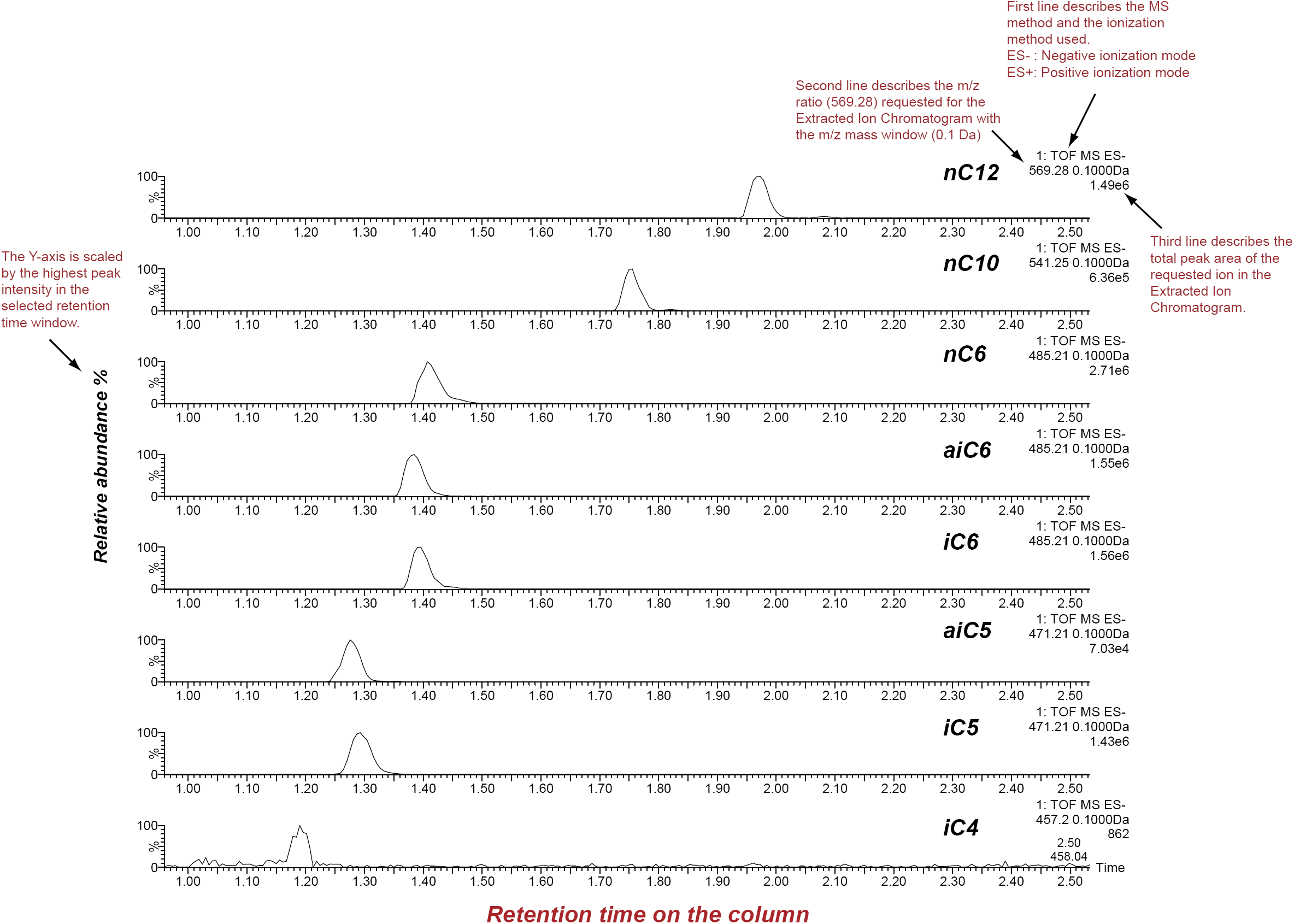
SsASAT1 reactions with different acyl CoA substrates. Shown are extracted ion chromatograms of SsASAT1 reactions using sucrose as the substrate and various acyl CoAs as donors. No product was found with acetyl CoA as donor. LC/MS analyses were conducted using a C18 column in the negative ionization mode as described in the Methods. Descriptions of different parts of the chromatogram that aid in understanding the chromatogram are highlighted in red.

**Figure 2-Figure Supplement 3:**
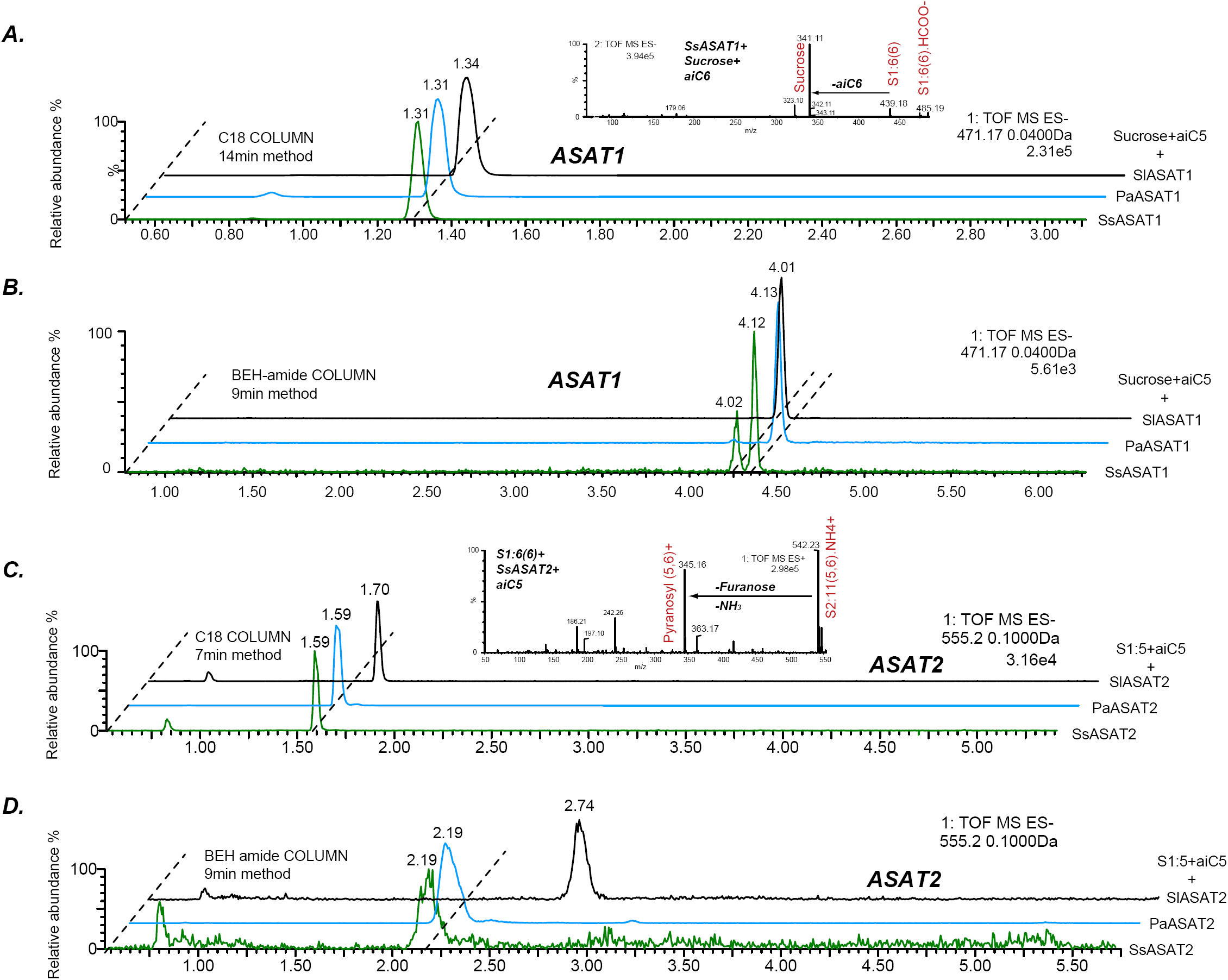
Comparative analyses of LC/MS retention times of enzyme reaction products. (A,B) Comparison between S1:5(5) produced by SsASAT1, PaASAT1 and SlASAT1 on the C18 (A) and the BEH amide (B) column. The dashed line connect the peaks aligned based on their retention times, and shows that the SsASAT1 product peaks align precisely with the PaASAT1 product peaks with two different chromatographic methods (A,B), suggesting structural identity. (C,D) Comparisons between SsASAT2, PaASAT2 and SlASAT2 enzymes similar to those shown above, suggest that SsASAT2 and PaASAT2 acylate the same positions on the sucrose molecule. Inset in (A) shows negative ion mode fragmentation of S1:6(6) product of the SsASAT1 reaction using aiC6 CoA as donor. Inset in (C) shows positive ion mode fragmentation of S2:11(5,6) product of SsASAT2 using S1:6(6) produced by SsASAT1 as substrate. Inset in (C) shows that both aiC5 and aiC6 are added on the same (pyranose) ring.

**Figure 2-Figure Supplement 4:**
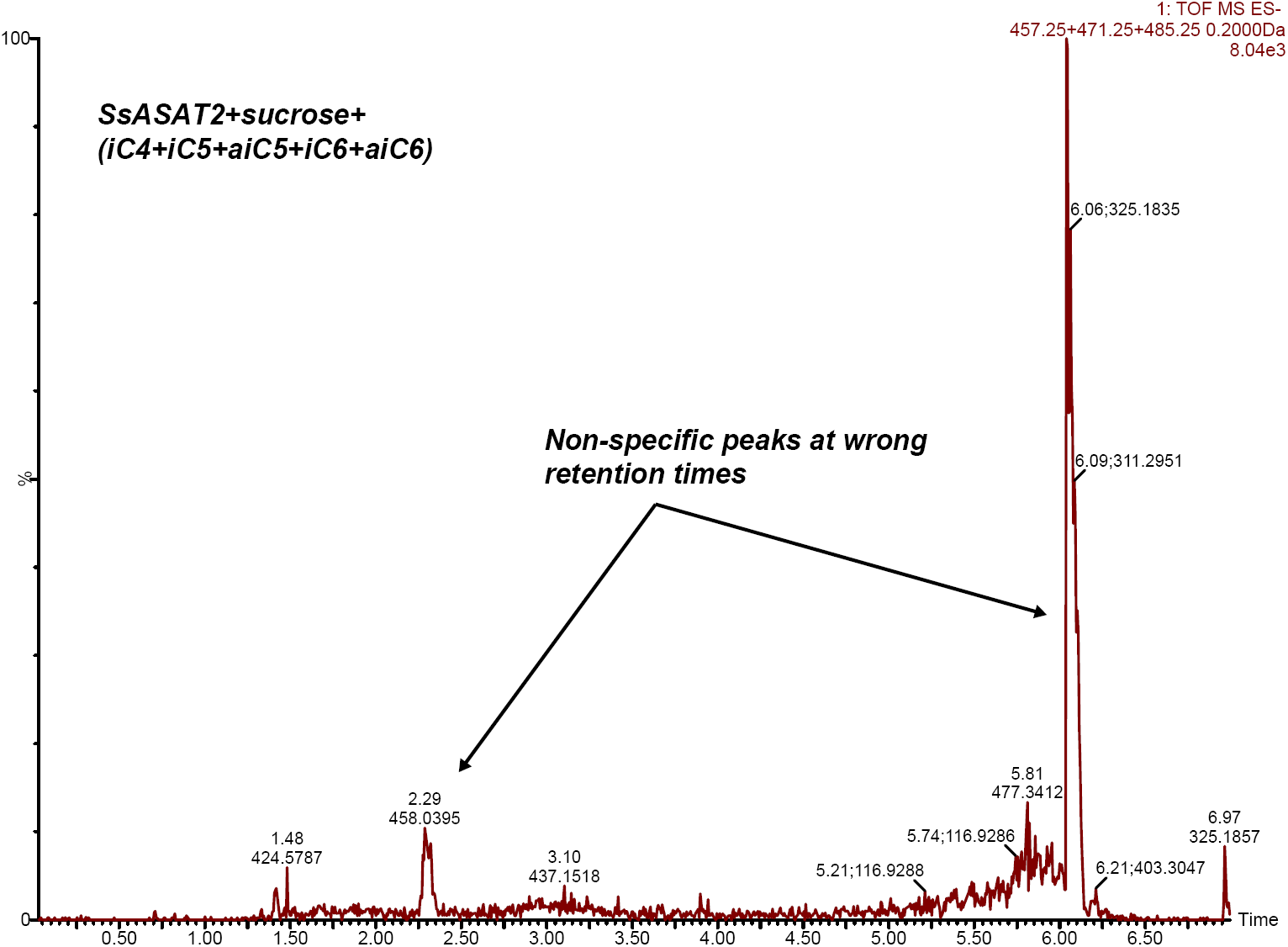
SsASAT2 does not acylate sucrose. Extracted ion chromatogram of masses corresponding to S1:4(4), S1:5(5) and S1:6(6). No peaks corresponding to these products are seen. Additional reactions with individual CoAs -- aiC6 CoA and nC12 CoA -- are shown in Figure 7-Figure Supplement 1.

**Figure 2-Figure Supplement 5:**
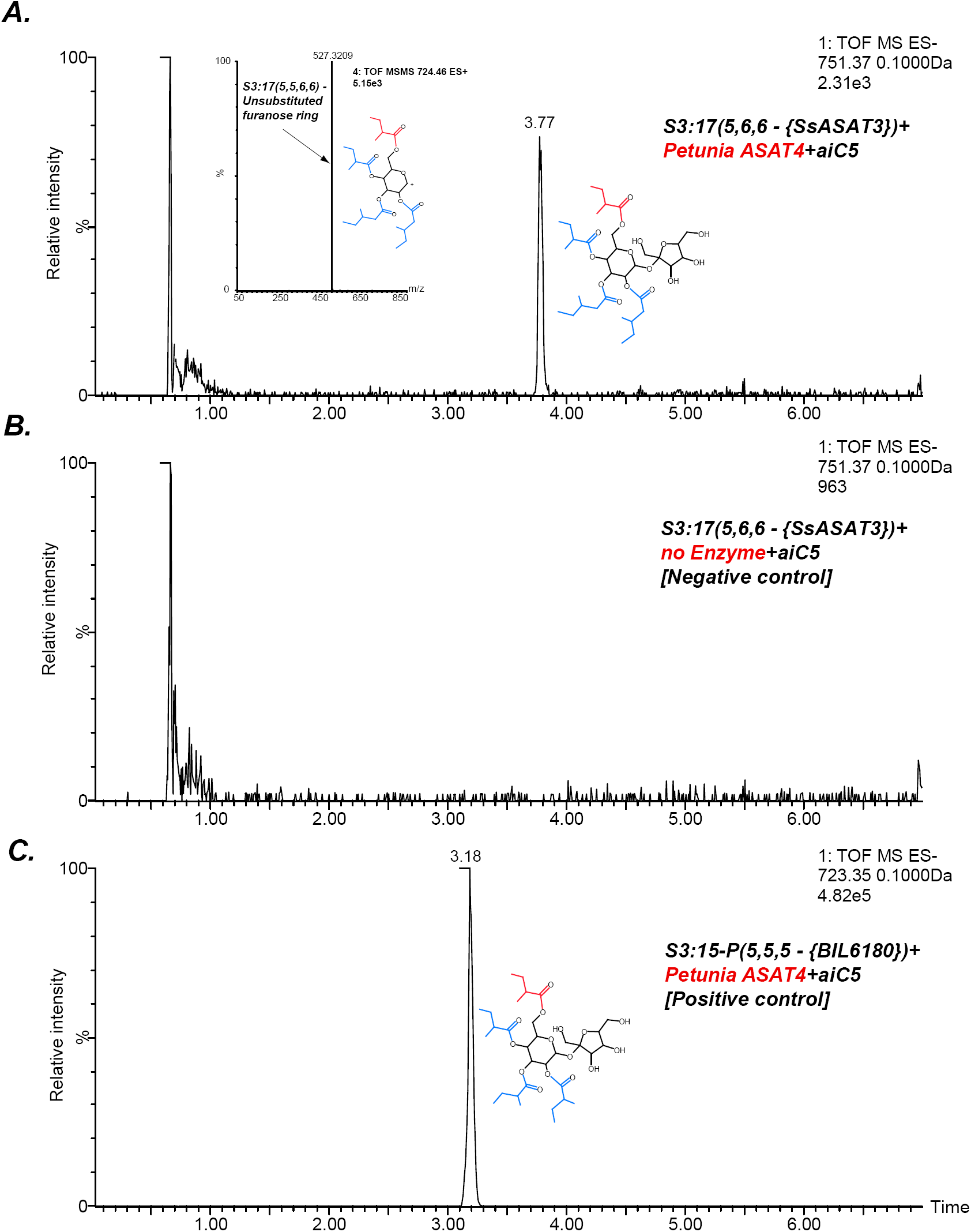
SsASAT3 acylates at the R3 position. An indirect assessment of SsASAT3 positional specificity using Petunia ASAT4, which acylates at the pyranose R6 position (A) The chromatogram shows that Petunia ASAT4, which is known to acylate at the R6 position, can acylate the tri-acylated sugar S3:17(5,6,6) produced by SsASAT3 using the SsASAT2 product S2:10(5,5). The inset shows positive mode data suggesting all four chains are on the same ring. Given R6 is the only free hydroxyl group available on the pyranose ring in S3:17(5,6,6), and since SsASAT1 and SsASAT2 acylate R2 and R4 positions, the only position SsASAT3 can acylate without inhibiting PaASAT4 is R3. (B) Negative control with no enzyme (C) Positive control with S3:15-P(5,5,5) purified from BIL6180, with R2, R3 and R4 positions substituted by C5 chains based on NMR data (unpublished data).

**Figure 2-Figure Supplement 6:**
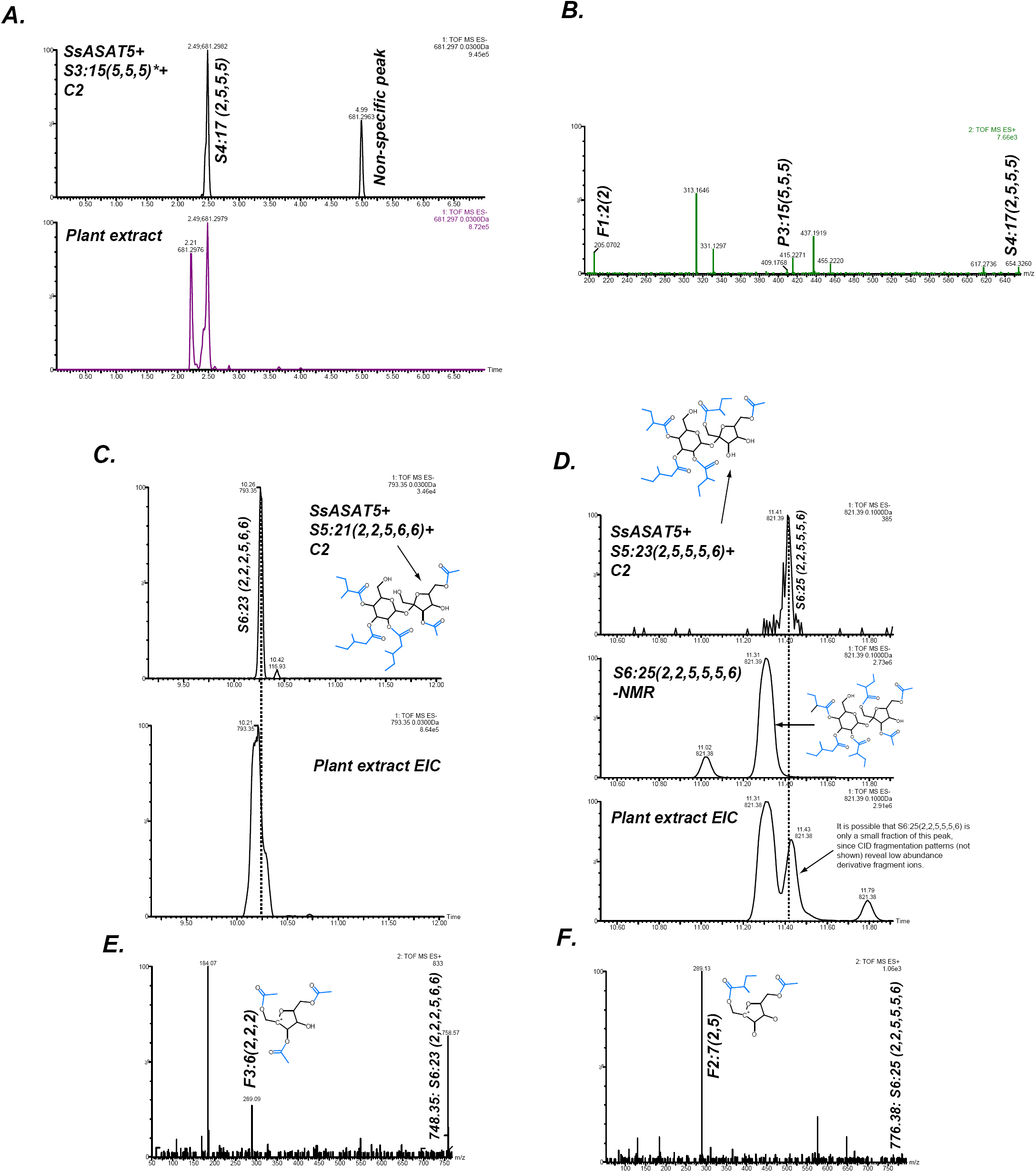
SsASAT5 putative secondary activities. (A) SsASAT5 can transfer C2 to S3:15 (5,5,5) isolated from *Solanum pennellii*-derived Backcross Inbred Line BIL6180 (unpublished data; Ofner et al, 2016), which has the three acyl chains on the R2, R3 and R4 pyranose (abbr: P) ring, same as Salpiglossis. (B) The C2 addition likely occurs on the furanose (abbr: F) ring, as seen by the presence of the m/z 205.07 in the positive-ion mode elevated collision energy mass spectrum. (C,D) SsASAT5 can transfer C2 to penta-acylated sucroses, and the product co-migrates with an acylsugar of the same molecular mass produced in the plant. Co-migrating peaks are aligned by the dashed line. Central panel in (D) shows that the S6 sugar produced by SsASAT5 is not the same as the most abundant acylsugar produced by the plant purified for NMR, which has two C2s on the furanose ring. Shown are extracted ion LC/MS chromatograms for the expected masses. (E,F) The most abundant fragment ions (m/z: 289.09, 289.13) are consistent with C2 addition occurring on the furanose and pyranose rings, respectively.

**Figure 3 – Table Supplement 1:**
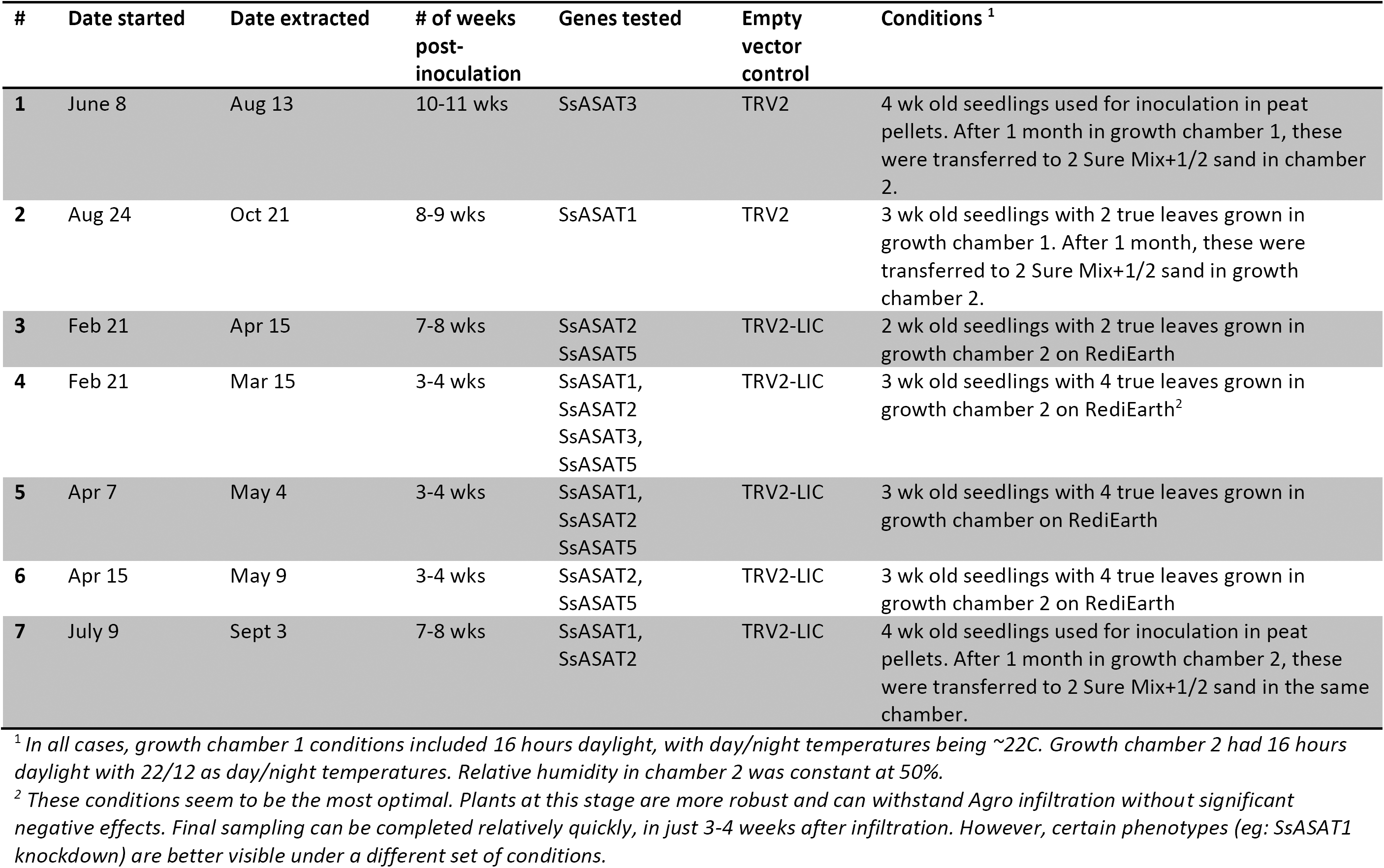
VIGS plants growth conditions.

**Figure 3-Figure Supplement 1:**
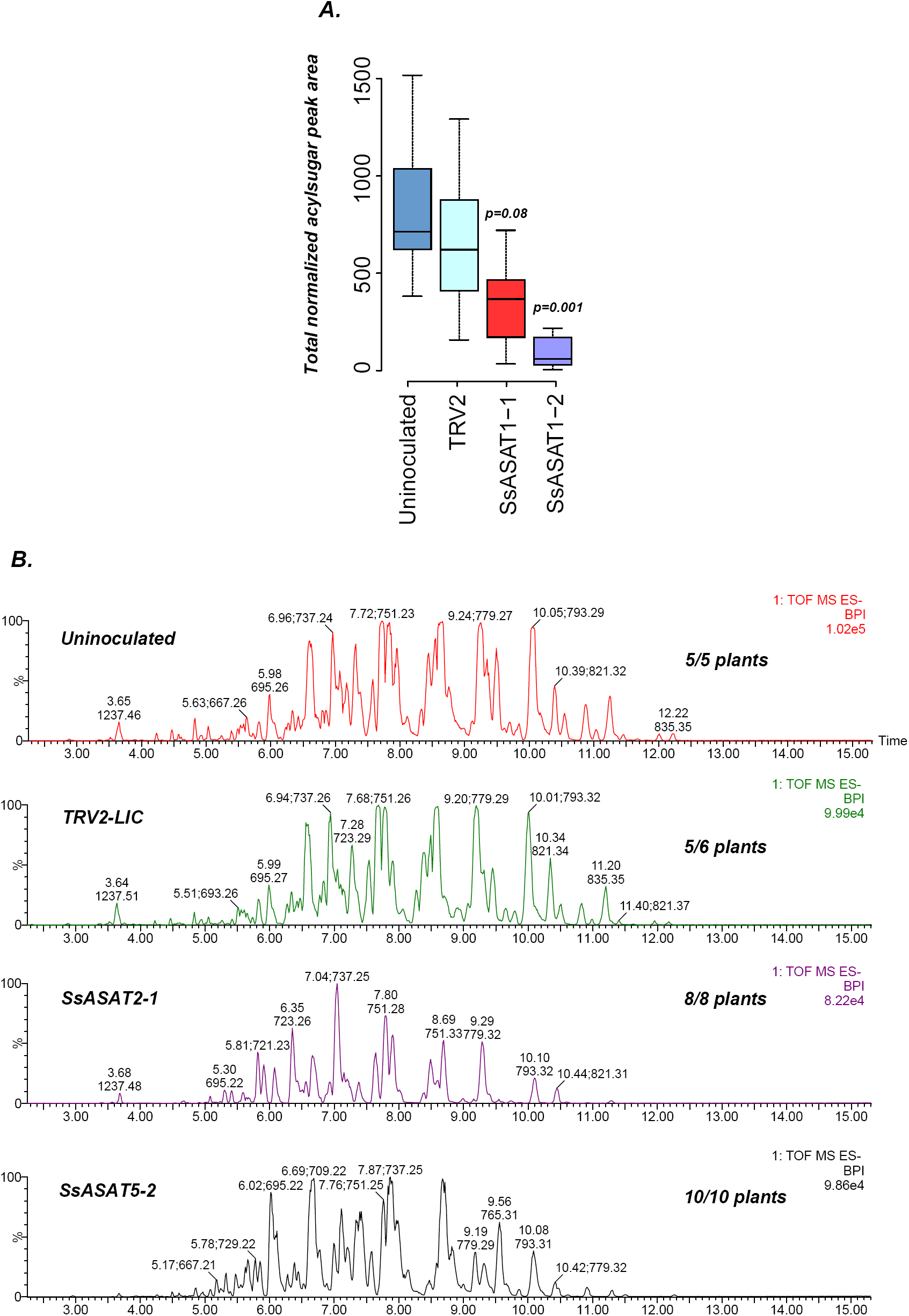
Results of knockdown of SsASAT1, SsASAT2 and SsASAT5 transcripts in a distinct replication of VIGS experiments. (A) SsASAT1 knockdown shows reduction of acylsugar levels. (B) SsASAT2 and SsASAT5 knockdown causes substantial changes in Salpiglossis acylsugar profiles. The accumulating acylsugars in these two knockdowns are different, as seen by the co-eluting *m/z* ratios. Number of plants showing depicted phenotype out of the total number of analyzed plants is noted on the right side of each chromatogram.

**Figure 3-Figure Supplement 2:**
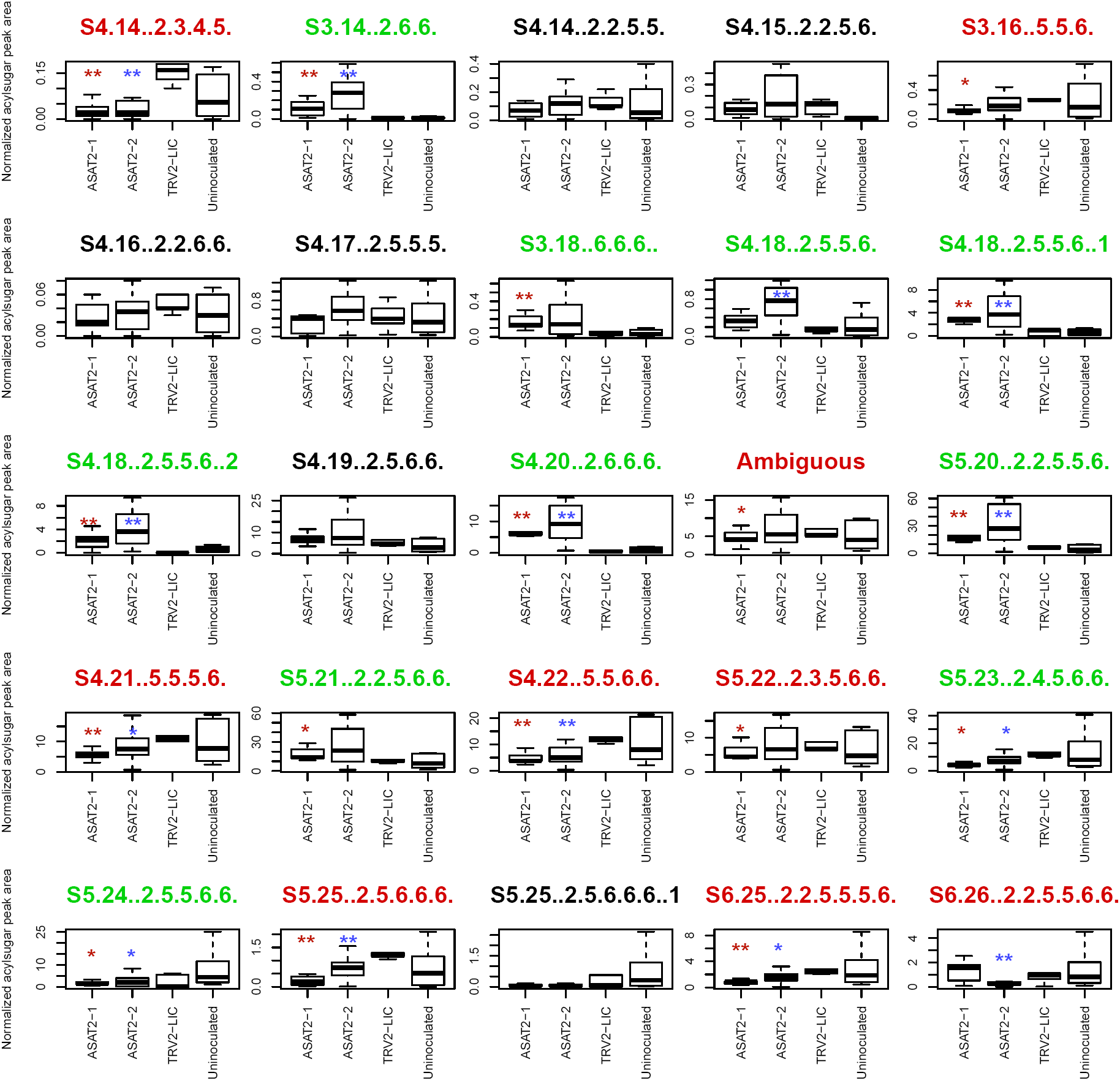
Individual acylsugar levels in SsASAT2 VIGS replicate 1. The boxplots denote the summed normalized acylsugar peak areas, obtained using the LC/MS extracted ion chromatograms, across all plants in the experiment. Blue and red asterisks indicate significant changes in SsASAT2-1_TRV2-LIC and SsASAT2-2_TRV2-LIC comparisons, respectively. *: p<0.05; **: p<0.01 using the KS test. Acylsugar names in green show significant increases and the names in red show significant decreases compared with the TRV2-LIC control. These results show that C2 acylated derivatives of S3:16(5,5,6), S3:17(5,6,6) and S3:18(6,6,6) [e.g.: S4:18(2,5,5,6); S5:20(2,2,5,5,6)] increased significantly, while C5 derivatives of these acylsugars [eg: S4:21(5,5,5,6); S5:25(2,5,6,6,6)] decreased significantly in the silenced lines. One possible explanation for this result is that another transcript was silenced in addition to SsASAT2. However, the single Salpiglossis transcript with 100% identity to the silencing fragment over 18 nucleotides is not preferentially expressed in the trichomes of uninfected Salpiglossis (∼360 reads in trichome; ∼550 reads in shaved stem) and is annotated as an ion channel. Thus, the reason for this unusual phenotype is still unclear.

**Figure 4-Figure Supplement 1:**
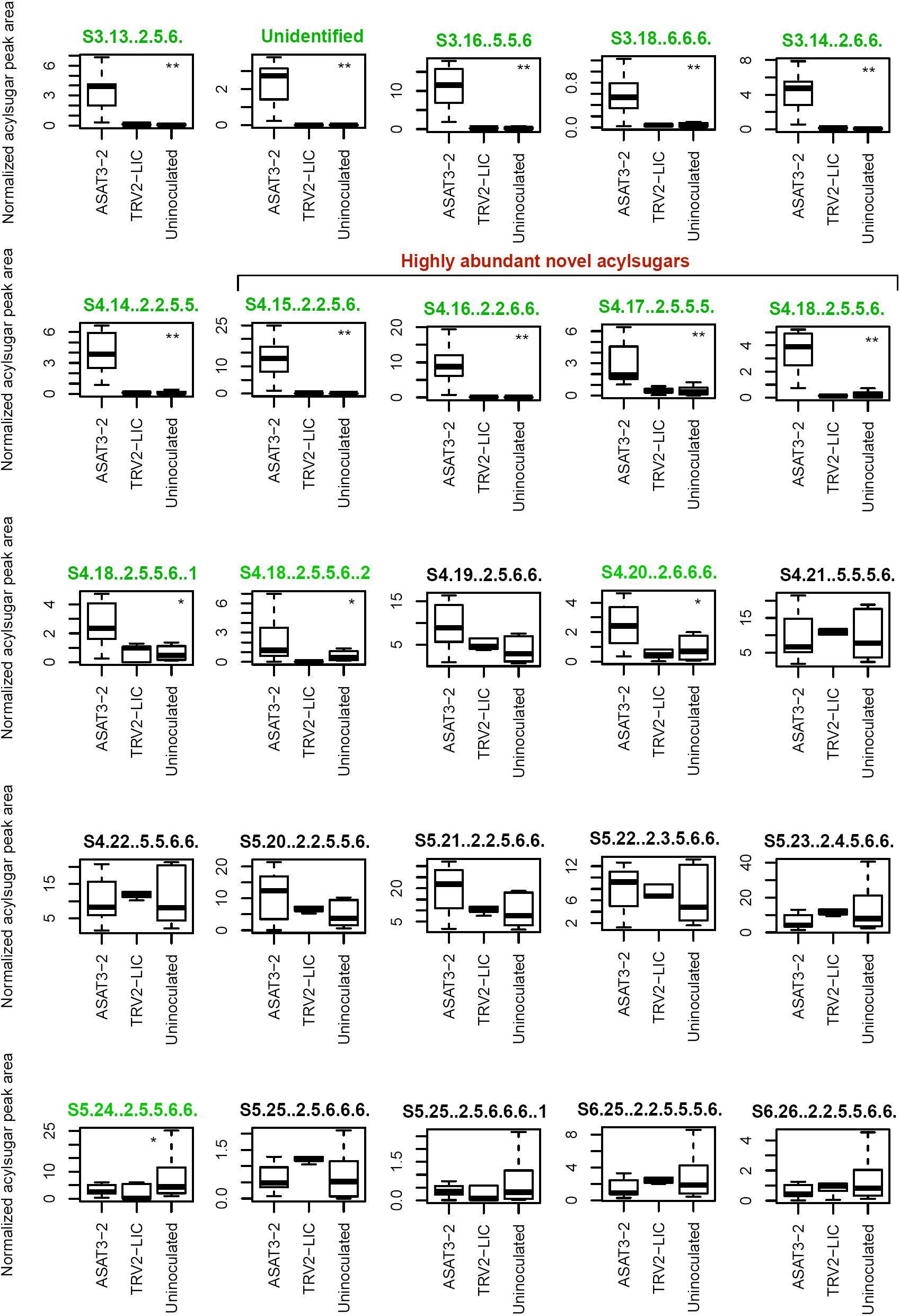
SsASAT3 knockdown boxplots for levels of individual acylsugars. Statistical significance of the difference between TRV2-LIC and ASAT3-2 was tested using Kolmogrov-Smirnov test. ** represents p<0.01 and * represents p<0.05. Boxes where the acylsugars had significantly different levels compared to TRV2-LIC are highlighted in green. Acylsugar peak area was normalized as described in the Methods.

**Figure 4- Figure Supplement 2:**
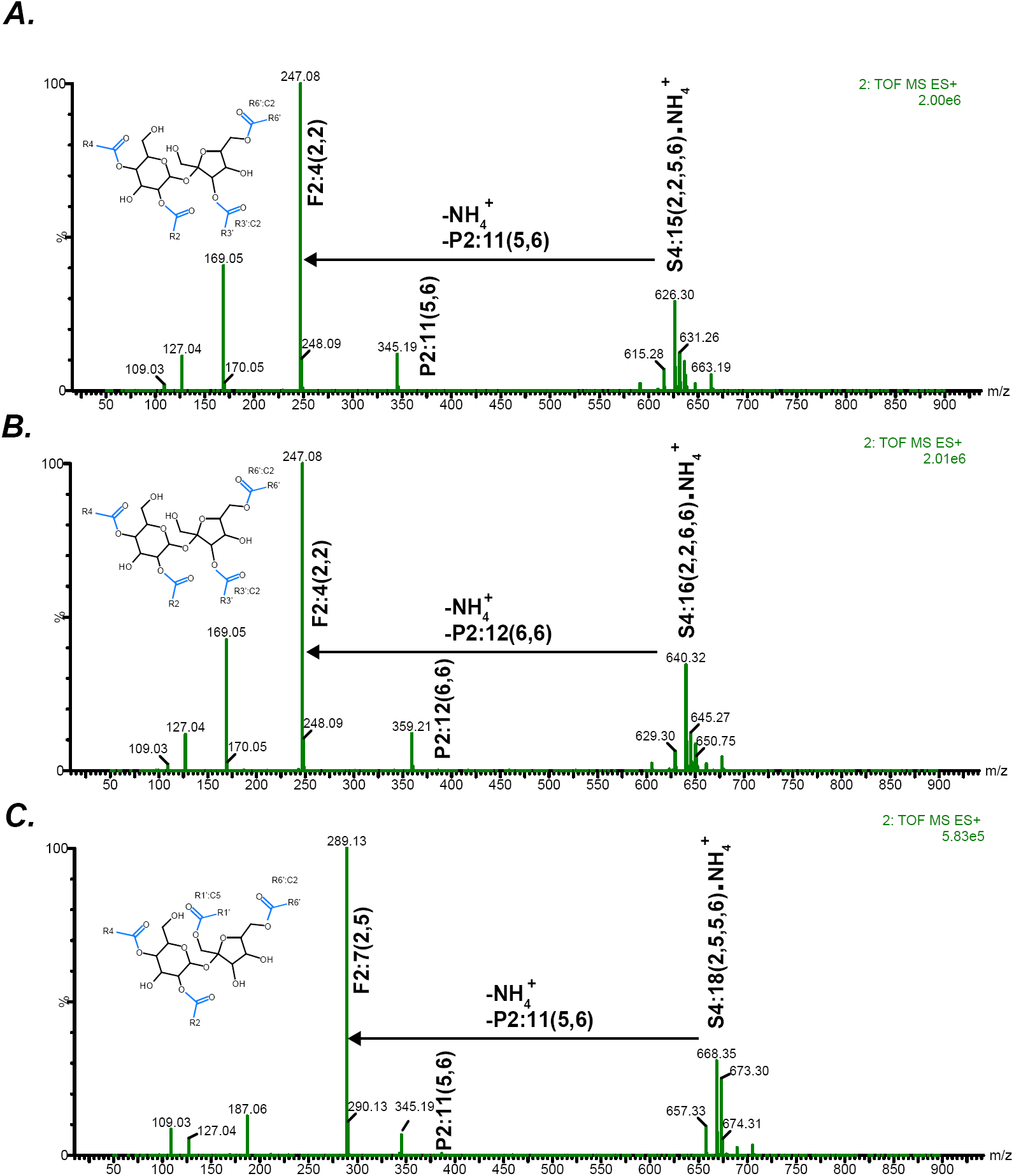
Positive mode fragmentation patterns of novel acylsugars found in SsASAT3 VIGS knockdown plants. In (A,B,C), mass spectra obtained at elevated collision energies of three different novel acylsucrose peaks are shown. We make the assumption that fragment ions and the most abundant pseudomolecular ions appearing at the same retention time are associated. In all three cases, assuming that the original mono- and di-acylations occurred on the pyranose (P) ring, we can infer that the subsequent C2 and C5 additions giving rise to the tri- and tetra-acylated sugars occurred on the furanose (F) ring.

**Figure 4- Figure Supplement 3:**
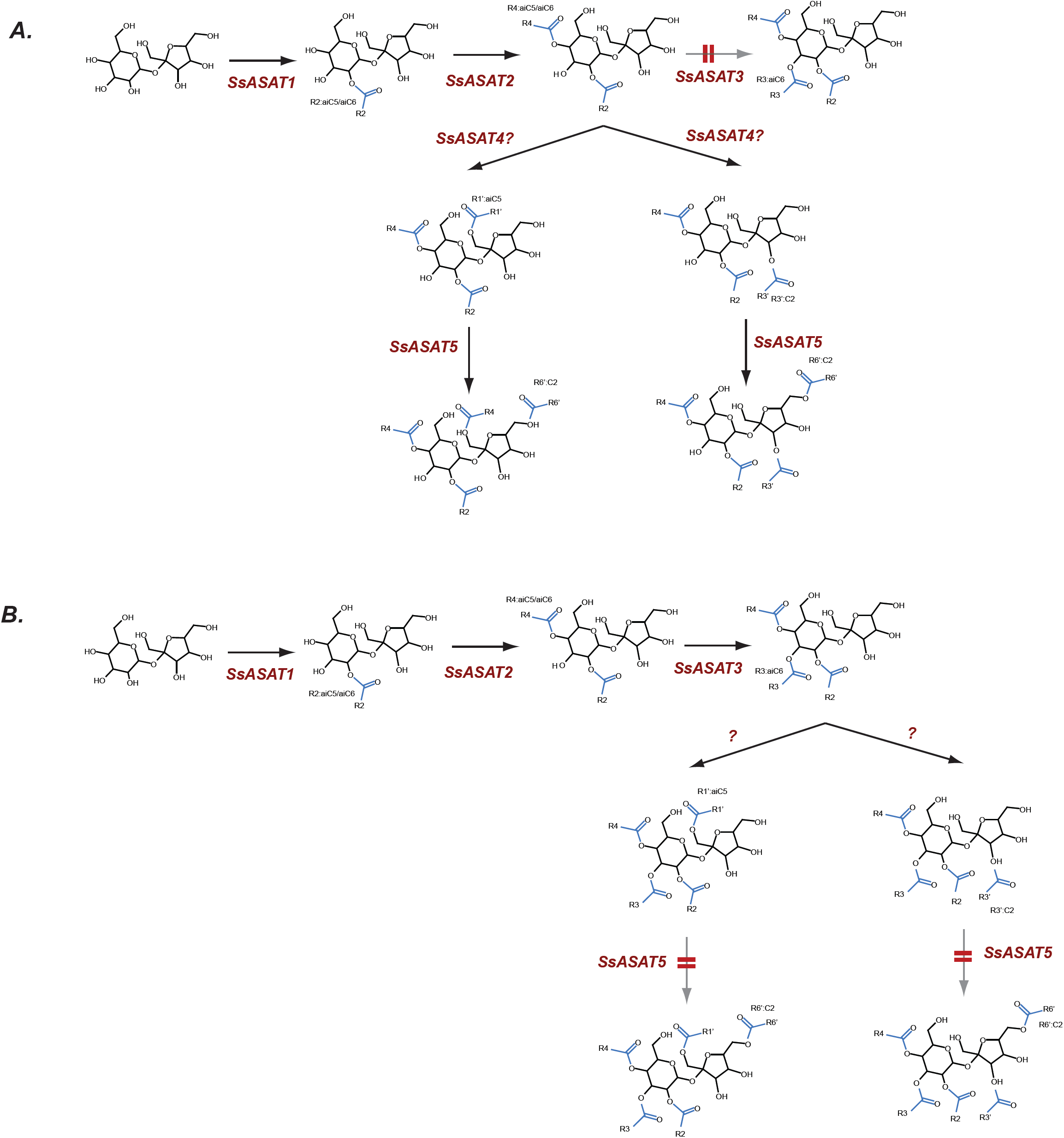
Hypothesized routes of metabolite flow in VIGS knockdown plants. (A) SsASAT3 knockdown and (B) SsASAT5 knockdown. The perturbed pathway steps are denoted by a blurred arrow and two red lines. All acylation positions are hypothesized based on *in vitro* data and knowledge of the enzyme activities

**Figure 6- Figure Supplement 1:**
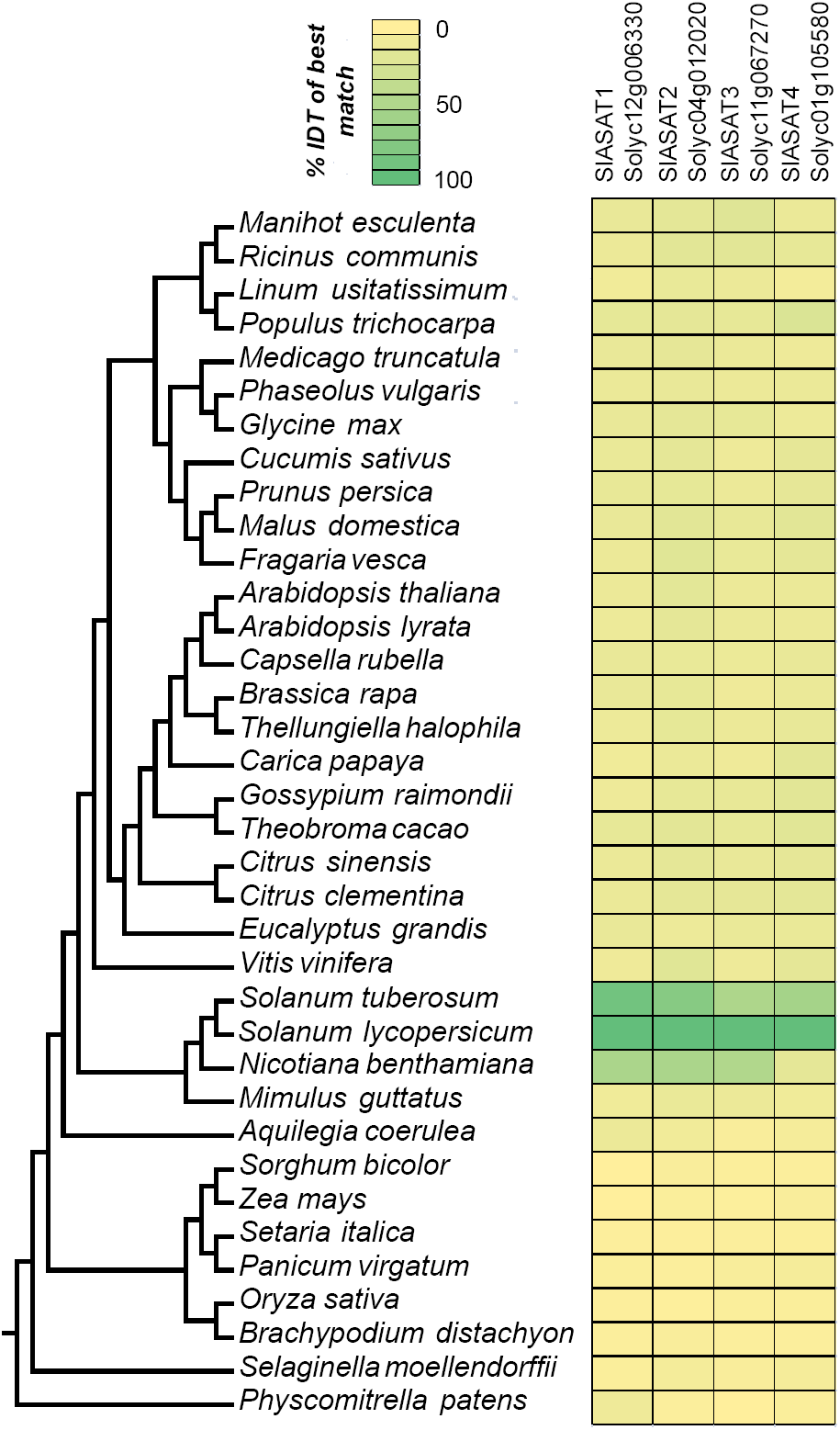
The phylogenetic context of tomato ASATs. Results of protein BLAST (BLASTP) between SlASAT sequences and proteomes of each species in the Phytozome database. The % identity (% IDT) values for the top BLASTP hits in each species are shown.

**Figure 6-Figure Supplement 2:**
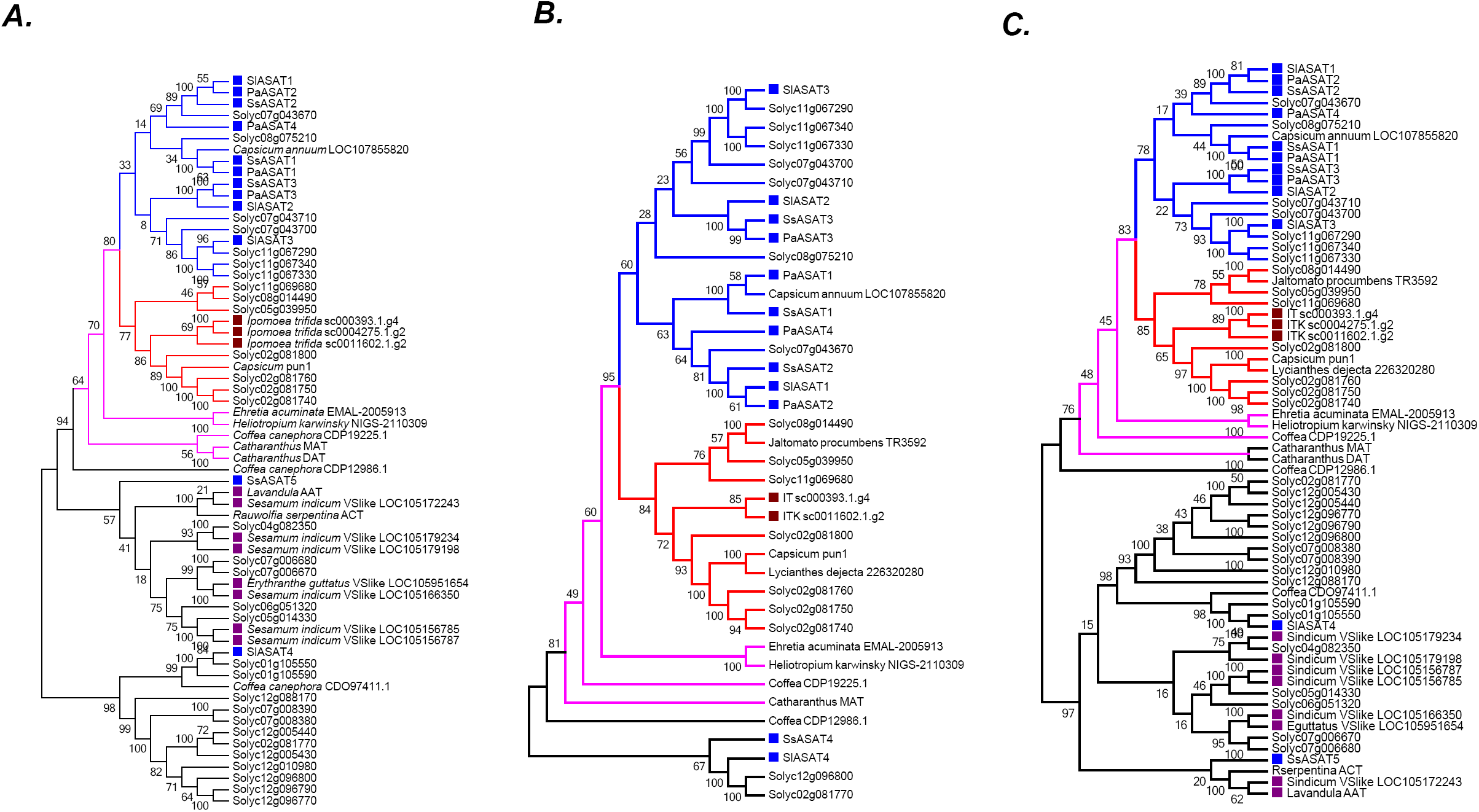
Robustness of the phylogenetic relationships. (A) A maximum likelihood tree obtained using the best model (JTT+G+I+F with 5 rate categories) as per maximum likelihood based search of multiple models. Sites with <70% coverage were deleted. (B) NJ tree using mostly the sequences in the pink+blue+red clade, with 100% site coverage and JTT model (C) NJ tree of all the sequences obtained using JTT. Only sites with 100% coverage were used for this analysis. Please see Fig. 6A for explanation of the various branch and label colors. Bootstrap support values are a result of 100 iterations.

**Figure 6-Figure Supplement 3:**
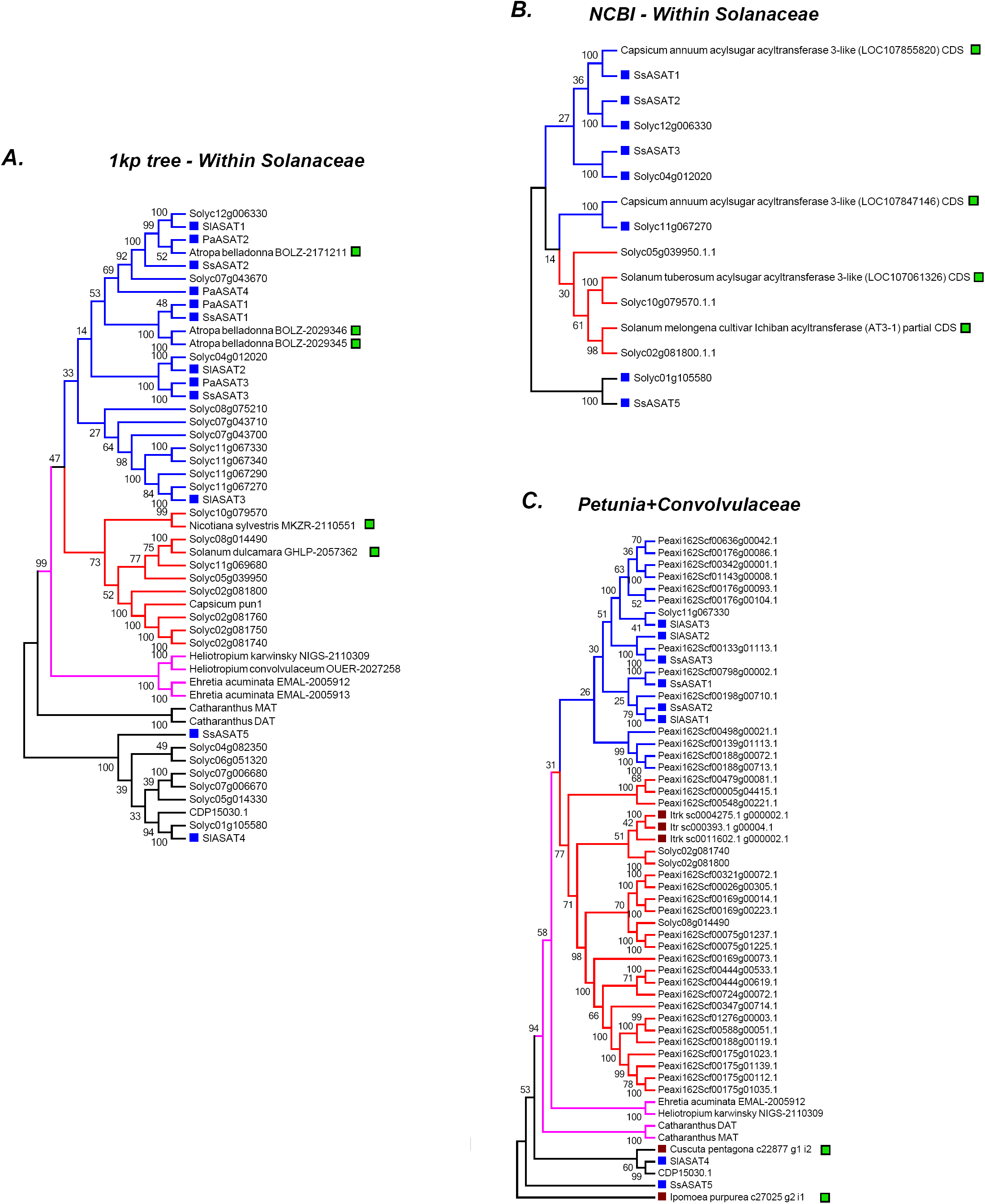
Additional related sequences do not affect our inferences regarding ASAT clade emergence. Hits obtained using various BLAST strategies (A,B) and using Petunia Clade III BAHDs (C) are shown in a phylogenetic context with ASATs and other relevant homologs. The novel hits that were not included in the final gene tree in Fig. 6 are shown by a green square. All trees were generated using NJ, using the JTT model with 100 bootstraps. All sites with <80% coverage were eliminated.

**Figure 7-Figure Supplement 1:**
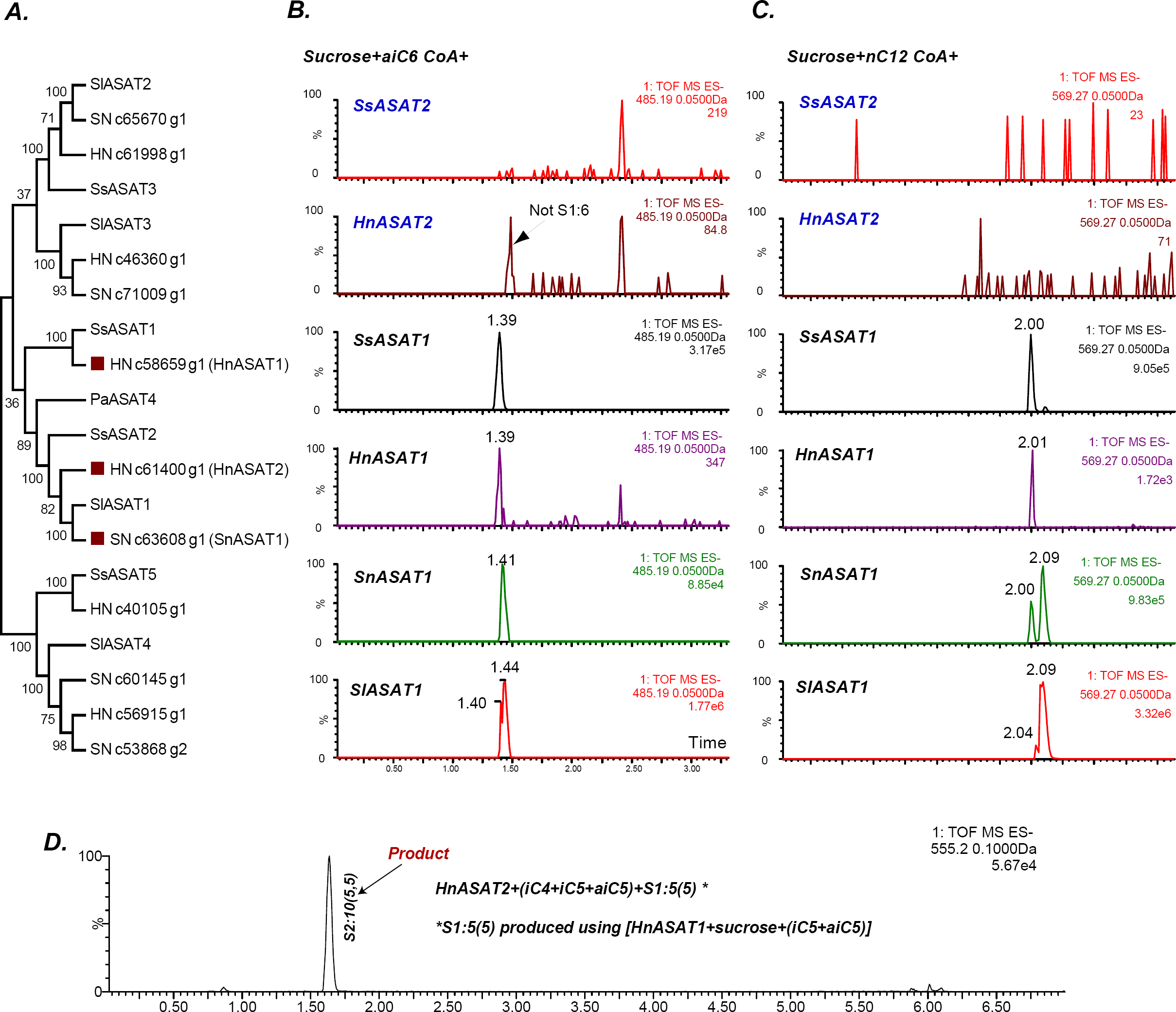
Some aASAT2 orthologs cannot catalyze sucrose to mono-acylsucrose conversion. (A) Neighbor-joining tree made using Salpiglossis and tomato ASATs and their best hits in *S. nigrum* (Sn) and *H. niger (Hn)* transcriptome database. Tree was made using default parameters in MEGA6 for alignment and phylogeny, and identifies putative aASAT2 orthologs in *S. nigrum* and *H. niger*. (B-D)Shown are LC/MS extracted ion chromatograms for S1:6(6) [B], S1:12(12) [C] and S2:10(5,5) [D]. All reactions were run on a C18 LC column. Results show that Hn, Ss, Sn and SlASAT1s can catalyze sucrose to mono-acylsucrose conversion using aiC6 and nC12 CoA. However, aASAT2 orthologs in Hn and Ss are not able to catalyze this reaction. (D) HnASAT2 is able to catalyze di-acylsucrose formation using the HnASAT1 product.

**Supplemental Table 1:** Description of the LC gradients used in this study.

| **7min-C18** | **Flow (mL/min)** | **%A** | **%B** | **Curve** |  |  |
| --- | --- | --- | --- | --- | --- | --- |
| Initial | 0.3 | 95 | 5 | Initial | A: Water+0.1% Formic acid | (-ve mode) |
| 1 | 0.3 | 40 | 60 | 6 | A: 10mM ammonium formate in water, pH 2.8 | (+ve mode) |
| 5 | 0.3 | 0 | 100 | 6 | B: Acetonitrile |  |
| 6 | 0.3 | 0 | 100 | 6 |  |  |
| 6.01 | 0.3 | 95 | 5 | 6 |  |  |
| 7 | 0.3 | 95 | 5 | 6 |  |  |

| **22min-C18** | **Flow (mL/min)** | **%A** | **%B** | **Curve** |  |  |
| --- | --- | --- | --- | --- | --- | --- |
| Initial | 0.3 | 90 | 10 | Initial | A: Water+0.1% Formic acid | (-ve mode) |
| 7 | 0.3 | 40 | 60 | 6 | A: 10mM ammonium formate in water, pH 2.8 | (+ve mode) |
| 15 | 0.3 | 0 | 100 | 6 | B: Acetonitrile |  |
| 20 | 0.3 | 0 | 100 | 6 |  |  |
| 20.01 | 0.3 | 90 | 10 | 6 |  |  |
| 22 | 0.3 | 90 | 10 | 6 |  |  |

| **110min-C18** | **Flow (mL/min)** | **%A** | **%B** | **Curve** |  |  |
| --- | --- | --- | --- | --- | --- | --- |
| Initial | 0.3 | 99 | 1 | Initial | A: Water+0.1% Formic acid | (-ve mode) |
| 1 | 0.3 | 99 | 1 | 6 | A: 10mM ammonium formate in water, pH 2.8 | (+ve mode) |
| 100 | 0.3 | 20 | 80 | 6 | B: Acetonitrile |  |
| 101 | 0.3 | 0 | 100 | 6 |  |  |
| 105 | 0.3 | 0 | 100 | 6 |  |  |
| 106 | 0.3 | 99 | 1 | 6 |  |  |
| 110 | 0.3 | 99 | 1 | 6 |  |  |

| **9min-BEH** | **Flow (mL/min)** | **%A** | **%B** | **Curve** | A: Water+0.1% Formic acid |
| --- | --- | --- | --- | --- | --- |
| Initial | 0.3 | 5 | 95 | Initial | B: Acetonitrile |
| 1 | 0.3 | 5 | 95 | 6 |  |
| 6 | 0.3 | 40 | 60 | 6 |  |
| 7 | 0.3 | 95 | 5 | 6 |  |
| 7.01 | 0.3 | 5 | 95 | 6 |  |
| 9 | 0.3 | 5 | 95 | 6 |  |

## Supplemental Methods

### Extraction and sequencing of RNA

Total RNA was extracted from 4-5 week old (*S. nigrum, S. quitoense*) or 7-8 week old plants (*H. niger,* Salpiglossis) using Qiagen RNEasy kit (Qiagen, Valencia, California) with on-column DNA digestion. In addition to trichome RNA, total RNA was extracted from shaved stems of *S. nigrum* and Salpiglossis, and shaved petioles of *S. quitoense* and *H. niger*. The quality of extracted RNA was determined using Qubit (Thermo Fisher Scientific, USA) and Bioanalyzer (Agilent Technologies, Palo Alto, California). Total RNA from all 16 samples (4 species x 2 tissues x 2 biological replicates) was sequenced using Illumina HiSeq 2500 (Illumina, USA) in two lanes (8X multiplexing per lane). Libraries were prepared using the Illumina TruSeq Stranded mRNA Library preparation kit LT, sequencing carried out using Rapid SBS Reagents in a 2x100bp paired end format, base calling done by Illumina Real Time Analysis (RTA) v1.18.61 and the output of RTA was demultiplexed and converted to FastQ format by Illumina Bcl2fastq v1.8.4.

### Confirmation of *Salpiglossis sinuata*

We confirmed the plant to be Salpiglossis using the chloroplast *ndhF* and *trnLF* marker based phylogenies (Figure 2-Figure Supplement 1A,B). Specifically, we amplified these regions using locus-specific primers, sequenced the amplicons and assembled the contigs using Muscle (Edgar, 2004). A neighbor joining tree including *ndhF* and *trnLF* sequences from NCBI Genbank was used to confirm the identity of the plant under investigation.

### Prediction of protein sequences and orthologous group assignments

The protein sequences corresponding to the longest isoform of all expressed transcripts (read count>10 in at least one dataset in a given species) were obtained using TransDecoder (Haas et al., 2013) and GeneWise v2.1.20c (Birney et al., 2004). Only the protein sequences of transcripts with ≥10 reads as defined by RSEM were used for constructing orthologous groups using OrthoMCL v5 (Li et al., 2003). We defined orthologous relationships between *S. lycopersicum*, *S. nigrum*, *S. quitoense*, *H. niger*, *N. benthamiana*, Salpiglossis and *Coffea canephora* (outgroup) using an inflation index of 1.5.

### Gene Ontology enrichment analyses

We transferred the tomato gene ontology assignments to the homologs from other species in the same orthologous group. GO enrichment analysis was performed using a custom R script, and enriched categories were obtained using Fisher Exact Test and correction for multiple testing based on Q-value (Storey, 2002).

#### qRT-PCR

Primers to specific regions of the targeted transcript were designed with amplicons between 100- 200 bp using Primer3 (Untergasser et al., 2012). The regions selected for amplification did not overlap with the region targeted in the VIGS analysis. Primer sets for Salpiglossis orthologs of PDS and elongation factor alpha (EF1a) were used as controls. We used 1 μg of total RNA from a single VIGS plant with an acylsugar phenotype to generate cDNA using the Thermo Fisher Superscript II RT kit. An initial amplification and visualization on a 1% agarose gel was performed to ensure that the primers yielded an amplicon with the predicted size and did not show visible levels of primer dimers. We first tested multiple primer sets per gene and selected primers within 85-115% efficiency range using a dilution series of cDNA from uninoculated plants. These primers were used for the final qRT-PCR reaction. The Ct values for the transcripts (on 1x template) were measured in triplicate, which were averaged for the analysis. Both uninoculated and empty vector controls were measured with all primer sets for ΔΔCt calculations.

### Similarity searches using BLAST

Similarity searches against the 1kp and NCBI nr databases were performed using TBLASTN, using SlASAT1, SlASAT3, SsASAT1 and SsASAT2 protein sequences as queries. For the 1kp database, TBLASTN was performed against all Asterid sequences, and the best non-Solanaceae sequences were analyzed using phylogenetic tree reconstruction (see below; Figure 6-Source Data 1). The BLAST at NCBI was performed against several specific groups of species, namely (i) orders Gentianales+Boraginales+Lamiales+Solanales-(family Solanaceae), (ii) Only Convolvulaceae, (iii) Solanaceae-(sub-family Solanoideae), (iv) Solanoideae-(genera Solanum+Capsicum), (v) Solanoideae-(genus Solanum), (vi) Solanum-(section Lycopersicon+species tuberosum). The top 10 hits from each search were manually curated and analyzed in a phylogenetic context with SlASATs and SsASATs (Figure 6-Figure Supplement 2,3). We used a similar approach to search the annotated peptide sequences in *C. canephora* (Denoeud et al., 2014) and *Ipomoea trifida* (Hirakawa et al., 2015) sequence databases. Sequences that provided additional insights into the evolution of the ASAT clade were integrated into the final gene tree shown in Figure 6A.

### Synteny analysis

Synteny between the Petunia, Capsicum and tomato genomes was determined as previously described using MCScanX (Moghe et al., 2014; Wang et al., 2012). Regions corresponding to PaASAT1 and its best matching homologs were used to make Figure 7B.

### Estimating the number of acylsugars

This and previous studies provide evidence for a number of acyl chains esterified to sucrose, classified into short (C2, C3, C4, iC5, aiC5, iC6, aiC6, C8), long (nC10, iC10, nC11, nC12) and non-aliphatic (malonyl). We only focused on aliphatic chains for this estimate. We typically see one long chain and 3-4 short chains on the sugar molecule across different Solanaceae species. In our calculation, we assumed that each acyl chain has the same probability of being incorporated into the acylsugar, with the core being a sucrose core. Under these assumptions, there are 8 acyl chains that could be incorporated at three positions on a tetra-acylsugar and 12 acyl chains on the fourth position. This gives a theoretical estimate of 8*8*8*12=6144 acylsucroses that could be produced with the above acyl chains.

